# Fast and accurate modeling and design of antibody-antigen complex using tFold

**DOI:** 10.1101/2024.02.05.578892

**Authors:** Fandi Wu, Yu Zhao, Jiaxiang Wu, Biaobin Jiang, Bing He, Longkai Huang, Chenchen Qin, Fan Yang, Ningqiao Huang, Yang Xiao, Rubo Wang, Huaxian Jia, Yu Rong, Yuyi Liu, Houtim Lai, Tingyang Xu, Wei Liu, Peilin Zhao, Jianhua Yao

## Abstract

Accurate prediction of antibody-antigen complex structures holds significant potential for advancing biomedical research and the design of therapeutic antibodies. Currently, structure prediction for protein monomers has achieved considerable success, and promising progress has been made in extending this achievement to the prediction of protein complexes. However, despite these advancements, fast and accurate prediction of antibody-antigen complex structures remains a challenging and unresolved issue. Existing end-to-end prediction methods, which rely on homology and templates, exhibit sub-optimal accuracy due to the absence of co-evolutionary constraints. Meanwhile, conventional docking-based methods face difficulties in identifying the contact interface between the antigen and antibody and require known structures of individual components as inputs. In this study, we present a fully end-to-end approach for three-dimensional (3D) atomic-level structure predictions of antibodies and antibody-antigen complexes, referred to as tFold-Ab and tFold-Ag, respectively. tFold leverages a large protein language model to extract both intra-chain and inter-chain residue-residue contact information, as well as evolutionary relationships, avoiding the time-consuming multiple sequence alignment (MSA) search. Combined with specially designed modules such as the AI-driven flexible docking module, it achieves superior performance and significantly enhanced speed in predicting both antibody (1.6% RMSD reduction in the CDR-H3 region, thousand times faster) and antibody-antigen complex structures (37% increase in DockQ score, over 10 times faster), compared to AlphaFold-Multimer. Given the performance and speed advantages, we further extend the capability of tFold for structure-based virtual screening of binding antibodies, as well as de novo co-design of both structure and sequence for therapeutic antibodies. The experiment results demonstrate the potential of tFold as a high-throughput tool to enhance processes involved in these tasks. To facilitate public access, we release code and offer a web service for antibody and antigen-antibody complex structure prediction, which is available at https://drug.ai.tencent.com/en.

## 1 Introduction

Antibodies, generated by clonal B cells, serve a pivotal function in the human adaptive immune system by specifically recognizing and responding to foreign molecules or antigens [1]. They contribute to immunity through three primary strategies: neutralization, which entails binding to pathogens and inhibiting their entry or damage to host cells; opsonization, which involves coating the pathogen to promote the pathogen elimination by macrophages and other immune cells; the initiation of pathogen destruction by activating supplementary immune responses, such as the complement pathway. The high specificity and affinity of antibodies in antigen recognition also render them highly promising therapeutic agents, garnering extensive research attention in recent years [2, 3]. Modelling the antibody-antigen complex structures can pave the foundation for elucidating the structural principles that regulate antibody–antigen interactions, which is therefore essential for enhancing our knowledge of the immune system’s properties in terms of molecular structures and advancing our capacity to design efficient biological drugs [1–3].

The high-throughput B-cell sequencing has generated vast amount of data needed to investigate complex mechanisms underlying the adaptive immune response, thus paving the way for data-driven investigations of antibodies [4]. Nevertheless, the high-throughput production of precise antibody-antigen complex structures remains a major challenge in both structural and computational biology [5, 6]. On one hand, experimental techniques such as X-ray crystallography, cryo-electron microscopy, nuclear magnetic resonance (NMR), and mutagenesis analysis for investigating antibody-antigen complex structures are usually expensive, time-consuming, and exhibit a low success rate, hindering their potential as high-throughput approaches [7]. On the other hand, computational methods like molecular docking for modelling antibody-antigen structures often result in high false-positive rates and seldom yield unique solutions, necessitating known structures of individual components as inputs [8–13]. Moreover, docking-based methods struggle to identify epitopes when protein binding sites are ambiguous and substantial conformational changes occur during binding, a common issue in antibody-antigen interactions. Recently, advanced deep learning-based methods, such as AlphaFold-multimer [5], have significantly advanced structure prediction for multi-chain protein-protein complexes. However, their performance on antibody-antigen complexes is sub-optimal, primarily because antibodies and antigens often originate from different species, precluding the acquisition of effective co-evolutionary signals [14]. Additionally, the substantial time consumption associated with these methods hinders their potential to serve as high-throughput tools.

In this study, we present a fast and accurate method for predicting the structure of the antibody and antibody-antigen complex, which leverages advanced deep-learning technologies to support research in immunology and the development of therapeutic antibodies. The proposed method equipped with a pre-trained large protein sequence model and transformer-based structure prediction modules, termed tFold, enables fast end-to-end atomic-resolution antibody (hereafter denoted as tFold-Ab) and antibody-antigen complex (hereafter denoted as tFold-Ag) structure prediction directly from the sequence, which has the potential to be a high-throughput tool. Moreover, the high speed and accuracy of tFold’s structure prediction capabilities offer the potential to improve various processes involved in the design and development of therapeutic antibodies. This study exemplifies the capability of using tFold for structure-based virtual screening of binding antibodies, as well as the advantage of using tFold for de novo co-design of structure and sequence for therapeutic antibodies, as two examples to illustrate its extensive potential applications.

## 2 Results

In overview, this section firstly presents the performance of tFold for antibody and nanobody structure prediction (tFold-Ab) in Section 2.1. Subsequently, it introduces the performance of tFold for antibody-antigen and nanobody-antigen complex structure prediction (tFold-Ag) in Section 2.2. Based on the high performance and computational speed of tFold-Ag in structure prediction, we further highlight two applications of tFold-Ag, namely, structure-based virtual screening of binding antibodies and de novo co-design of structure and sequence for therapeutic antibodies, in Section 2.3 and Section 2.4, respectively. To thoroughly evaluate the performance of tFold in the aforementioned tasks, we have curated a total of ten datasets from three data sources (Table B1).

### 2.1 Fast and accurate prediction of antibody structures

The accurate prediction of antibody structures is crucial for examining their function, which serves as a prerequisite stage for tFold in generating predictions of antibody-antigen complex structures. Besides the ability for structure prediction of antibody-antigen complex, tFold is also capable of independently generating antibody structure predictions. The component of tFold responsible for predicting antibody structures is referred to as tFold-Ab, which is specifically designed as a high-throughput, end-to-end approach that directly produces high-resolution, atomic-level three-dimensional (3D) structures of antibodies, accompanied by a per-residue confidence score, utilizing their amino acid sequences as input.

The tFold-Ab consists of four main modules including a pre-trained large protein language model: ESM-PPI, a feature updating module: Evoformer-Single, a structure module and a recycling module (Fig. 1a). The ESM-PPI module works to extract both the intra-chain and inter-chain information of the protein complex to generate features for the down-streaming structure prediction task. We develop ESM-PPI by extending the current ESM-2 model [15] through further pre-training using both monomers and, more importantly, multimers curated from four large databases, including UniRef50 [16], Protein Data Bank (PDB), Protein-Protein Interaction (PPI), and the Antibody database (refer to Appendix C.3 for details). This enhancement enables the model to extract inter-chain information effectively. The Evoformer-Single module updates and refines the input protein features from the language model, iteratively if using the recycling mechanism, to produce the informative sequence and pairwise representations. The structure module, which performs SE(3)-equivariant updates using invariant point attention [17], then maps the obtained representation to predicted atomic-level 3D structures. Finally, the recycling module allows tFold-ab to reuse the features and structure predictions of the previous iteration to improve the quality of the final structure prediction. To conduct a comprehensive evaluation of tFold-Ab’s performance, we have curated a test set, held out from the training and validation data, based on a cutoff date of 01 July 2022. It comprises two non-redundant benchmark subsets, namely SAbDab-22H2-Ab and SAbDab-22H2-Nano, which include 170 paired antibodies and 73 single-chain nanobodies respectively. We utilize the widely adopted temporally separated approach [17, 18] during data preparation to guarantee fair comparisons between tFold-Ab and other existing methods [5, 15, 18–24].

**Fig. 1:**
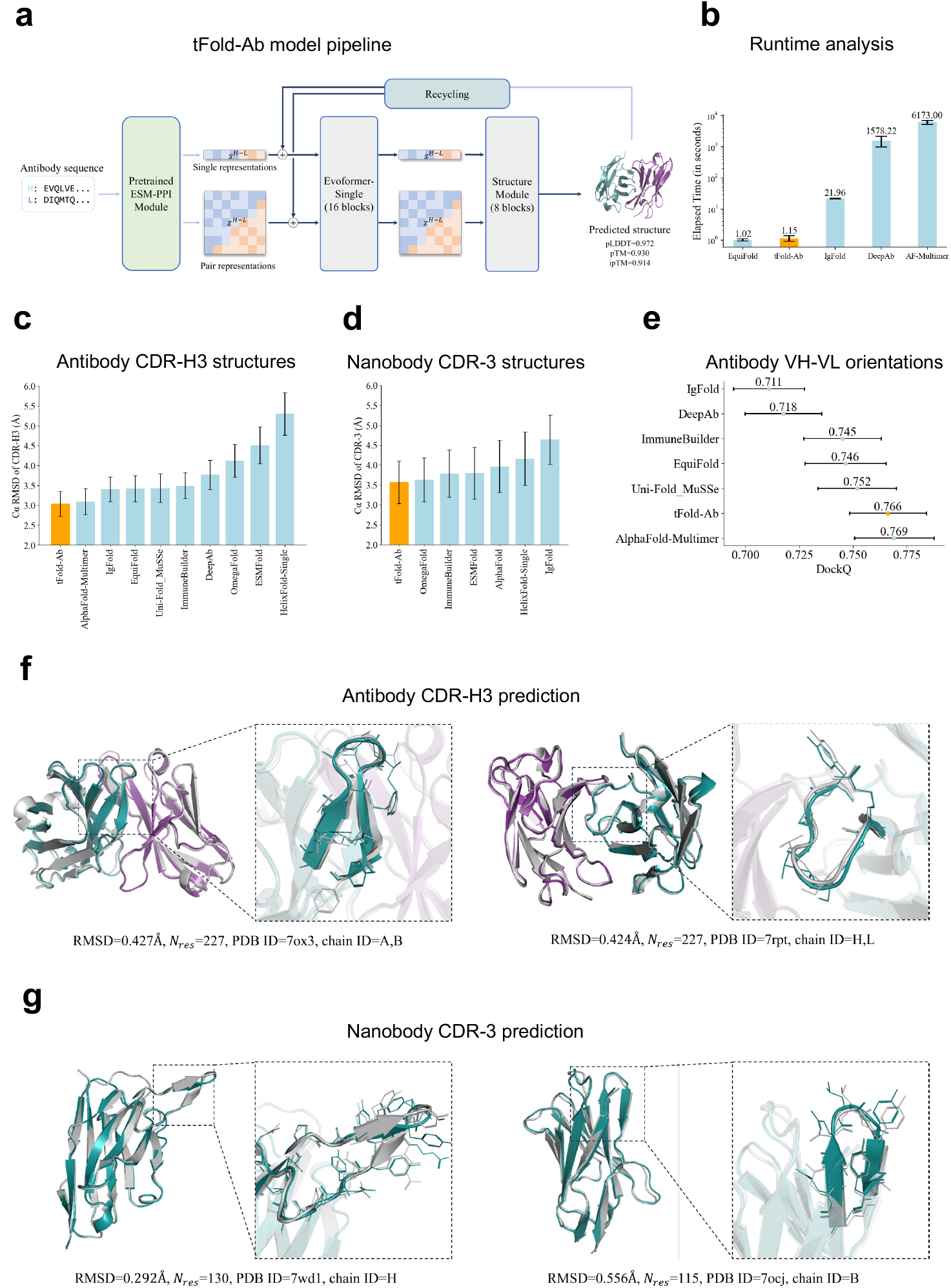
tFold-Ab method and result. (a) Overview of the tFold-Ab architecture, with arrows indicating the direction of information flow. Gradient backpropagation is only enabled for dark arrows, but not for light arrows. (b) Runtime analysis for tFold-Ab on 170 antibodies in SAbDab-22H2-Ab. Comparison to DeepAb, IgFold, AlphaFold-Multimer and EquiFold. All runtime are measured on a single NVIDIA A100 GPU with 21 CPU cores. (c-d) Performance comparison of tFold-Ab with other antibody structure prediction method on SAbDab-22H2-Ab and SAbDab-22H2-Nano test sets, evaluated by backbone RMSD in CDR-H3 for antibody and CDR-3 for nanobody, represented as mean data with a 95% confidence interval. (e) DockQ evaluation performance on the SAbDab-22H2-Ab. (f) Comparison of our predicted structures for antibody targets (PDB 7ox3 and 7rpt, blue for the heavy chain, purple for light chain) with their respective experimental structures (gray). The accurate prediction of the CDR-H3 region by tFold-Ab for both side and main chains is particularly noteworthy. (g) Comparison of our predicted structures for nanobody targets (PDB 7wd1 and 7ocj, blue for heavy chain) with their respective experimental structures (gray). The accurate prediction of the CDR-H3 region by tFold-Ab for both side and main chains is particularly noteworthy.

As illustrated in Fig. 1c and supplementary Table B2, tFold-Ab results in high performance in antibody structure prediction with average root-mean-squared-deviation (RMSD) values of 0.61 Å and 0.57 Å for the framework regions(FR) in the heavy chains and light chains, respectively. In the more complex task of predicting the complementarity-determining regions (CDRs), tFold-Ab achieves average RMSD of 0.92, 0.84, and 3.04 Å in the CDR-1, CDR-2, and CDR-3 regions of the heavy chains (denoted as H1, H2, and H3 regions), along with RMSD of 0.87, 0.73, and 1.12 Å in the corresponding CDRs of the light chains (denoted as L1, L2, and L3 regions). All RMSD scores are calculated over the backbone heavy atoms, following the alignment of the respective framework residues. We compared tFold-Ab with the currently existing general protein structure prediction methods including AlphaFold-Multimer [5], EquiFold [20], Uni-Fold MuSSe [23], ESMFold [15], OmegaFold [24], HelixFold-Single [21] as well as antibody-specific methods including IgFold [18], DeepAb [19], ImmuneBuilder [22]. It should be noted that ESMFold [15], OmegaFold [24] and HelixFold-Single [21] are single-sequence (monomer) structure prediction methods. The heavy and light chains are separately predicted when compared with other methods that directly predicted the paired chain complexes. In the framework regions, all the examined methods consistently exhibited the highest performance compared to other regions. The CDR-H3 and CDR-L3, however, are the most challenging components for prediction, due to the significant sequence and structural diversities in these regions. Overall, tFold-Ab demonstrates superior performance compared to both general protein structure prediction methods and antibody-specific methods across all framework regions and CDRs of both chains (Fig. 1c and supplementary Table B2).

The anticipated orientation between the heavy and light chains plays a crucial role in determining the overall binding surface. To assess the accuracy of the interchain orientation, we measured the orientational coordinate distance (OCD)[25]. Our findings indicate that, in general, tFold-Ab achieves significantly better OCD values compared to the other methods evaluated, with a substantial margin. Specifically, tFold-Ab yields an OCD of 4.74, while the competing methods range from 5.20 to 5.88. To further evaluate the accuracy of the predicted antibody complex structure of paired chains, we include four metrics including the DockQ, fraction of native contacts (Fnat), ligand root-mean-square deviation (LRMS), and the interface root-mean-square deviation (iRMS). tFold-Ab and AlphaFold-Multimer achieve the best performance among the evaluated methods, demonstrating comparable results with average DockQ scores of 0.766 and 0.769, respectively. In particular, tFold-Ab achieves the lowest average LRMS (1.889 Å vs a range of 2.077 to 2.332 Å) and iRMS (1.339 Å vs a range of 1.416 to 1.622 Å) scores, indicating its exceptional accuracy in predicting both ligand and interface positions, while AlphaFold-Multimer obtains the best Fnat score of 0.786 among all evaluated methods (Supplementary Table B3).

Nanobodies, as a promising format for therapeutic development, have gained considerable attention currently. Different from the paired antibodies, nanobodies lack the second immunoglobulin chain [18, 26]. This characteristic, coupled with the increased length of the nanobody CDR-3 loop, results in a wide range of CDR-3 loop conformations that are not typically observed in paired antibodies [18, 26]. The performance of tFold-Ab and other existing general and antibody-specific methods [5, 15, 18, 19, 21, 22, 24] are compared in Fig. 1d and supplementary Table B4, where we omit EquiFold [20] and Uni-Fold MuSSe [23] from the comparison since they are only suitable for paired antibody structures. All evaluated methods exhibit their highest accuracy within the framework regions, underscoring the predictability of these conserved areas. However, the CDR-3 loop region pose the greatest challenge, reflecting the complexity and variability of these loops. Generally, tFold-Ab results in superior performance in most regions including the framework region (RMSD=0.67 Å vs RMSDs ranging from 0.71 to 0.93 Å), CDR-2 region (RMSD=1.20 Å vs RMSDs ranging from 1.20 to 1.51 Å), and CDR-3 region (RMSD=3.57 Å vs RMSDs ranging from the 3.63 to 9.03 Å).

Fig. 1f and 1g present example predicted structures of antibodies and nanobodies, offering an intuitive visualization of the prediction results. It is observed that tFold-Ab is capable of providing highly accurate predictions for CDR-H3 and CDR-3, which are the most challenging regions in the structure prediction of antibodies and nanobodies respectively. In addition to its superior prediction accuracy, tFold-Ab also achieves state-of-the-art computational speed in antibody structure prediction. This efficiency is primarily obtained through the employment of a pre-trained protein language model (ESM-PPI) rather than traditional MSA-based approaches for extracting co-evolutionary information. Moreover, tFold-Ab employs a specially optimized Evoformer-Single module and structure module to predict both backbone and side-chain conformations in an end-to-end fashion. In contrast, existing methods such as DeepAb [19] and IgFold [18] rely on Rosetta-based energy minimization for side-chain structure prediction, which is time-consuming. We compare the inference time of tFold-Ab and existing methods in Fig. 1b. As observed, tFold-Ab is 5,367 times faster than AlphaFold-Multimer [5], owing to the use of ESM-PPI and the Evoformer-Single module. We also compared tFold-Ab with an AlphaFold-Multimer variant without employing the MSA and template search procedures, the superior speed of tFold-Ab suggests the computational efficiency of the Evoformer-Single module compared to the conventional Evoformer in the AlphaFold-Multimer. EquiFold [20], which employs geometrical structure representations instead of representations extracted from a pre-trained protein language model, achieves a marginally faster computational speed than tFold-Ab (1.02 vs 1.15 second). However, its performance is significantly inferior (P-value=0.049) compared to tFold-Ab. (Inference times are measured on a single NVIDIA A100 GPU with 21 CPU cores for all methods, and more details can be found in Appendix section C.4.9).

### 2.2 Fast and accurate prediction of antibody-antigen complex structures

The structure of the constructed antibody-antigen complex prediction model, termed tFold-Ag for ease of differentiation from tFold-Ab, is depicted in Fig. 2a. It mainly consists of three modules, i.e. the antibody feature generation module, the antigen feature generation module and the AI-driven flexible docking module. The antibody feature generation module reuses the previously trained tFold-Ab to extract the sequence and pair representations as well as the initial structures (atom coordinates) of the antibodies. During the training of tFold-Ag, the antibody feature generation module remains fixed to facilitate the convergence and optimization of the entire model. The antigen feature generation module employs the pre-trained AlphaFold2 [17] to extract the sequence and pair representations together with initial structures (atom coordinates) of the antigens. This design considers the extensive diversity of antigens, which may originate from a variety of sources, including bacteria, viruses, cancer cells, and so on. As AlphaFold2 was pre-trained on a considerably broad range of proteins, this strategy enables tFold-Ag to exhibit enhanced generalizability to diverse antigens. The AI-driven flexible docking module comprises two main components: a specially designed feature fusion module and a complex structure prediction module. Upon obtaining the sequence and pair representations as well as the initial coordinates of the antibody and antigen from the aforementioned feature generation modules, the innovative feature fusion module is subsequently employed to effectively integrate this information, thereby generating the initial sequence and pair representations of the antibody-antigen complex. Following this, the proposed complex structure prediction module, which includes an Evoformer-Single stack with 32 blocks and a structure module with 8 blocks, updates the representations (via the Evoformer-Single blocks) and subsequently maps these representations to the predicted complex structure and provides the predicted confidences (via the structure blocks). In the AI-driven flexible docking module, tFold-Ag not only calculates the conformation of the antibody-antigen complex but also updates the structures of the antibody and antigen themselves, which allows tFold-Ag to refine the initially extracted structures (refer to the method section and Appendix C.5 for details). For instance, the antibody structure prediction accuracy of the most variable CDR-H3 region is further improved in tFold-Ag (when the prediction of the epitope is accurate or when additional epitope information is available), with the RMSD decreasing from 3.21Å to 3.07Å compared to the predicted conformation in tFold-Ab. It is worth noting that, in addition to the complex structure prediction-related modules mentioned above, there is an extra sequence recovery module between the Evoformer-Single stack and structure module in the complex structure prediction module, which is employed for novel antibody design (Details are provided in sections 2.4 and 4.5). During the training phase, we simultaneously train the shared and specific modules for complex structure prediction and antibody design, which makes tFold-Ag a multi-task model. The optimization of the model for each task can be considered as a regularization task that benefits the training of its counterpart task (structure prediction and antibody design).

**Fig. 2:**
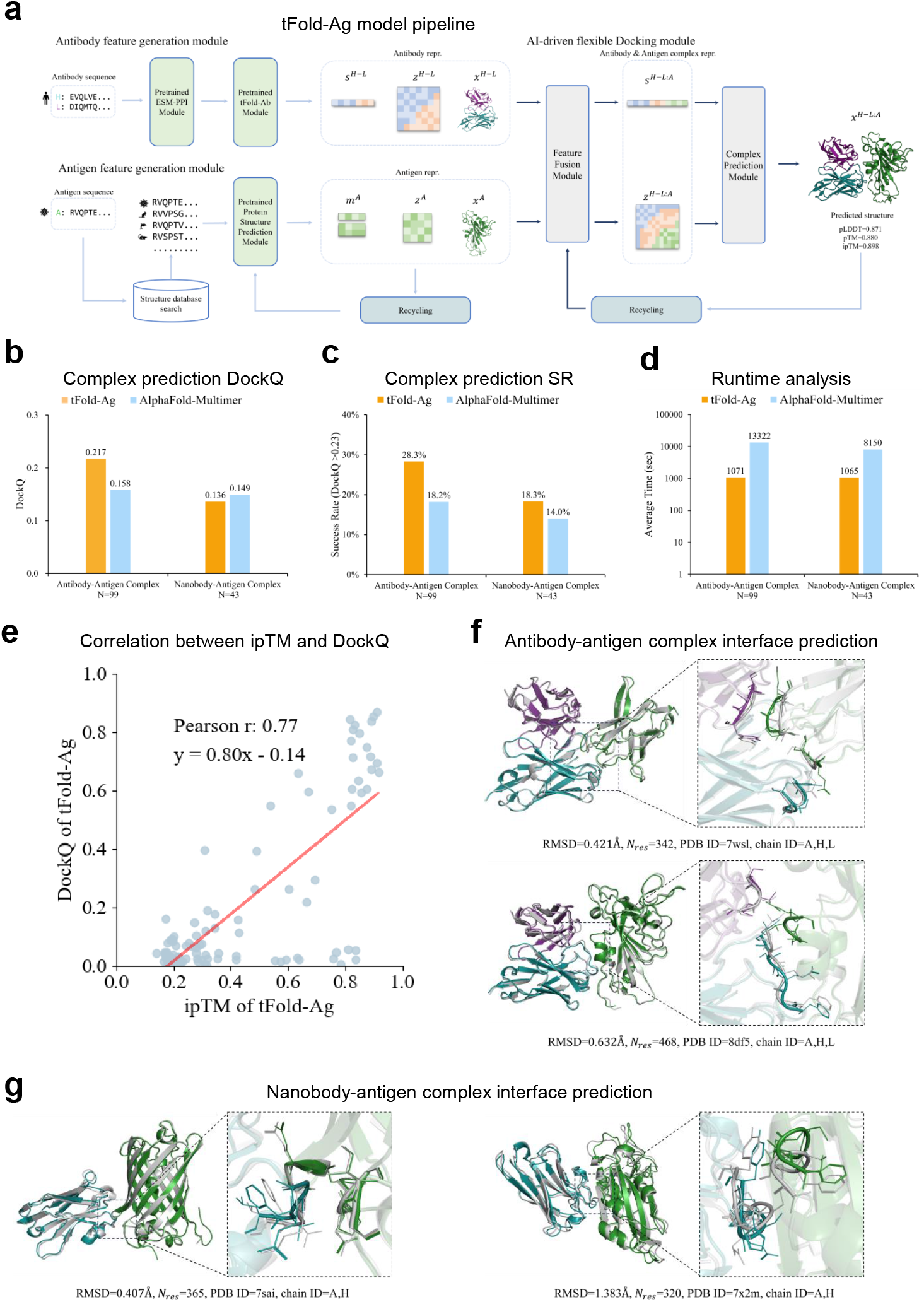
tFold-Ag method and result. (a) Overview of the tFold-Ag architecture. Arrows show the information flow direction. The dark arrows indicate gradient is used while the light arrows gradient is not used. Here, repr. denotes representation. (b-d) The antibody-antigen interaction accuracy, success rate and running time comparison of tFold-Ag and AlphaFold-Multimer on SAbDab-22H2-AbAg and SAbDab-22H2-NanoAg test set. (e) Correlation between ipTM and DockQ. Least-squares linear fit DockQ = 0.80 × ipTM - 0.14 (Pearson’s r = 0.77, P-value = 1.2 *×* 10^*−*20^). n = 99 protein chains in SAbDab-22H2-Ab. (f) Comparison of our predicted prediction for antibody-antigen complex targets 7wsl and 8df5 (blue for heavy chain, purple for light chain and green for antigen chain) with their respective experimental structure (gray). The interface of antigen and antibody is well predicted. (g) Comparison of our predicted prediction for nanobody-antigen complex targets 7sai and 7×2m (blue for heavy chain and green for antigen chain) with their respective experimental structure (gray). The interface of nanobody and antibody is well predicted.

Similar to the data preparation process in antibody structure prediction, we curate a hold-out test set to evaluate the performance of structure prediction methods on antibody-antigen complexes from the SAbDab database [27], employing a temporally separated approach [17, 18]. To ensure a fair comparison with existing methods such as AlphaFold-Multimer [5], we make sure that none of the evaluated methods have been exposed to the structures included in the test set by setting the cutoff date to July 1, 2022. The finalized test set comprises two non-redundant benchmark subsets, namely SAbDab-22H2-AbAg and SAbDab-22H2-NanoAg, which include 99 antibody-antigen complexes and 43 nanobody-antigen complexes, respectively.

We compare tFold-Ag with currently available end-to-end methods, including AlphaFold-Multimer [5], Uni-Fold MuSSe [23], and RoseTTAFold2 [28], as well as docking-based methods such as conventional docking methods like ZDock [9], Clus-Pro [8], HDock [10], and deep-neural-network-involved docking methods such as EquiDock [11], dyMEAN [12], and ColabDock [13]. tFold-Ag and other end-to-end methods (sequences-to-complex structure) can directly predict the complex structure using input sequences, whereas docking-based methods (structures-to-complex structure) necessitate the structures of individual components to predict the final structure of the complex. To make a fair comparison, we utilized the currently available best predictor to generate the structures of antibodies (AlphaFold-Multimer [5]), nanobodies, and antigens (AlphaFold2 [17]) as initial inputs for the docking-based methods. As illustrated in Fig. 2b, Fig. 2c and supplementary Table B5, the proposed tFold-Ag outperforms existing end-to-end and docking-based methods in predicting the structure of the antibody-antigen complex, achieving a DockQ score of 0.217 and a TM-score of 0.708. The most recently updated AlphaFold-Multimer (v2.3.2) ranks second, yielding a DockQ score of 0.158 and a TM-score of 0.665. Compared to the AlphaFold-Multimer, tFold-Ag demonstrates a more robust performance achieving a higher successful rate (SR) of 0.283 versus 0.182 (SR denotes the proportion of predicted structures that meet the acceptable criterion (DockQ *>* 0.23)). A similar conclusion can be drawn in the structure prediction of the nanobody-antigen complex. tFold-Ag and AlphaFold-Multimer perform best with DockQ scores of 0.136 and 0.149 as well as TM-scores of 0.678 and 0.692, respectively (Fig. 2b, Fig. 2c and supplementary Table B6). Fewer experimental structures for nanobody-antigen complexes increase reliance on general protein prediction models, possibly explaining tFold-Ag’s slightly lower performance compared to AlphaFold-Multimer. In terms of performance robustness, tFold-Ag achieves 32% SR score enhancement compared to the AlphaFold-Multimer. Comparing the overall performance of involved methods employed in the structure prediction of nanobody-antigen and antibody-antigen complexes, it was observed that all methods demonstrated inferior performance for nanobody-antigen complexes. This can be primarily attributed to the previously mentioned factor that there exist fewer experimental nanobody-antigen structures available for training the models. Fig. 2f and 2g illustrate examples of predicted antibody-antigen and nanobody-antigen complexes, along with the corresponding experimental structures, demonstrating that tFold-Ag can provide high-quality predictions of the interaction interfaces between antibodies or nanobodies and antigens.

In structural biology practice, aside from experimental direct structure determination methods such as cryo-electron microscopy, there are relatively cost-effective and more accessible experimental approaches capable of providing interaction interface information. This information can be employed as structural constraints in the prediction of protein complex structures. For example, chemical cross-linking (XL) technology provides the distance between two residues connected by fixed-length reagents, which can be transformed into contacts between the antibody and antigen. Concurrently, Deep Mutation Scanning (DMS) can offer protein-protein interaction (PPI) information, including antigen epitopes and antibody paratopes. Although these experimental constraints are sparse and insufficient to fully determine the protein complex structure, they can offer crucial insights into the interaction interface of the components. In tFold-Ag, we propose to integrate inter-chain feature for better protein complex structure prediction via utilizing a specially designed inter-chain feature embedding module, which synthesizes the interface information into the sequence and pair representations obtained after the feature fusion module of tFold-Ag (Referring to section 4.4.2 and Appendix C.5.3 for more details). We employ the terms tFold-Ag-ppi and tFold-Ag-contact to represent tFold-Ag utilizing PPI) information (antigen epitope sites or both antigen epitope and antibody paratope sites), as well as more detailed contact information (contact map between epitope and paratope sites), respectively. Our findings suggest that tFold-Ag can significantly benefit from incorporating additional structural constraints, and the more detailed the interaction interface information is, the greater the enhancement in performance (Supplementary Table B5 and Table B6). For example, when given information on both antigen epitope and antibody paratope sites, tFold-Ag achieves a DockQ score of 0.416 in the prediction of antibody-antigen structures and a DockQ score of 0.316 in the prediction of nanobody-antigen structures (supplementary Table B5 and Table B6).

Furthermore, we have examined the correlation between the prediction confidence score and the structure prediction accuracy of tFold-Ag, using the interface pTM (ipTM) and DockQ as metrics, respectively. We observed considerable positive correlations between ipTM and DockQ on antibody-antigen data (Pearson correlation coefficient r=0.77), and relatively strong positive correlations on nanobody-antigen data (r=0.49). The primary factor leading to the correlation difference between antibody-antigen and nanobody-antigen is assumed to be the lesser amount of data available for training the model in the case of nanobodies compared to antibodies. These positive correlations consistently exist when incorporating interface information into tFold-Ag, such as tFold-Ag-ppi using epitope information or both epitope and paratope information (Fig. 2e, refer to Appendix section C.5.8 for details).

As representative methods of without and with using the MSA of antibodies, it is worth noting that employing both tFold-Ag and AlphaFold-Multimer cooperatively in an ensemble manner can lead to improved antibody-antigen and nanobody-antigen complex structure predictions. This observation is derived from a point-to-point performance comparison analysis between tFold-Ag and AlphaFold-Multimer on the involved test sets, where we discovered that these two methods produce highly complementary results. Specifically, tFold-Ag and AlphaFold-Multimer exhibited superior performance in different cases and a higher prediction confidence score obtained from these two models mostly indicates the better obtained prediction structure (additional information can be found in Section C.5.9). Consequently, we design an ensemble strategy utilizing a selection mechanism referring to the prediction confidence scores obtained from the two involved models, i.e. the prediction with a higher prediction confidence score is selected as the ensembled prediction. With using this strategy, the DockQ and Success Rate (SR) can be enhanced from 0.217 and 0.283 (when using tFold-Ag independently) to 0.288 and 0.404 for antibody-antigen complexes (Table B5 and Table C19). In the case of nanobody-antigen complexes, the DockQ can reach 0.218 (from 0.136), and the SR can be enhanced to 0.279 (from 0.186) when employing the ensemble strategy.

In addition to providing state-of-the-art accuracy in predicting antibody-antigen and nanobody-antigen complex structures, another advantage of tFold-Ag is its optimized computational speed. tFold-Ag is over 10 times faster than AlphaFold-Multimer in the structure prediction of protein complexes with different overall sequence lengths (Supplementary Fig. C9). The enhancement in computational speed primarily results from two factors. The first is our design strategy that employs the pre-trained large protein language model ESM-PPI to extract both intra-chain and inter-chain residue-residue contact information, as well as evolutionary relationships, instead of relying on the time-consuming multiple sequence alignment (MSA) strategy. The second factor is our optimization and simplification of the remaining model architecture, such as the AI-driven flexible docking component. The speed enhancement of tFold-Ag, compared to AlphaFold-Multimer, is less pronounced than that of tFold-Ab. This is due to the retention of the MSA in the antigen feature generation module of tFold-Ag to ensure better accuracy and generalizability. Therefore, the speed advantage of tFold-Ag will become increasingly significant when predicting the complex structures of a set of antibodies and a specific target antigen in a high throughput manner. Considering that the antigen feature generation module is the most time-consuming component in tFold-Ag, owing to the employment of MSA, the average time required for predicting each antibody-antigen complex will decrease correspondingly with an increase in the number of antibodies screened against the particular antigen.

### 2.3 Structure-based virtual screening of binding antibodies

In the early discovery phase of therapeutic antibody development, the primary task is to select high-affinity binding antibodies, also known as binders, from candidate antibodies obtained through animal immunization or phage display technologies. Traditional wet-lab techniques such as ELISA, FACS, or more recently emerged optofluidic systems, can filter out part of non-binding antibodies, but these methods are not as accurate as high-precision macromolecular interaction measurements like Surface Plasmon Resonance (SPR) or Bio-Layer Interferometry (BLI) [29]. However subjecting thousands of antibodies to expression and purification followed by SPR or BLI, would entail considerable time and experimental costs. To address this, we have explored the potential of the tFold-Ag model to predict the structure of antibody-antigen complexes. This computational approach aims to assess an antibody’s binding potential by evaluating the predicted confidence scores, which reflect the likelihood of strong binding interactions. The tFold-Ag model, trained exclusively on binding antibody-antigen pairs, assigns lower confidence scores to unstable, non-binding complexes, reflecting their reduced likelihood of forming stable structures. Informed by previous work [30], we expect that the confidence scores will serve as a reliable metric to distinguish between binding and non-binding antibody-antigen pairs.

We conduct virtual screening experiments on two target antigens, including the programmed cell death protein 1 (PD-1, ligand: PD-L1), and the receptor-binding domain (RBD, receptor: ACE2) of the spike protein in SARS-CoV-2, to evaluate the performance of tFold-Ag in virtual screening, i.e., identifying binding antibodies from a pool of candidate antibody sequences.

As illustrated in Fig. 3a, tFold-Ag demonstrates a remarkable ability to differentiate PD-1-binding antibodies from non-binding antibodies (AUC = 0.76), based on the computed confidence scores. Remarkably, within the top 1% of the antibody ranking list, as determined by the tFold-Ag confidence score, we observe an enrichment factor (EF^1%^) of 7.41. This means that out of 7 antibodies identified by the model as high-confidence binders, 2 are positive samples, as confirmed in Supplementary Table B7. Consequently, for PD-1, a strong correlation exists between the tFold-Ag confidence score and the antibodies’ antigen-binding capability, which indicates the availability of tFold-Ag in virtual screening. Subsequently, we assess the validity of our predicted PD-1-antibody complex structures. All the anti-PD1 antibodies used in the evaluation are sourced from Thera-SAbDab and are either in clinical use or approved. Therefore, these antibodies are supposed to be capable of inhibiting the PD1-PDL1 interaction, serving as their pharmacological function. Using tFold-Ag, we predict the structure of Lipustobart, the top-ranked antibody, and confirm the similarity of its binding site to PDL1 and its potential for competitive binding with PD1 through structural superimposition with the PD-1/PD-L1 complex (PDB ID: 4zqk [31]), shown in Fig. 3b. In evaluating the virtual screening performance for the SARS-CoV-2 RBD, we employ antibodies from a dataset of single-B cell repertoire sequencing, distinct from the donors of the antibodies in the training set. This approach enables us to assess the model’s predictive accuracy on novel data distributions, thereby confirming its generalizability. The normalized number of effective sequences (Neff) for the MSA of the SARS-CoV-2 RBD is 4.9, as opposed to 8.3 for PD-1. This indicates that the RBD has fewer diverse homologous sequences than PD-1, making the prediction more challenging. Moreover, the cutoff date for tFold-Ag’s training set is 31 December 2021, and the majority of antibody structures capable of binding with the SARS-CoV-2 RBD are released in 2022 or later. This situation heightens the prediction difficulty due to the limited samples encountered by the model during the training stage. In this context, tFold-Ag achieves an AUC of 0.70 (Fig. 3d) and an EF^1%^ of 2.35, suggesting moderate success in discriminating binding from non-binding antibodies, despite the prediction challenges posed by the novel antigen and its limited homologous sequences. By applying the Kabsch algorithm, we superimpose a high-confidence predicted antibody-antigen structure with the RBD/ACE2 complex (PDB ID: 6m0j [32]), revealing overlapping epitopes and spatial clashes (Fig.3e) indicative of competitive binding, which was experimentally validated. However, due to the relatively low confidence scores of tFold-Ag’s predictions for the antibody-RBD complex, the predicted epitopes are close, which complicates the assessment of competitive potential based solely on structural data. This means that while the predicted structures can provide some insights, they may not be entirely accurate, and this uncertainty can make it difficult to definitively determine whether an antibody is competitive or not based solely on these predictions. More details of our analysis are given in the appendix. C.6.

**Fig. 3:**
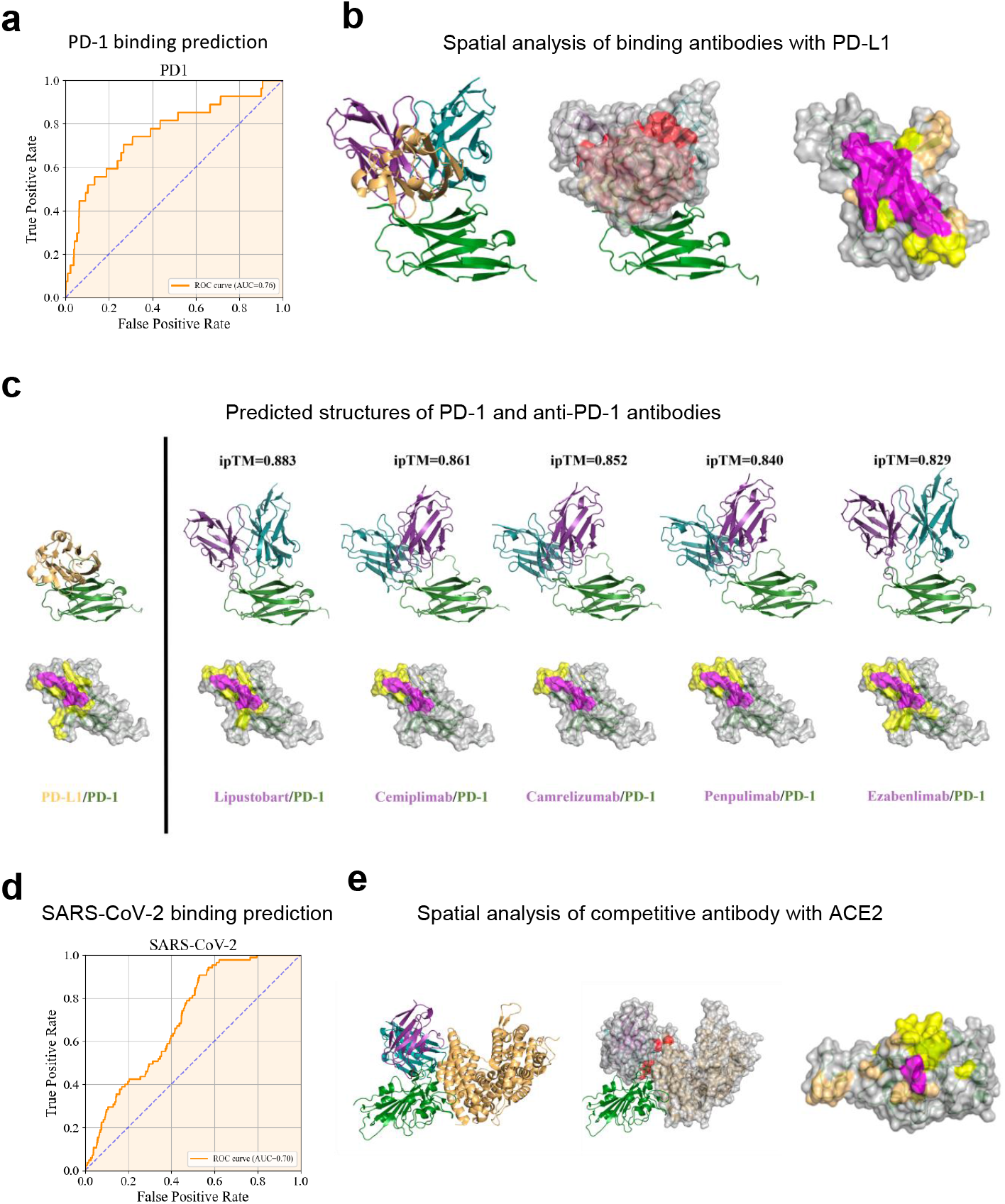
Structure-based virtual screening of binding antibodies using tFold-Ag. (a) ROC curve for PD-1 set in distinguishing between binding and non-binding antibodies. (b) Spatial analysis of binding antibodies (Lipustobart, ranked 1st by confidence predicted by tFold-Ag, blue for heavy chain, purple for light chain and green for antigen chain) with PD-L1 (PDB ID: 4zqk, orange for ligand and green for antigen chain). The antibody-antigen structure is superimposed with the antigen-receptor structure by applying the Kabsch algorithm to the shared antigen component in both complex structures. The competing antibodies exhibit spatial clashes (highlighted in red) and their antigenic epitopes overlap with the binding interface between the antigen and its ligand (shown in purple). (c) tFold-Ag predicted structures of PD-1 in complex with anti-PD-1 antibodies. The selected antibodies were the top 5 positive sample antibodies of our method, all with confidence scores greater than 0.8. The ribbon models (top) and surface models (bottom) of PD-L1 are displayed in the same orientation, and the antibody heavy and light chains are colored blue and purple, respectively. The epitope on the surface of the PD-1 is colored yellow while the shared regions on the epitopes of the five antibodies and the PD-L1 binding site are colored purple. (d) ROC curve for SARS-CoV-2 set in distinguishing between binding and non-binding antibodies. (e) Spatial analysis of a competitive antibody (ranked 4th with true competitive label by confidence predicted by tFold-Ag, blue for heavy chain, purple for light chain and green for antigen chain) with ACE2 (PDB ID 6m0j, orange for receptor and green for antigen chain). The competing antibodies exhibit spatial clashes (highlighted in red) and their antigenic epitopes overlap with the binding interface between the antigen and its ligand (shown in purple).

In summary, the evaluation results validate tFold-Ag’s proficiency in identifying binding antibodies by accurately reconstructing antibody-antigen complexes. It is important to note that while MSAs of antigens are still required and play a crucial role in this process, the MSAs of antibodies are not necessary, which offers a significant speed advantage in high-throughput virtual screening. For a specific antigen, this structure-based screening approach, distinct from Ligand-Based Drug Design (LBDD), can identify binding antibodies to novel antigens unseen in the training set. This broadens its applicability for single B-cell receptor repertoire sequencing data and supports de novo antibody design in complex pharmaceutical scenarios.

### 2.4 Co-design of structure and sequence for therapeutic antibodies with tFold

As previously mentioned in section 2.2, tFold-Ag is a multitask model that incorporates two heads i.e. with a structure module for protein complex conformation prediction (complex conformation prediction head) and a sequence recovery module for antibody design (antibody design head). In the antibody design phase, tFold-Ag is capable of predicting the structure of antigen-antibody complexes while designing antibody sequences due to the two-head design, making it a structure and sequence co-design approach. During the training phase, tFold-Ag is optimized to perform both tasks simultaneously. The training mechanism for complex structure prediction follows the similar approach as existing methods [5, 17], utilizing sequences as input and experimentally determined structures as the ground truth for supervision. The training mechanism for antibody design is specially designed. To enable the model to acquire the ability to design novel functional antibodies, we note the expected design regions of input antibody sequences (for instance CDRs on the heavy and light chains) by using the mask token. These masked antibody sequences, along with the antigen sequence, are then used as input. The original unmasked antibody sequences serve as supervision for the antibody design head, while the corresponding experimentally determined structures are used as supervision for the complex conformation prediction head. Through this training mechanism, tFold-Ag is capable of considering the interaction between antibodies and antigens during prediction, which is compelled to learn to generate novel antibodies with reasonable sequences and similar structures to their natural references.

To jointly optimize tFold-Ag for both complex structure prediction and antibody design tasks, we implemented a three-stage training process. In the initial training stage, the focus is on training the model for structure prediction, hence the sequence-masking process is not included. However, in the subsequent two fine-tuning stages with different parameters such as the learning rate and sequence crop size, we introduce a random masking process on the CDRs of antibodies with a 30% probability of masking an amino acid (70% probability of keeping its original) within the regions to co-train the model for both tasks (Referring to the Appendix section C.5.6 for details).

Figure 4a depicts the tFold-Ag process during the antibody design phase. It involves antibody sequences with masked regions and the sequence of the targeted antigen as input and outputs designed novel antibody sequences and the structure prediction of the antibody-target antigen complex. As previously stated in section 2.2, tFold-Ag permits the integration of interaction interface information to enhance its performance in structure prediction, such as tFold-Ag-ppi employing the Protein-Protein Interaction (PPI) information and tFold-Ag-contact utilizing the comprehensive contact information. Similarly, tFold-Ag allows for the incorporation of this information as an additional feature to benefit antibody design.(Referring to the method section 4.5 for details). The advantages of tFold-Ag for antibody design are twofold. Firstly, it is a sequence and structure co-design method, which comprehensively ensures the suitability of the generated antibody from both sequence and structure perspectives, whereas conventional methods typically focus on only one of these aspects. More importantly, tFold-Ag does not require any prior information for inference, unlike existing methods that rely on prior knowledge such as experimentally determined antibody-antigen complex structures [33], the structure of the antibody [34, 35], or the structure of antigen along with the binding epitope [12, 36].

**Fig. 4:**
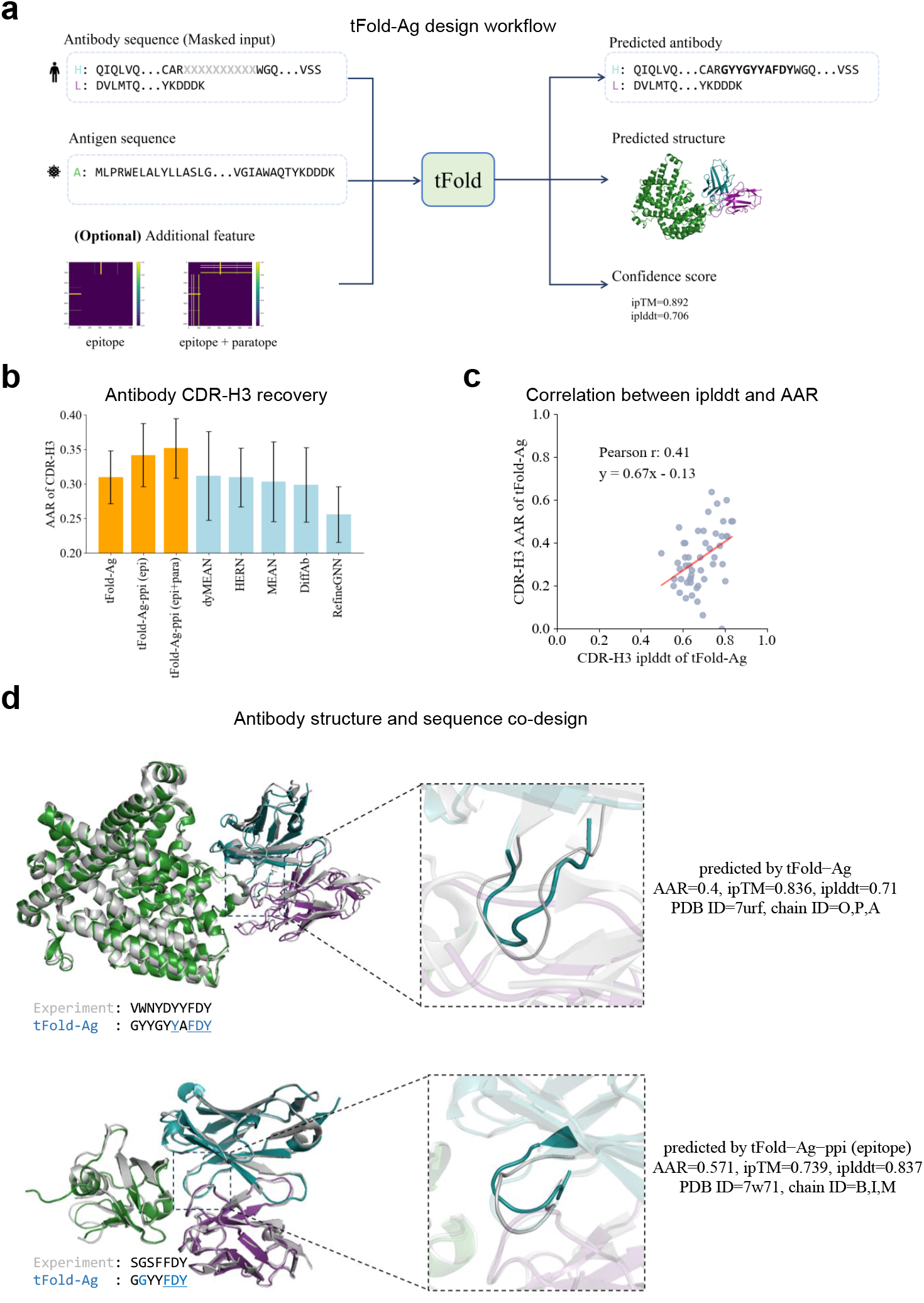
Antibody structure and sequence co-design using tFold-Ag. (a) The workflow of structure-based antibody CDRs design using tFold-Ag. The grey X denotes the masked amino acid. (a) Performance of antibody amino acid recovery for CDR-H3 of tFold-Ag and other antibody design methods on SAbDab-22-Design-Ab test set. (c) Correlation between iplddt and AAR of CDR-H3. Least-squares linear fit AAR = 0.67 × iplddt - 0.13 (Pearson’s r = 0.41, P-value = 0.003). n = 50 protein chains in SAbDab-22-Deisng-Ab. (b) Comparison of our predicted structures and CDR-H3 amino acid type for an antibody target for 7urf and 7w71 (blue for heavy chain, purple for light chain) with their respective experimental structures (gray).

We assess the performance of tFold-Ag in designing CDR loops for both chains of the antibody through in silico evaluation. For this purpose, we curated two test sets, termed SAbDab-22-DesignAb (including 50 antibody-antigen pairs) and Cov-AbDab-DesignAb (77 antibody-antigen pairs), respectively. We assessed the generalization ability of tFold-Ag to generate binding antibodies for unseen antigens on the SAbDab-22-DesignAb set and novel antibodies for target antigens on the Cov-AbDab-DesignAb set. The performance of tFold-Ag to generate binding antibodies for unseen antigens is illustrated in Fig. 4b and supplementary Table B8, where it is also compared with other existing methods that require prior knowledge, including DiffAb [34], RefineGNN [37], MEAN [35], HERN [36], and dyMEAN [12]. We employ the Amino Acid Recovery (AAR) and Contact Amino Acid Recovery (CAAR) metrics for in silico evaluation. It is worth noting that a higher AAR/CAAR is not necessarily better, and a sequence that doesn’t match the original one doesn’t automatically imply an ineffective antibody design. We observed that tFold-Ag, without utilizing any prior information, achieves the CDR-H3 AAR of 0.309 and 0.267 in SAbDab-22-DesignAb and Cov-AbDab-DesignAb, respectively. This recovery rate is comparable to the performance of the current leading method, dyMEAN, which, however, necessitates the inclusion of prior epitope information. When evaluating the recovery rate of residues in contact, tFold-Ag yields a CAAR of 0.212. We then evaluate the benefit of utilizing interchain features for tFold-Ag in antibody design. As illustrated in Table B8, we observe that incorporating the PPI information can dramatically improve the design performance (tFold-Ag-ppi vs tFold-Ag). When epitope information is input into the model, the CDR-H3 AAR and CAAR for SAbDab-22-DesignAb are enhanced to 0.347 and 0.251 respectively, outperforming dyMEAN (AAR=0.310, CAAR=0.244). This result highlights the importance of the epitope for CDR-H3 design. The recovery rates of tFold-Ag in CDR-H2 and CDR-H1 are not as good as dyMEAN [12]. This is because CDR-H2 and CDR-H1 are both encoded by the V gene, and a decent performance (AAR=0.726 for CDR-H1 and 0.427 for CDR-H2) can be achieved merely by using the sequence-based model ESM-PPI (Table B8). When tFold-Ag incorporates antigen structure information for these regions, it often introduces interference, as CDR-H1 and CDR-H2 do not always bind with the antigen. The specific details are discussed in the Appendix section C.7.2. In addition to designing paired antibodies, tFold-Ag is also suitable for nanobody design, more details of our proof-of-concept results are given in section C.7.1.

In Fig. 4c, we investigate the correlation between the confidence score of structure prediction and the quality of the antibody design (evaluated by AAR). Our in silico experiment results suggest that although the confidence score evaluated by ipTM is not related to the quality of the design, the residue-level confidence score of our designed region, which we call iplddt, is correlated with the quality of the antibody design. Predictions with better iplddt usually lead to better-designed antibodies (Refer to Appendix section C.7.3 for detailed analysis). This allows us to use the iplddt score as a metric to assess the performance of the antibody CDR-H3 sequences designed by tFold-Ag.

In Fig. 4d, we showcase the designed antibodies generated from tFold-Ag and tFold-Ag-ppi (epitope). It’s noteworthy that tFold is capable of producing innovative antibody sequences (CDRs) with multiple amino acid modifications compared to existing sequences in an end-to-end manner, while still preserving a high structural resemblance to naturally functioning counterparts. This technique is more effective than the existing artificial-evolution methods, which necessitates multiple rounds of mutation suggesting and sequence optimization [38, 39].

## 3 Conclusion and discussion

In conclusion, tFold transcends its role as a efficient tool for fast and accurate structure prediction and shows the potential as a platform technology advancing innovation in the field of antibody design. It enables the exploration of the extensive landscape of potential antibody-antigen interactions, thereby facilitating the design of effective antibodies. By enhancing the accuracy and speed of antibody-antigen complex structure predictions and leveraging these advantages to enable applications such as virtual screening and therapeutic antibody design, we expect that tFold will provide biologists with a comprehensive tool to accelerate recent advancements in structural bioinformatics. This could potentially lead to insightful revelations in the fields of molecular biology and immunology, and foster significant breakthroughs in therapeutic applications.

### 3.1 Jointly modeling antibody and antigen jointly improve antibody structure prediction

In our study, we explored the benefits of jointly modelling the antibody and antigen during complex structure prediction. While tFold-Ag employs the modules of tFold-Ab to extract the initial sequence and pair representations of antibodies, these representations can be refined and adjusted in the subsequent AI-driven flexible docking module of tFold-Ag, taking into account the interaction between the antibody and antigen. In addition to refining the representations, the AI-driven flexible docking module plays a crucial role in contemporaneously updating the structure prediction of both the antibody and antigen. This module accepts the pre-computed *C*_*α*_ coordinates of both the antibody and antigen as input and employs Evoformer-Single stack and structure modules to identify the optimal docking conformation and refine the binding site. This process is similar to the conventionally defined protein docking process but with the added feature of allowing initial input conformation updating during docking, essentially constituting a form of flexible docking.

The proposed joint modelling approach is particularly beneficial for modelling the CDR-H3 region of the antibody, which is often disordered and heavily influenced by its interaction with the antigen. Our evaluations on the SAbDab-22H2-AbAg dataset compared the structure prediction accuracy of tFold-Ag against tFold-Ab, focusing on the antibody’s CDR-H3 region that usually contacts with the antigen (evaluation using the backbone RMSD). The results, as summarized in Table B11, initially suggested that tFold-Ag did not significantly outperform tFold-Ab in terms of CDR-H3 prediction accuracy (3.21 to 3.20Å). This was attributed to the challenges in accurately identifying the antigen’s epitope, which is crucial for the correct modelling of the antibody structure. Misidentification can lead to inaccurate predictions, as the antibody may bind to an incorrect epitope. However, when we incorporated PPI information (epitope and paratope) as an additional feature, the accuracy of tFold-Ag’s predictions improved, with a decrease in RMSD for CDR-H3 (3.07 compared to 3.21Å). This improvement was more pronounced when analyzing the subset of 28 antigen-antibody complexes that tFold-Ag successfully predicted, where the RMSD of CDR-H3 was significantly lower than that predicted by tFold-Ab (2.75 vs 3.10Å), as depicted in Fig. A1a.

Furthermore, we observed a correlation between the confidence score of tFold-Ag, measured with ipTM, and the structure prediction accuracy of the antibody CDR-H3 region. Higher ipTM scores generally indicated more accurate predictions (Spearmanr’s *ρ* of 0.22, P-value=0.03), as shown in Fig. A1b. Conversely, predictions obtained with lower ipTM scores, which may suggest incorrect epitope binding, did not demonstrate significant improvements in the accuracy of the CDR-H3 region compared to the predictions made by tFold-Ab.

These findings highlight the advantage of using tFold-Ag’s AI-driven flexible docking module to jointly model the antibody and antigen. This approach not only considers the dynamic nature of the CDR-H3 region but also enhances the overall prediction accuracy of the antibody structure, as demonstrated by both ground-truth evaluations and prediction confidence scores. The obtained results support the hypothesis that when the prediction of the epitope is accurate (or when additional epitope information is available), the joint modelling of antibody and antigen structures can lead to more precise antibody predictions, especially in the context of highly variable and complex regions such as the CDR-H3.

### 3.2 Potential approach to de novo design of antigen-binding antibody design with tFold

The successful de novo design of the CDR-H3 region suggests the potential for designing antigen-binding antibodies from scratch. The antigen-binding property of antibodies is determined by the antigen-binding fragment (Fab) domain which is composed of both the variable and constant domains from each heavy and light chain. The variable domain confers antigen specificity to the antibodies, while the constant domain provides a structural scaffold. The variable domain, also known as the variable fragment (Fv) region, contains three CDRs interspersed among four relatively conserved FRs.

The de novo design of antigen-binding antibodies aims to create novel functional variable domains for antibodies. This process entails the simultaneous selection of FR templates from a vast template space (determining the sequences of the FR1, FR2, FR3, and FR4 regions) and the previously described CDRs loop design (determining the sequences of the CDR1, CDR2, and CDR3 regions). To facilitate this, a representative FR template library can be constructed and curated using the Observed Antibody Space (OAS) database [40]. From this database, all unique FR region pairs of both heavy and light chains that appear more than five times can be collected.

During the de novo design process, the entire FR template library can be screened, and CDRs loop design can be performed to generate the sequences of CDRs on top of each candidate FR template. Subsequently, the designed antibodies with the highest prediction confidence scores can be selected as final outputs. These confidence scores can be calculated by tFold-Ag during the prediction of candidate antibody-target antigen complex structures.

The de novo design of antigen-binding antibodies offers the opportunity to create highly specific and efficient antibodies from scratch, bypassing the need for traditional hybridoma technology or phage display techniques. This could significantly expedite the process of antibody discovery and development, reducing costs and increasing the speed at which new therapies can be brought to market.

Moreover, the use of tFold in predicting candidate antibody-target antigen complex structures may enhance the accuracy and reliability of antibody design. This could lead to the creation of antibodies with improved specificity and affinity for their target antigens, potentially increasing the efficacy of antibody-based therapies.

Most importantly, the ability to design antibodies de novo could also facilitate the development of antibodies against novel or difficult-to-target antigens, broadening the range of diseases that can be treated with antibody-based therapies.

### 3.3 Limitations

While tFold has demonstrated exceptional performance in antibody-antigen complex structure prediction and antibody design, there are still several limitations to address. Firstly, tFold-Ag relies on pre-trained structural prediction models to provide the initial representations and structure of the antigen. However, antigens are diverse, ranging from monomers to short peptides and protein complexes. In cases where the antigen is not a monomeric protein, tFold-Ag’s ability to accurately model the antibody-antigen complex structure is restricted by the performance limitation of the pre-trained structural models.

Furthermore, as shown in Appendix section C.5.9, tFold-Ag and AlphaFold-Multimer exhibit strong complementarity in performance, with the confidence score serving as a criterion to select the better-performing model. This highlights that, despite the scarcity of co-evolutionary information between antigens and antibodies, valuable insights can still be extracted from paired MSAs. However, the use of antibody MSAs incurs a significant computational cost. Balancing the trade-off between time expenditure and performance loss presents a challenge that needs to be addressed. Additionally, in Section 2.3, we demonstrated that the structural prediction confidence score of tFold-Ag (ipTM) could be utilized for structure-based virtual screening of binding antibodies. However, this approach is accompanied by a high rate of false positives. Since the functionality of antibodies is closely related to their binding affinity with antigens, incorporating an additional prediction head to estimate affinity could be a possible solution that brings valuable enhancement.

Beyond these limitations, there is considerable room for improvement within the tFold framework. tFold-Ab has shown that pre-training with general proteins can enhance the accuracy of antibody predictions. Theoretically, this approach could also improve the predictive performance of tFold-Ag, especially for Nanobody-Antigen complexes, which are often underrepresented in data. Besides, self-distillation has been proven to be an effective strategy in the field of structural prediction [17]. We have not yet implemented self-distillation for tFold. Given that the OAS [40] database contains a vast array of antibody sequences without experimentally determined structures, and PLAbDab [41] includes over 150,000 pairs of antibodies and antigens known to bind, there is substantial potential for future accuracy improvements if incorporating the self-distillation strategy.

## 4 Methods

### 4.1 Data

#### 4.1.1 Datasets for the prediction of antibody structures

We construct the training, validation and test sets from the SAbDab database [27] based on the widely applied temporally separated protocol [17, 18]. Specifically, we curate all the experimentally determined antibody structures released before 31 December 2021, to constitute the samples in the training set. The resulting training set consists of 8,264 antibody complexes containing both heavy and light chains, 1,693 antibody samples with only heavy chain information, and 376 antibody samples with only light chain information. In the training phase, we grouped the training samples into various clusters referring to the sequence similarity calculated by the CD-Hit [42]. Each cluster is composed of antibodies exhibiting greater than 95% sequence identity, resulting in a total of 2,873 clusters. During each training epoch, we randomly select one antibody sample from each cluster Data to form the training samples for the current epoch. The validation set consists of antibody structures released between 01 January 2022 and 30 Jun 2022, on which we perform hyper-parameter tuning and model selection. Sequences in the validation set with high similarity to those in the training set (sequence identity greater than 95%) are removed. The test set consists of two non-redundant benchmark subsets (sequence identity lower than 99% among included samples), i.e, SAbDab-22H2-Ab and SAbDab-22H2-Nano, which include 170 antibodies and 73 nanobodies released between 01 July 2022 and 31 December 2022 respectively. Similarly, we eliminate redundant samples in the test set that have 100% sequence identity to those in the training set, as a larger sequence identity threshold is advantageous in investigating the model’s sensitivity to mutations.

#### 4.1.2 Datasets for antibody-antigen complex prediction

With a similar protocol, we create the training, validation, and test sets of antibody-antigen complex structures based on the SAbDab database [27]. The training set consists of experimentally determined antibody-antigen complex structures (4,834 samples) and nanobody-antigen complex structures (1,319 samples, including instances of single domain antibody-antigen complexes) released before 31 December 2021. During the training phase, we grouped training samples into clusters based on antigen sequence similarity (samples with over 95% sequence identity of the antigen are assigned to a cluster), resulting in a total of 2,459 clusters. In each training epoch, a single sample is randomly chosen from each cluster to constitute the training samples for the current epoch. The validation set comprises antibody-antigen and nanobody-antigen complexes released between 01 January 2022 and 30 Jun 2022, comprises 99 antibody-antigen complexes and 40 nanobody-antigen complexes. The curated test set, comprising a total of 99 antibody-antigen complexes and 43 nanobody-antigen complexes, is composed of antibody-antigen complexes and nanobody-antigen complexes released between 01 July 2022 and 31 December 2022. It is organized into two non-redundant benchmarks, dubbed SAbDab-22H2-AbAg and SAbDab-22H2-NanoAg, representing antibody-antigen and nanobody-antigen sets, respectively (sequence identity lower than 95% among included samples). To ensure a fair comparison with existing methods such as AlphaFold-Multimer [5], we excluded all structures of the test set for the training phase of all evaluated methods. Additional data pre-processing steps for the test set include (1) removing samples that share 95% sequence identity with those in the training set, (2) excluding antibody-antigen and nanobody-antigen samples without contact, and (3) eliminating samples with antigen sequence lengths exceeding 600.

##### MSA generation

For each antigen chain in SAbDab, we produced its multiple sequence alignment (MSA) by executing MMseqs2 following the default ColabFold pipeline. This was performed on the UniRef30 [16] library of March 2021, and the colabfold envdb [43] as of August, 2021.

#### 4.1.3 Datasets for virtual screening of binding antibodies

In order to evaluate the efficacy of tFold-Ag in the high-throughput screening of binding antibodies, we constructed two test datasets to perform in silico virtual screening, with each targeting a different antigen to assess the efficacy of our approach. One test dataset focuses on the PD-1 antigen, while the other is dedicated to the wild-type strain of the SARS-CoV-2 virus.

##### PD-1 set

To evaluate the capability of tFold-Ag in distinguishing authentic antibody therapeutics from non-therapeutic antibodies, an assessment was conducted with a dataset that included 27 anti-PD-1 therapeutic antibodies and 718 random antibodies from healthy human donors. The therapeutic antibodies were obtained from Thera-SAbDab [44], a database that curates all antibody therapeutics recorded by the World Health Organization (WHO) with sequences available to the public. To ensure the integrity of the evaluation, 9 antibodies with existing experimental structures were excluded to prevent potential bias, as tFold-Ag may have used these structures during training. The 718 random antibodies, which are paired in heavy and light chains, were obtained from the OAS database [45], which were originally derived from the healthy human donors in an earlier study of human rhinovirus infection [46]. In this evaluation, tFold-Ag was utilized to identify the 27 anti-PD-1 therapeutic antibodies from the set of 745 antibodies, given their sequence data only.

##### SARS-CoV-2 set

To simulate real-world scenarios in drug development, we constructed a test set using a single-B repertoire sequencing dataset obtained from the OAS database [40]. This dataset was initially derived from mice immunized with the SARS-CoV-2 spike protein [47], encompassing 11,388 antibodies with paired heavy and light chains. We excluded 125 antibodies with incomplete VH/VL domains, likely due to sequencing errors, from the analysis. The subsequent deduplication process yielded a set of 1,595 distinct antibodies with unique VH and VL sequences. Notably, none of these antibody sequences had corresponding structures in the SAbDab database. In the original study [47], functional characterization was conducted on 26 antibodies, specifically evaluating their binding to the SARS-CoV-2 receptor-binding domain (RBD) using ELISA (enzyme-linked immunosorbent assay) and their competitive inhibition of the angiotensin-converting enzyme 2 (ACE2) using BLI (Biolayer interferometry). In the present analysis, 85 antibodies were identified as RBD binders and 72 as ACE2 blockers. These antibodies were determined to be clonally related to the previously characterized antibodies through the alignment of their germline gene identifiers and the sequences of the heavy/light chain CDR3. In this evaluation, tFold-Ag was tested to distinguish RBD binder from the set of 1,595 antibodies, given their sequence data only.

#### 4.1.4 Datasets for antibody design

To evaluate the generalizability of our model across different antibodies and antigens, we constructed two distinct test sets: one aimed at antigen generalization, and the other focused on antibody generalization.

The first, named **SAbDab-22-DesignAb**, was derived from the SAbDab database, by selecting antigen sequences that cluster at a 70% sequence identity threshold. SAbDab-22-DesignAb contains antibody-antigen complex structures in SAbDab that were released from 01 January 2022, through 31 December 2022. We excluded antibody-antigen pairs in SAbDab-22-DesignAb that any antigen sequences shared more than 70% identity with sequences in our training set to avoid redundancy. Additionally, we omitted complex structures with an experimental resolution larger 2.5 Å to maintain the accuracy of prior features. To ensure the design task was relevant, we selected only those CDR-H3 regions requiring design that were in contact with the antigen, defined as being within a 10 Å distance. These criteria yielded a total of 50 antibody-antigen structures. Given the availability of actual structures within the dataset, we were able to extract additional input information such as protein-protein interaction (PPI), antigen epitope, and contact details from these real structures. We utilized this set of 50 structures to evaluate the model’s design performance across various inputs.

The second test set, named **Cov-AbDab-DesignAb**, was extracted from the CovAbDab [48] database. This set included all antibodies from Cov-AbDab capable of binding to the antigen, excluding those with 100% sequence identity to antigen in the training set. By clustering antibody sequence pairs based on a 70% sequence identity threshold, a total of 77 antibody-antigen pairs were obtained. It should be noted that Cov-AbDab exclusively collects antibodies with the ability to bind (or nonbind) to coronaviruses, implying that the antigen sequences in Cov-AbDab-DesignAb are similar. The clustering of these antibody sequences provides a valuable method to evaluate the proficiency of our design model in engineering antibodies across a variety of antibody frameworks. A majority of the antibody-antigen pairs in Cov-AbDab lack experimental structures, which means we cannot extract additional input from these structures. To address this, we employed AlphaFold2 [17] to generate predicted structures of the antigens for those design methods requiring antigen structure.

We refrained from utilizing the previously commonly employed benchmark RAbD [49], as the antigen-antibody complex structures contained within it have already been represented in our training set.

Our model can also be applied to design nanobody using the same approach. However, due to the lack of suitable methods for comparison, we constructed a test set called SAbDab-22-DesignNano and demonstrated the performance of our model on this test set. The test set and the results are presented in Appendix C.7.

### 4.2 Evaluation criteria

For antibody structure prediction, we present the backbone root-mean-square deviation (RMSD) for various framework and CDRs. We use Chothia [50], a structure-based numbering scheme for antibody variable regions. Additionally, we compare OCD (orientational coordinate distance) [25] and DockQ score to verify how well the relative position between heavy and light chains is estimated.

For antibody-antigen complex prediction, we report DockQ score and success rate (SR) as determined by DockQ algorithm [51]. Further, we employ the TM-Score to evaluate the overall prediction. Moreover, we analyze the correlation between the confidence scores provided by different prediction methods and their respective prediction accuracy using the coefficient of determination (*R*^2^). For the prediction-based model, we use the same genetic databases to build a multiple sequence alignment (MSA) and the same template databases to extract template features. For the docking-based model, the antibody structure is generated by AlphaFold-Multimer while the antigen structure is generated by AlphaFold2. When prior information such as protein-protein interactions (PPIs), antigen epitope and inter-chain contact are specified, the extra features are calculated from the native complex. In methods that yield energy as an output, the results are organized according to their respective energy levels. The model demonstrating the lowest energy is subsequently chosen for evaluation. On the other hand, for methods that generate a confidence score, the optimal output is discerned from a plethora of model predictions. For instance, in the case of AlphaFold-Multimer, which generates 25 predictions, the selection is made based on the confidence scores, as the confidence score measures the level of certainty associated with each predicted conformation.

For the task of antibody design, we employ the following metrics for in silico evaluation: Amino Acid Recovery (AAR), defined as the overlap ratio between the generated sequence and the known binding antibody; Contact Amino Acid Recovery (CAAR) [52], which calculates the AAR specifically for binding residues located within 6.6Å of epitope residues.

### 4.3 Antibody structure prediction using tFold-Ab

As illustrated in Fig. 1a and Algorithm 1, the proposed antibody structure prediction pipeline mainly consists of four components: 1) initial antibody feature extractor using pre-trained language models; 2) Evoformer-Single stack for iterative update of sequence embedding; 3) structure module using invariant point attention (IPA) [17] for structure prediction; and 4) additional refinement with recycling iterations.

#### 4.3.1 Extract multimer inter-chain information using language model

Pre-trained language models (PLM) for protein sequences have been widely demonstrated to effectively capture the dependency among residues and thus provide meaningful feature embeddings [15, 53–56]. These models are typically trained with a masked language modeling (MLM) loss, where randomly masked amino-acid sequences are fed into the network and it is tasked with recovering the original ones via self-attention. The attention weights in these models reflect how different residues interact with each other, and can therefore be naturally used as initial pair features between residues.

A recent study [23] has confirmed that the integrating chain relative positional encoding with pre-trained language models, and further pre-training the language model using group sequences derived from PPIs and complexes in PDB, enables the language model to extract inter-chain information. We modified the architecture of ESM-2 [15] to accommodate both single sequence and pair sequence inputs, ensuring the preservation of the efficacy of the pre-trained parameters. The ESM-2 model, which comprises 650 million parameters, has been selected as the base model. During the further pre-training phase, only self-attention with chain relative position embedding, adding no more than 1 million parameters, has been incorporated.

Our PLM model, named ESM-PPI (ESM base model with protein-protein interaction preservation), uses the algorithm proposed in [23] to distinguish different chains for further pre-training. We have also adapted the masking strategy of protein language models, tailoring it to pair antibody sequences and homomer sequences. Additionally, we have developed a novel cropping method to ensure that the cropped sequences maintain their interaction. The data and detailed information for ESM-PPI are introduced in Appendix C.3.

#### 4.3.2 Sequence-based antibody prediction

In our approach, we begin by extracting the final sequence embeddings and all-layer pairwise attention weights for each heavy and light chain from the pre-trained ESM-PPI model, keeping the parameters fixed. To distinguish clearly between heavy and light chains, we employ three distinct positional encoders: the multimer positional encoder, the relative positional encoder, and the chain relative positional encoder. The specifics of the sequence embedding module for tFold-Ab are detailed in Algorithm 2. These encoders are designed to incorporate additional positional information into the initial sequence embedding and pairwise representation, enriching the model’s input data.

The advanced structure prediction [15, 17] consists of two stages: iterative updates of single or MSA features and pair features, followed by a structure module for 3D structure prediction. As illustrated in Fig. 1a, our architecture is inspired by the design of AlphaFold [17]. We introduce a nuanced alteration where the standard Evo-former stack, responsible for updating MSA and pair features, is substituted with a simplified Evoformer-Single stack tailored for single sequence inputs (please refer to Appendix C.4.3 for details).

Both antibodies and nanobodies are processed using the same model parameters within the Evoformer-Single and structure modules, as we have empirically discovered that this approach is effective due to two reasons: 1) the antibody and nanobody features are derived from the same pre-trained language model, ensuring consistency in the feature distribution; and 2) the independent positional encodings provide chain-type specific information that is crucial for accurate antibody structure prediction. This allows us to treat the input of antibodies and nanobodies as a single chain for the purposes of our model. Additionally, we have incorporated a recycling strategy, as outlined in Algorithm 7 and Appendix C.4.4. This strategy effectively deepens the network, enhancing the sequence and pairwise representations within the Evoformer-Single stacks without necessitating an increase in the number of parameters.

#### 4.3.3 Improve the generalizability of models through pre-training

Antibodies and nanobodies, while differing in their CDRs, share considerable structural similarities in other areas. Besides, the number of these entities with experimental structures in the PDB is relatively small. This poses a risk of overfitting when models are trained exclusively on datasets comprising only antibodies and nanobodies. To mitigate this risk, we have adopted a pre-training strategy that utilizes a diverse array of general monomers and multimers sourced from the PDB. Given that an antibody is a complex protein comprising two entities and a nanobody is a monomeric protein, this strategy significantly expands the breadth of the training dataset. Consequently, it enhances the model’s ability to generalize and reduces the likelihood of overfitting. The details of pre-training on general proteins are presented in Appendix C.4.6.

Our ablation studies, detailed in Appendix C.4.8, provide empirical evidence supporting this approach. The results demonstrate that pre-training on general proteins markedly improves prediction accuracy across different regions. This improvement is particularly notable in the SAbDab-22H2-Nano test set, underscoring the efficacy of our methodology in producing robust and accurate structural predictions.

### 4.4 Antibody-antigen complex prediction using tFold-Ag

The overall network architecture of tFold-Ag is shown in Fig. 2a, which consists of three modules: the antibody feature generation module, the antigen feature generation module, and the AI-driven flexible docking module. As shown in Algorithm 8, the antigen and antibody sequences are fed into dedicated modules designed to extract sequence features, MSA features and the initial coordinates of antibody and antigen. Subsequently, these features are integrated within the AI-driven flexible docking module to predict the structure of the antigen-antibody complex. Additionally, the predicted structures are accompanied by a series of confidence scores, which serve as an indicator of the reliability of the prediction.

#### 4.4.1 Generate features of antibodies and antigens using pre-trained structure prediction models

In the tFold-Ag, the antibody feature generation module takes antibody sequences as input and utilizes the pre-trained tFold-Ab model to produce sequence embeddings, pair representations, and initial coordinates.

Conversely, our antigen feature generation module faces a distinct challenge, as highlighted by our experiments detailed in Table B10. We observed that sequence-based prediction models, despite their rapid processing time, fall short in accurately predicting antigen structures when compared to MSA-based models. This discrepancy is likely attributable to the vast diversity inherent to antigens. To address this issue and improve the model’s generalizability, we opted to employ structure prediction models pre-trained on a wide array of general proteins for extracting antigen features. Recent works [57, 58] have demonstrated the Evoformer’s capability to encode both structural and functional properties of proteins from MSA data. Leveraging this insight, we construct an MSA by querying genetic databases, then extract MSA embeddings and pair representations from the final layer of the Evoformer stack within the pre-trained AlphaFold2 model. We also gather the predicted coordinates for the antigens.

Furthermore, as indicated in Table B10, disabling the recycling strategy in AlphaFold does not significantly impact the performance of antigen monomer structure prediction (with TM-scores of 0.817 vs 0.864). Our ablation study in Section C.5.7, corroborates this finding, showing that the use of antigen features extracted by the recycling model does not substantially improve the performance of predicting antibody-antigen complexes. Consequently, we have chosen to disable the recycling strategy in the pre-trained AlphaFold2 when extracting antigen features.

#### 4.4.2 AI-driven flexible docking module

In the docking module of our tFold-Ag method, we hypothesized that neural networks are capable of accurately reconstructing the 3D coordinates of antibodies and antigens from their respective features. To better capture the intricate correlation between the antigen and antibody, we have devised a feature fusion module. For the latest multimer prediction method [5, 28], the pairing MSAs are used to generate the multimer initial features for Evoformer stack’s input. For the multimer prediction of the antibody-antigen complex specifically, we concatenate their amino acid sequences without any delimiters. This is mirrored in the feature space, where each chain’s single features are also concatenated along the sequence dimension to serve as the initial single features for multimer structure prediction. By considering the antibody as a single entity (whether it is in a paired state or not) and by incorporating positional encoding, we ensure that the model is informed of the chain type and position of each residue.

As illustrated in Fig. C4 and detailed in Algorithm 9, we partition the pair features for multimer structure prediction into four segments. The diagonal blocks are populated with pair features and initial coordinates from the antibody and antigen chains. For the off-diagonal segments, we employ the ‘Outer product mean’ from Algorithm 10 to transform the antigen MSA representation and antibody sequence representation into an inter-chain pair feature. Additionally, for the initial sequence feature, we utilize cross attention mechanisms [59] to generate sequence embeddings for the antibody-antigen complex. This enables each residue in one sequence to attend to all residues in the other sequence.

Following the feature fusion module, we feed the initial single and pair features into a subsequent sub-network. This sub-network is composed of 32 Evoformer-Single blocks and 8 structure modules, which share parameters, for the prediction of multimer structures. Although utilizing the same subnetwork architecture as tFold-Ab, the task of identifying binding epitopes in antibodies and antigens presents a significantly greater challenge than differentiating between the light and heavy chains of antibodies. To address this complexity, tFold-Ag is equipped with a larger number of model parameters, enabling it to effectively model the complex interactions between antibodies and antigens. This design choice is pivotal, as it allows the model to refine feature embeddings within the global conformational context of the antibody-antigen complex.

#### 4.4.3 Improving accuracy using extra structure restraints

tFold-Ag has demonstrated commendable proficiency, with a success rate approaching 30%, in the prediction of antibody-antigen complexes, even in the absence of prior knowledge. However, the integration of structural restraints, sourced from experimental techniques, can significantly amplify its predictive accuracy. These additional restraints, encompassing the protein-protein interactions (PPI) (including the antigen epitope site and the antibody paratope site), and inter-chain contacts, have been shown to be highly advantageous.

For example, chemical cross-linking (XL) offers the distance between two residues connected by fixed-length reagents, which can be interpreted as a contact between the antibody and antigen. Additionally, the PPI information, including antigen epitopes and antibody paratopes, which could be obtained by Deep Mutation Scan (DMS), is also important. Despite these experimental restraints being sparse and unable to fully determine the protein complex structure, they can offer critical insights into the interaction interface of the components.

In the field of protein docking and structure prediction, the use of additional structural restraints has been investigated. Previous studies [8, 60] have included them during the conformation search phase to guide the docking towards more feasible structures. Other works [7, 61] have applied these restraints to sift through and select the most plausible structures from a pool of candidates. More recently, with the advent of AlphaFold, researchers [13] have been exploring how to combine these restraints with pre-trained models to enhance prediction accuracy.

For tFold-Ag, we’ve taken this concept a step further by developing a new interchain feature embedding module. This module is designed to seamlessly integrate various inter-chain structural features, including both sequential and pairwise features, into the model. By adding the feature embeddings through a residual connection to the original sequence representations, we ensure that the core training of the model remains unaffected. This allows the model to benefit from the additional information provided by the structural restraints without disrupting the learning process that has already been established. The details are elaborated in the Appendix C.5.3.

### 4.5 Co-design of antibody structure and sequence

To facilitate the co-design of antibody structure and sequence, we have expanded the model’s input requirements by accommodating both complete sequences and sequences with masked regions. During the design phase, the masked region signifies the location where we aim to generate new amino acids. Specifically, in our pre-trained protein language model ESM-PPI, we represent amino acids of different types as distinct tokens (N=20) and note the expected designed regions in the sequence with special *<*Mask*>* tokens. This design makes tFold-Ag suitable not only for structure prediction but also for antibody design. Subsequently, tFold-Ag integrates the features of masked antibody and unmasked antigen sequences within the feature fusion module (see Appendix section C.5.2 for details). This fusion takes into account the interaction between the masked antibody and antigen, which is crucial for effective antibody design. After processing through the Evoformer-Single Stack in the antigen-antibody complex prediction module, tFold-Ag incorporates an additional auxiliary head (implemented as feed-forward layers) to generate novel antibodies by imputing different amino acids with high predicted likelihood into the masked regions. Upon obtaining the designed complete sequence, tFold-Ag can then predict the structure of the antibody-antigen complex and provide an antibody design score. This score, referred to as iplddt, is based on the predicted local distance difference test (lddt) of the masked region, indicating the confidence level of the predictions for sequence imputation. By considering both the structural confidence score and the sequence confidence score, we can effectively evaluate our designs, with the details described in Appendix C.7.3.

To enable the model to incorporate structural information during the sequence design, we have modified our training strategy (see Appendix section C.5.6 for details). As a result, tFold-Ag is extended to be a structure and sequence co-design method for antibodies.

Considering the CDRs are the most crucial regions determining the antigen-binding specificity of antibodies, we initially utilize tFold’s co-design capability to create novel antibodies by generating new functional CDRs for those antibodies (referred to as CDRs loop design in this paper). Most current methods for structure-based antibody CDRs loop design rely on the availability of a known antibody-antigen complex structure [33], incorporate additional docking software [34, 35], or necessitate knowledge of the antigen’s epitope [12, 36]. In contrast, our approach uniquely enables the generation of the complete antibody complex using only the partial antibody sequence and antigen MSA, employing an end-to-end, full-atom framework. To the best of our knowledge, our method is the only one that doesn’t require any prior information and solely starts from the sequence data for antibody CDR design.

In the CDRs loop design task, we primarily focus on the generation of heavy chain CDR H1-H3 within antibodies. For instance, in the task of CDR-H3 design, as illustrated in Fig. 4a, we mask the entire CDR-H3 region within the known antibody sequences. These sequences, along with the antigen MSA, are then input into our tFold-Ag system, which aims to accurately predict the amino acid composition of the CDR-H3 region and compare it to that of actual antibodies. Our predictions extend beyond the amino acid types of the CDR-H3 sequence; we also provide the predicted antibody structure, a structure prediction confidence score, and an antibody design confidence score. These metrics are crucial for assessing the accuracy of our antibody-antigen complex predictions and the efficacy of our antibody design process.

It is important to note that when additional structural constraints are applied, our model does not utilize inter-chain features for the masked residues. While our methodology is equally capable of designing light chain CDRs, we have chosen to exclude these details to maintain focus and conciseness.

## 5 Data Availability

All input data are freely available from public sources. At prediction stage, for each antigen chain, we produced antigen’s MSA following the default ColabFold pipeline, against the UniRef30 [16] library of March 2021, and the colabfold envdb [43] as of Augest, 2021. Experimental structures are drawn from the same copy of the PDB; we show structures of 7OX3[62], 7RPT, 7WD1[63], 7OCJ[64], 7WSL[65], 8DF5[66], 7SAI[67], 7X2M[68], 7URF[69] and 7W71[70] for evaluation.

## 6 Code Availability

The source code, weights and inference scripts for the tFold models are available at https://github.com/TencentAI4S/tfold

## Acknowledgments

We thank for Junhong Huang, Jie Ren, Zihan Wu, Xiaoyang Jing and Xiangzhe Kong for helpful discussion; Qiang Li for web server design; Zhenxing Zhang and IPRC for computational resources. We also thank our colleagues at AI Lab, Tencent for their encouragement and support.

## Author Contributions

Conceived the study: FW, JW, JY; Trained and evaluated tFold-Ab: JW, FW; Trained and evaluated tFold-Ag: FW, JW, RW, YX; Virtual screening: FW, JW, BJ, NH, LH, HJ; Antibody design: BH, YZ, BB, HL; Code and server: CQ, FW, RW, YL; Wrote the manuscript: FW, YZ, JW, BH, FY, NH, YX, YR, TX; Offered supervision throughout the project: WL, PZ, JY; All authors read and contributed to the manuscript.

## Competing interests

The authors declare no competing interests.

## Appendix A Supplementary Figures

**Fig. A1:**
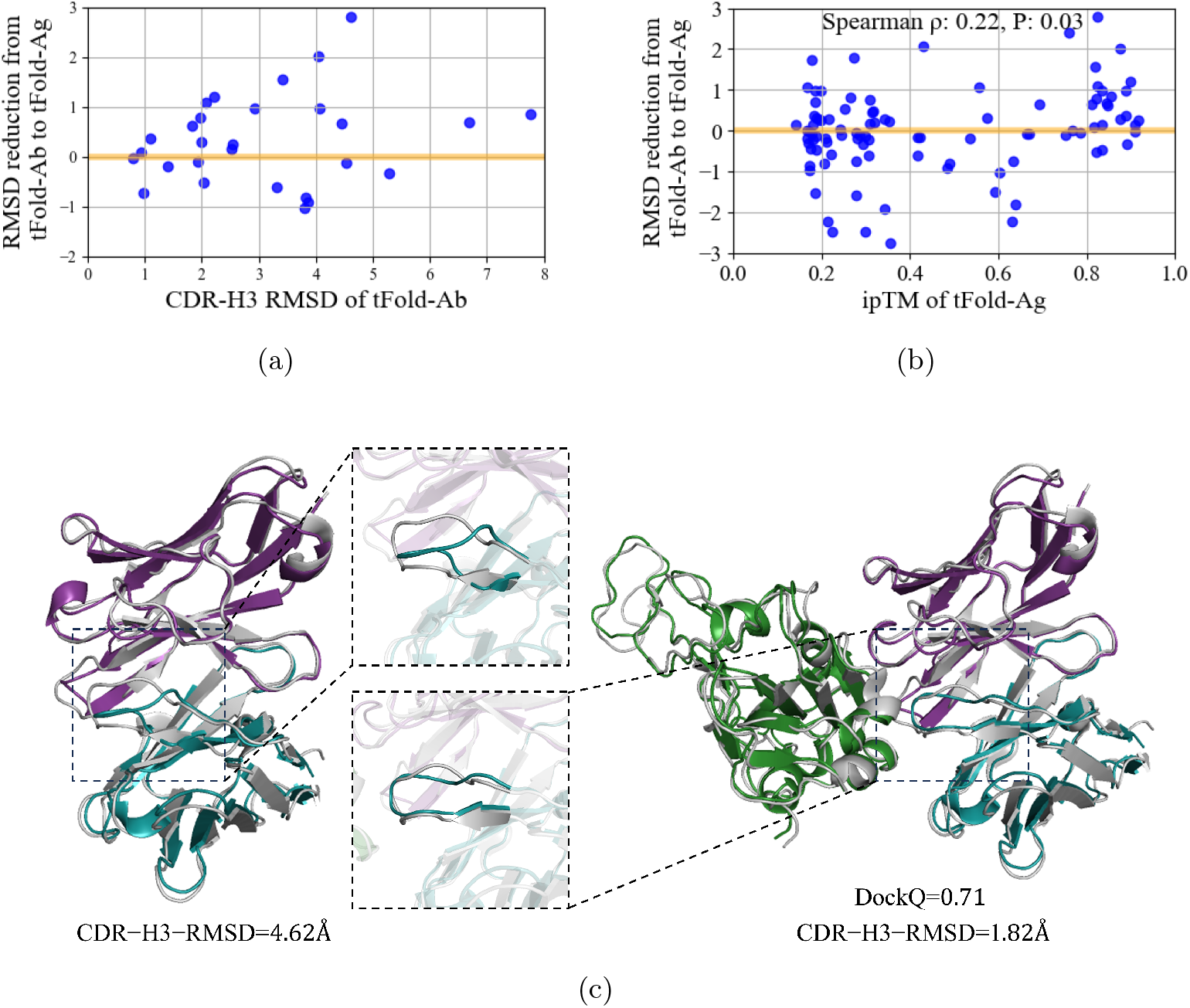
(a) Comparison of the antibody predictive performance of tFold-Ag and tFold-Ab within CDR-H3 based on SAbDab-22H2-AbAg subset, where tFold-Ag accurately predicts the epitope without prior information. The positive y-axis indicates the RMSD reduction from tFold-Ab to tFold-Ag. (b) Correlation between ipTM and RMSD reduction in antibody CDR-H3 from tFold-Ab to tFold-Ag with a Spearmanr’s *ρ* of 0.22 and a P-value of 0.03, across n = 99 protein chains in SAbDab-22H2-AbAg. (c) left: Comparison of tFold-Ab predicted structures for an antibody target (PDB 7wo7, blue for heavy chain with chain ID ‘A’, purple for light chain with chain ID ‘B’) with respective experimental structures (gray). The backbone RMSD of CDR-H3 prediction by tFold-Ab is 4.62 Å. right: Comparison of tFold-Ag predicted structures for an antibody-antigen complex (PDB 7wo7, blue for heavy chain with chain ID ‘A’, purple for light chain with chain ID ‘B’ and green for antigen chain with chain ID ‘C’) with respective experimental structures (gray). The backbone RMSD of the antibody’s CDR-H3 prediction by tFold-Ag is 1.82 Å, the DockQ of antibody-antigen interaction is 0.71.

## Appendix B Supplementary Tables

**Table B1:**
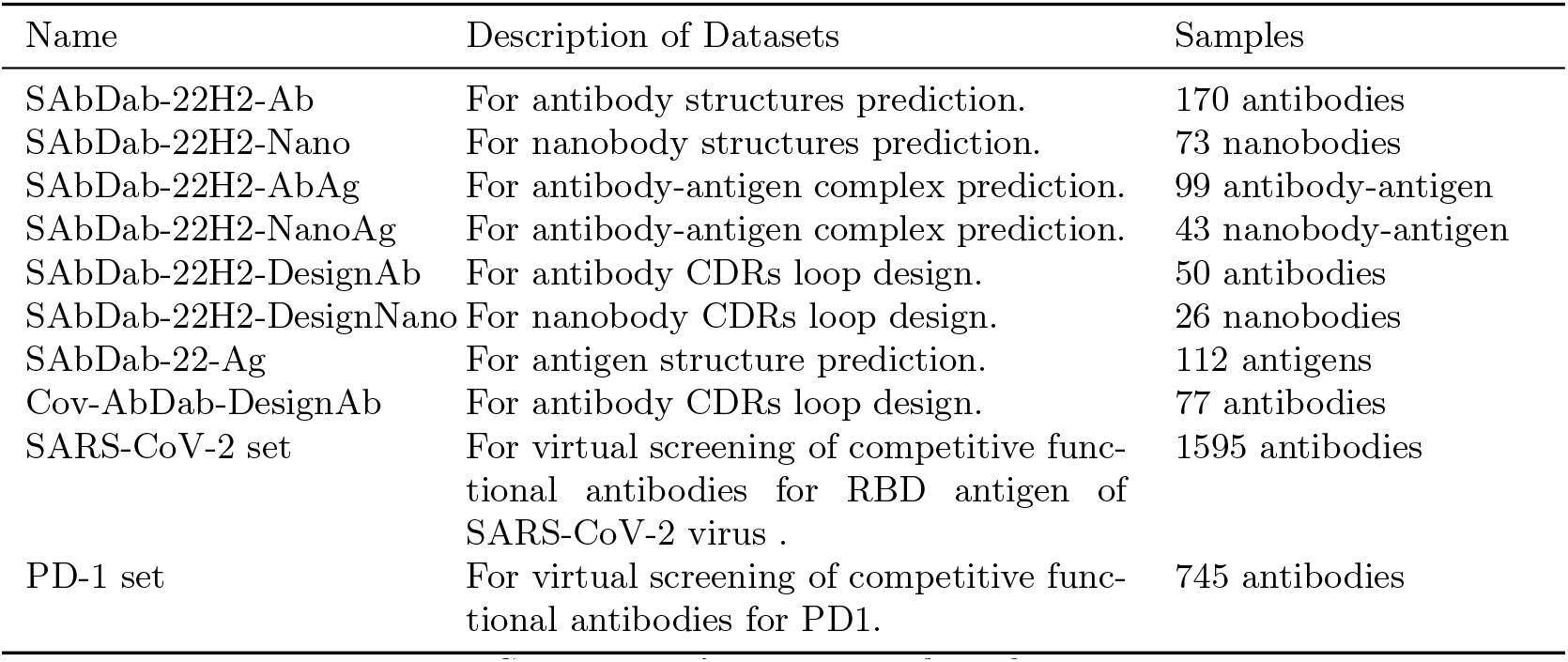
Summary of test set used in this paper.

| Name | Description of Datasets | Samples |
| --- | --- | --- |
| SAbDab-22H2-Ab | For antibody structures prediction. | 170 antibodies |
| SAbDab-22H2-Nano | For nanobody structures prediction. | 73 nanobodies |
| SAbDab-22H2-AbAg | For antibody-antigen complex prediction. | 99 antibody-antigen |
| SAbDab-22H2-NanoAg | For antibody-antigen complex prediction. | 43 nanobody-antigen |
| SAbDab-22H2-DesignAb | For antibody CDRs loop design. | 50 antibodies |
| SAbDab-22H2-DesignNano | For nanobody CDRs loop design. | 26 nanobodies |
| SAbDab-22-Ag | For antigen structure prediction. | 112 antigens |
| Cov-AbDab-DesignAb | For antibody CDRs loop design. | 77 antibodies |
| SARS-CoV-2 set | For virtual screening of competitive functional antibodies for RBD antigen of SARS-CoV-2 virus . | 1595 antibodies |
| PD-1 set | For virtual screening of competitive functional antibodies for PD1. | 745 antibodies |

**Table B2:**
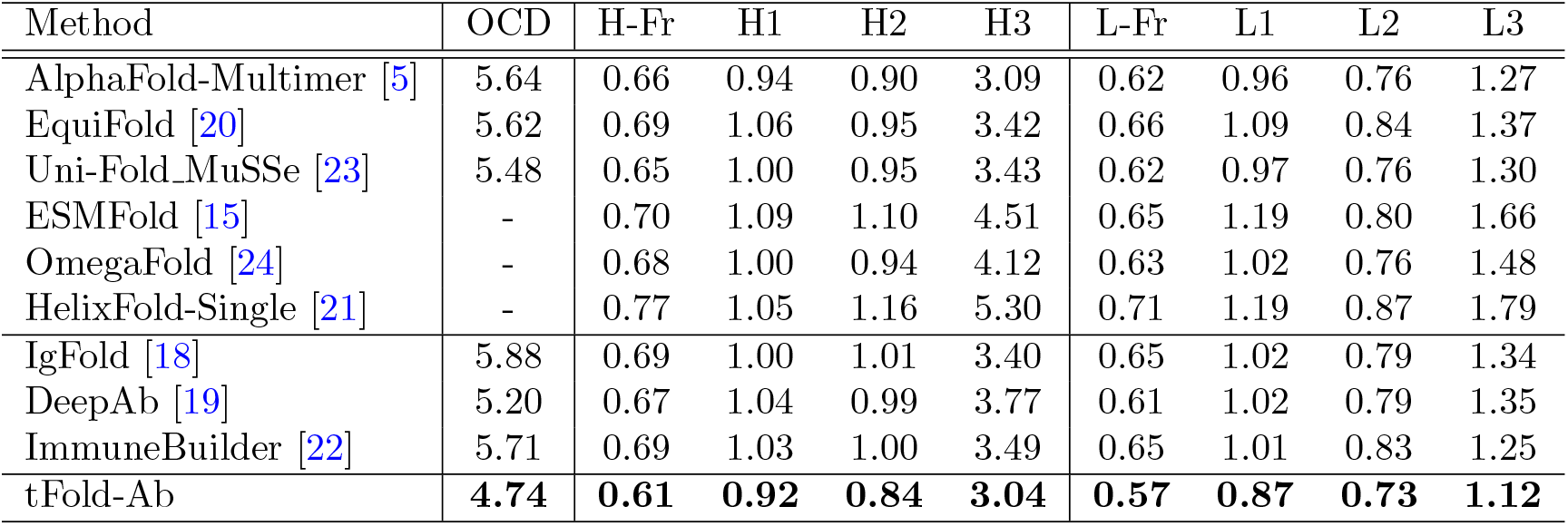
Antibody structure prediction performance on the SAbDab-22H2-Ab benchmark. OCD, backbone RMSD in different framework and CDR regions are reported. For monomer structure prediction methods, the heavy and light chains are predicted separately, and the OCD metric is not evaluated (denoted by ‘-’). The antibody numbering scheme we use is Chothia. H-Fr indicates the Fr of H chain and H1-H3 indicate the CDRs of H chain. L-Fr indicates the Fr of L chain and L1-L3 indicate the CDRs of L chain.

**Table B3:** DockQ evalution performance on the SAbDab-22H2-Ab benchmark.

| Method | DockQ ↑ | Fnat ↑ | LRMS ↓ | iRMS ↓ |
| --- | --- | --- | --- | --- |
| DeepAb [19] | 0.718 | 0.708 | 2.293 | 1.622 |
| EquiFold [20] | 0.746 | 0.752 | 2.248 | 1.521 |
| IgFold [18] | 0.711 | 0.689 | 2.332 | 1.599 |
| ImmuneBuilder [22] | 0.745 | 0.749 | 2.133 | 1.508 |
| Uni-Fold_MuSSe [23] | 0.752 | 0.771 | 2.199 | 1.515 |
| AlphaFold-Multimer [5] | **0.769** | **0.786** | 2.077 | 1.416 |
| tFold-Ab | 0.766 | 0.746 | **1.889** | **1.339** |

**Table B4:**
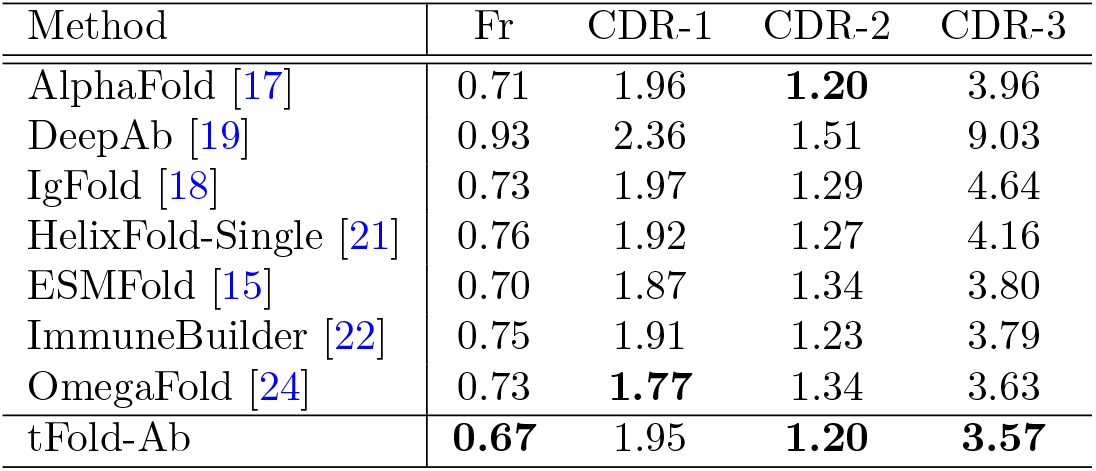
Nanobody structure prediction performance on the SAbDab-22H2-Nano benchmark. Backbone RMSD in different framework and CDR regions are reported. The antibody numbering scheme we use is Chothia.

**Table B5:** Antibody-antigen complex prediction performance on the SAbDab-22H2-AbAg benchmark. SR denotes DockQ success rate defined by DockQ algorithm. TM-score denotes the accuracy of the prediction in comparison to the ground truth structure, with a range from 0 to 1 and a threshold of 0.5 denoting the correct prediction. *R*^2^ is correlation of determination between the confidence score and DockQ.

| Method | Extra Features | DockQ ↑ | SR ↑ | TM-Score ↑ | $R^2$ ↑ |
| --- | --- | --- | --- | --- | --- |
| AlphaFold-Multimer [5] (v2.3) | - | 0.158 | 0.182 | 0.665 | 0.600 |
| AlphaFold-Multimer (v2.2) | - | 0.108 | 0.121 | 0.649 | 0.375 |
| Uni-Fold_MuSSe [23] | - | 0.048 | 0.030 | 0.590 | - |
| RoseTTAFold2 [28] | - | 0.030 | 0.000 | 0.501 | - |
| ZDock [9] | - | 0.031 | 0.000 | 0.599 | - |
| ClusPro [8] | - | 0.038 | 0.000 | 0.607 | - |
| HDock [10] | - | 0.034 | 0.020 | 0.607 | - |
| EquiDock [11] | - | 0.004 | 0.000 | 0.586 | - |
| DyMEAN [12] | epitope | 0.160 | 0.283 | 0.700 | - |
| ColabDock [13] | contact | 0.266 | 0.414 | 0.722 | 0.387 |
| tFold-Ag | - | **0.217** | **0.283** | **0.708** | 0.593 |
| tFold-Ag-ppi | epitope | 0.303 | 0.465 | 0.761 | 0.346 |
| tFold-Ag-ppi | epitope and paratope | 0.416 | 0.626 | 0.814 | 0.407 |
| tFold-Ag-contact | contact | 0.703 | 0.970 | 0.918 | 0.561 |

**Table B6:** Nanobody-antigen complex prediction performance on the SAbDab-22H2-NanoAg benchmark. SR denotes DockQ success rate defined by DockQ algorithm. TM-score denotes the accuracy of the prediction in comparison to the ground truth structure, with a range from 0 to 1 and a threshold of 0.5 denoting the correct prediction. *R*^2^ is correlation of determination between the confidence score and DockQ.

| Method | Extra Features | DockQ ↑ | SR ↑ | TM-Score ↑ | $R^2$ ↑ |
| --- | --- | --- | --- | --- | --- |
| AlphaFold-Multimer [5] (v2.3) | - | **0.149** | 0.140 | **0.692** | 0.545 |
| AlphaFold-Multimer (v2.2) | - | 0.113 | 0.116 | 0.680 | 0.241 |
| Uni-Fold MuSSe [23] | - | 0.048 | 0.047 | 0.590 | - |
| RoseTTAFold2 [28] | - | 0.022 | 0.000 | 0.501 | - |
| ZDock [9] | - | 0.028 | 0.000 | 0.599 | - |
| ClusPro [8] | - | 0.026 | 0.000 | 0.607 | - |
| HDock [10] | - | 0.029 | 0.000 | 0.607 | - |
| EquiDock [11] | - | 0.010 | 0.000 | 0.586 | - |
| ColabDock [13] | contact | 0.267 | 0.419 | 0.760 | 0.429 |
| tFold-Ag | - | 0.136 | **0.186** | 0.678 | 0.237 |
| tFold-Ag-ppi | epitope | 0.227 | 0.349 | 0.733 | 0.182 |
| tFold-Ag-ppi | epitope and paratope | 0.316 | 0.628 | 0.761 | 0.229 |
| tFold-Ag-contact | contact | 0.731 | 1.000 | 0.918 | 0.595 |

**Table B7:** Performance of virtual screening for binding antibodies on a set of 745 antibodies targeting PD-1, including 27 positive samples. Our method yielded 2 positive samples within the top 1% of the ranking, resulting in an *EF* (1%) of 7.41.

| rank | antibody_id | label | score |
| --- | --- | --- | --- |
| 1 | Lipustobart | 1 | 0.883 |
| 2 | Healthy_donor_antibody | 0 | 0.867 |
| 3 | Cemiplimab | 1 | 0.861 |
| 4 | Healthy_donor_antibody | 0 | 0.860 |
| 5 | Healthy_donor_antibody | 0 | 0.859 |
| 6 | Healthy_donor_antibody | 0 | 0.858 |
| 7 | Healthy_donor_antibody | 0 | 0.855 |

**Table B8:** CDRs loop deisgn performance on the SAbDab-22-DesingAb benchmark.

| Method | Extra Features | H1 |  | H2 |  | H3 |  |
| --- | --- | --- | --- | --- | --- | --- | --- |
|  |  | AAR | CAAR | AAR | CAAR | AAR | CAAR |
| DiffAb[34]<sup>a</sup> | - | 0.686 | 0.545 | 0.395 | 0.322 | 0.290 | 0.190 |
| RefineGNN[37]<sup>ab</sup> | - | - | - | - | - | 0.251 | - |
| MEAN[35]<sup>a</sup> | - | - | - | - | - | 0.311 | 0.243 |
| HERN[36] | epitope | - | - | - | - | 0.310 | 0.141 |
| DyMEAN[12]<sup>c</sup> | epitope | 0.731 | 0.615 | 0.596 | 0.519 | 0.310 | 0.244 |
| ESM-PPI<sup>b</sup> | - | 0.726 | - | 0.427 | - | 0.240 | - |
| tFold-Ag | - | 0.651 | 0.531 | 0.422 | 0.331 | 0.309 | 0.212 |
| tFold-Ag-ppi | epitope | 0.696 | 0.550 | 0.509 | 0.390 | 0.347 | 0.251 |
| tFold-Ag-ppi | epitope and paratope | 0.691 | 0.551 | 0.524 | 0.400 | 0.358 | 0.265 |

<sup>a</sup>Adope the pipline: Antibody use ground truth, Antigen use AlphaFold2, dock with HDock[10]
<sup>b</sup>Does not use antigen structure as input, so there is no need to calculate CAAR
<sup>c</sup>More than 70% of the test set in SAbDab-22-Design was in the training set for dyMean. This overlap could potentially lead to an overestimation of its performance

**Table B9:** CDRs loop deisgn performance on the Cov-AbDab-DesignAb benchmark.

| Method | H1 | H2 | H3 |
| --- | --- | --- | --- |
|  | AAR | AAR | AAR |
| DiffAb [34]<sup>a</sup> | 0.634 | 0.331 | 0.239 |
| RefineGNN [37]<sup>a</sup> | - | - | 0.232 |
| MEAN [35]<sup>a</sup> | - | - | 0.261 |
| ESM-PPI | **0.690** | 0.370 | 0.192 |
| tFold-Ag | 0.532 | **0.416** | **0.267** |

<sup>a</sup>Adope the pipline: Antibody use tFold-Ab, Antigen use AlphaFold2, dock with HDock[10]

**Table B10:**
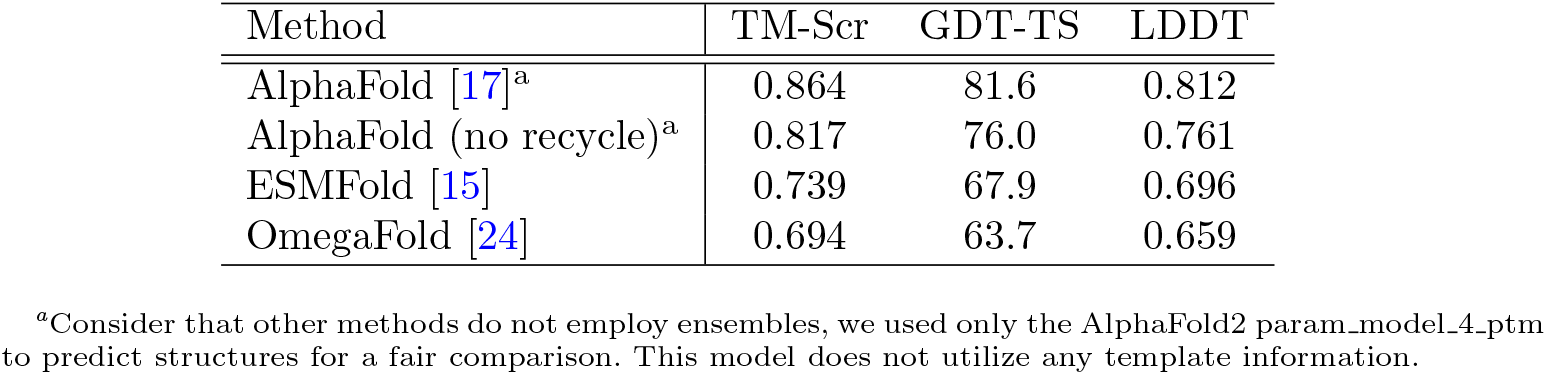
Antigen structure prediction performance on the SAbDab-22-Ag benchmark. We have collected all antigen sequences and structures released in the SAbDab database in 2022. Using mmseqs with 40% sequence consistency for clustering, we obtained a total of 112 sequences. We then compared the performance of the currently leading MSA-based structure prediction algorithm, AlphaFold2, with famous single-sequence-based structure prediction methods, ESMFold and OmegaFold, on this test set. We also evaluated the performance of alphafold2 when the recycle strategy was disabled.

**Table B11:** Antibody structure prediction performance on SAbDab-22H2-AbAg when antigen is consider or not. Backbone RMSD in CDR-H3 are reported. tFold-Ag’s AI-based docking module refine the prediction of CDR-H3 loop.

| Dataset | tFold-Ab | tFold-Ag | tFold-Ag-ppi |
| --- | --- | --- | --- |
| SAbDab-22H2-AbAg | 3.21 | 3.20 | 3.07 |
| SAbDab-22H2-AbAg(DockQ $\geq 0.23$)<sup>a</sup> | 3.10 | 2.75 | 2.54 |

<sup>a</sup>Considering the difficulty in identifying the epitope of the antigen and the paratope of the antibody, in order to correctly evaluate the simultaneous modeling of antibody and antigen structures by tFold-Ag, compared to tFold-Ab’s independent prediction of antibodies, we want to see if it can improve the predictive performance of the antibody CDR-H3. We selected 28 successful antigen-antibody pairs predicted by tFold-Ag from SAbDab-22H2-AbAg (DockQ $\geq 0.23$ ). For these data points, tFold-Ag can correctly predict the antigen epitope without using prior epitope information.

## Appendix C Supplementary Methods

### C.1 Overview

In this section, we provide a detailed account of our methods and results. We begin with an introduction to the notations and conventions used throughout our work in the Section C.2. Following this, we present the data used by our pre-trained language model along with the detail of the training process in Section C.3. Subsequently, we introduce the algorithms, loss functions, training details and additional results of tFold-Ab and tFold-Ag that are introduced in the main text in Section C.4 and Section C.5. Finally, we showcase some of our research on two applications of our models, structure-based virtual screening and antibody design in Section C.6 and Section C.7. These applications demonstrate the practical utility of our models and their potential to contribute significantly to the field of bioinformatics.

### C.2 Notation

We summarize all algorithms in Table C12, covering the main algorithms of this paper, tFold-Ab and tFold-Ag as well as the algorithms that these two algorithms call. In detail, we provide hyperlinks to the 12 algorithms proposed in this paper so that the reader can learn their details. For the remaining algorithms proposed by existing works (e.g., AlphaFold2), we provide the algorithm names and references so that readers can refer to the literature for details. For some well-known algorithms, e.g., ‘Linear’, we provide their synopses.

In Table C13, we provide a brief introduction to symbols and operators used in the algorithms and equations.

**Table C12:**
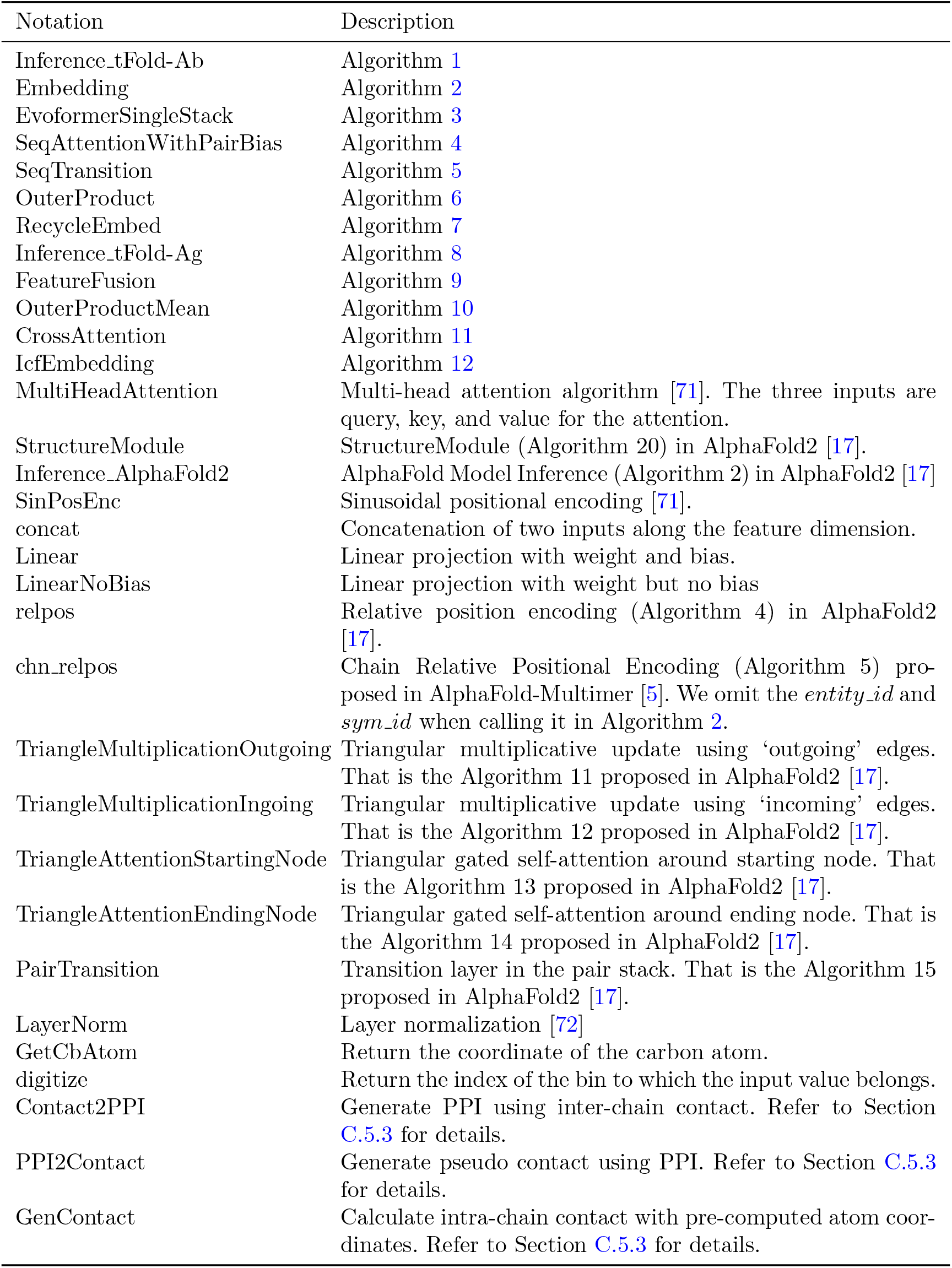
Summary of algorithms used in this paper.

**Table C13:** Summary of notations used in this paper.

| Notation | Description |
| --- | --- |
| Indices $i, j$ | Indices on the residue dimension |
| Indices $k, l$ | Indices on the antibody dimension |
| Indices $s, t$ | Indices the sequence dimension |
| Index $h$ | Index on the attention heads dimension |
| Superscripts H, L, A | Superscripts H, and L represent the heavy chain and light chain of antibody, respectively. Superscript A represents the antigen. Superscript H-L represents the concatenation of heavy and light chains. Superscript H-L-A represents the concatenation of antibody and antigen. |
| Superscripts ab, ag | Superscripts ab and ag represent antibody and antigen, respectively. They are mainly used in Algorithm 9: Feature Fusion. In Algorithm 9, the superscript ab also represents nanobody if applicable. |
| Superscript pre, init, prev | Superscripts ‘pre’, ‘init’, and ‘prev’ represent ‘pre-trained’, ‘initial’, and ‘previous’, respectively. |
| Superscript plddt, iptm | Superscripts ‘plddt’, ‘iptm’ in $r_i^{\text{plddt}}$ and $r^{\text{iptm}}$ are used to identify pLDDT and ipTM confidence scores. |
| $\mathbf{f}_i$ | Feature. $\mathbf{f}_i^{\text{H}}$, $\mathbf{f}_i^{\text{L}}$ are the sequence feature of heavy and light chain of antibody. $\mathbf{f}_i^{\text{A}}$ is the MSA feature of antigen. |
| $\mathbf{s}_i$ | Single representation. |
| $\mathbf{z}_{ij}$ | Pair representation. |
| $\mathbf{m}_{si}$ | MSA representation. |
| $\mathbf{x}_i$ | Structure representation, i.e. the coordinates of each atom in the protein. |
| $r$ | Confidence score. $r_i^{\text{plddt}}$ is the pLDDT confidence score for each residue and $r^{\text{iptm}}$ is the ipTM confidence score for each antibody. |
| $\mathbf{F}_i^s$ | PPI feature. |
| $\mathbf{F}_{ij}^z$ | Inter-chain feature. |
| $\{\cdot\}$ | Set of vectors. For example, $\{\mathbf{s}_i\}$ denotes the set of single represents. |

### C.3 ESM-PPI

#### C.3.1 Dataset

A combination of four datasets is used in the further pre-training for ESM-PPI:

- **UniRef50** [16], March 2023 version, approximately 60 million monomeric sequences in it. To ensure the validity of our validation set, we randomly selected 4,000 sequences that have never appeared in the training and validation sets of ESM2. These sequences were obtained by taking the difference set between UniRef50 and UniRef100 released in September 2021, and were used to construct the validation and test set.
- **PDB**, January 2023 version, a total of 188k of sequences of solved protein multimers in the Protein Data Bank. We selected 4,000 pairs of interacting chains from 4,000 complex structures that were released after 2022 from the PDB to serve as the validation and test set.
- **PPI**, the dataset compiles 1.3 million pairs of protein sequences that are known to potentially interact, gathered from various existing databases. we amalgamated pairs of interacting proteins from the following sources: HINT [73], intACT [74], HIPPIE [75], prePPI [76], BioGRID [77], comPPI [78] and huMAP [79]. Given that these data come from various sources, redundancy is inevitable. To address this, we employed MMseqs2 [80] to filter out duplicated PPI based on 100 sequence identities. In addition, we filter the interactions with low confidence scores in intACT (score lower than 0.3), comPPI (score lower than 0.3), HIPPIE (quality lower than 0.63) for higher data quality. After that, we select 4,000 pairs for validation and test.
- **Antibody**, This dataset includes 1.5 million paired antibody sequences collected from the OAS [40]. Considering that the antibody’s CDR3 region can be affected by the antigen, and the OAS data does not take it into account, we did not select any paired antibody as a validation or test set.

Ultimately, we obtained a total of 12,000 monomeric sequences and interaction pairs, which were designated for validation and testing. These were evenly split, with half being selected as the validation set and the remaining half assigned to the test set. This balanced division ensures a comprehensive and rigorous assessment of our model’s performance.

In order to partially prevent overfitting and to avoid data leakage, we have reduced the redundancy in the training data. For monomeric sequence in UniRef50, we removed some sequences from the training set via the procedure described in [81]. MMseqs2 search (*-min-seq-id 0*.*5 -alignment-mode 3 -max-seqs 1000 -s 7 -c 0*.*8 -cov-mode 0* ) is run using the training set as query database and the validation & test set (including monomeric sequences and each single chain from multimer) as target database. All train sequences that match a validation & test sequence with 50% sequence identity under this search are removed from the training set. For multimeric sequence from PPI and PDB, we calculate the length-weighted sequence similarity using MMseqs2 search (*-min-seq-id 0*.*9 -alignment-mode 3 -max-seqs 1000 -s 7 -c 0*.*8 -cov-mode 0* ). Considering the small number of PPI and PDB data, we only chose 90% identity threshold for multimer.

#### C.3.2 Training detail of ESM-PPI

To optimize the training of ESM-PPI for handling the large number of residues in multimeric sequences, which can be a challenge for GPU memory and efficiency, we adapted the *ContiguousCropping* algorithm from AlphaFold-Multimer [5]. Specifically, for multimeric structures from the PDB, we randomly select two interacting chains and proportionally crop them according to their lengths, ensuring the preservation of structural contacts. In the absence of structural data for PPI sequence pairs, cropping is based solely on length. For homomers, we employ a uniform cropping method across all chains to maintain consistency. Antibody sequence pairs, typically shorter, do not require this cropping step.

For monomeric sequences and PPI pairs without structural data, we implement a masking strategy akin to that used in traditional masked language models. During training, 15% of amino acids are randomly chosen: 80% of these are masked, 10% are substituted with a random residue, and 10% are left as is. To avoid data leakage in homomers, we ensure that the masking is uniform across all chains.

When dealing with protein sequences that have experimental structures, we increase the likelihood to 30% that residues at the contact interface between two chains are masked. This approach improves the model’s proficiency in learning protein interactions from sequence data. For antibodies, given the predictability of amino acid types in the framework regions, we adopt a strategy where residues in the CDRs loop have a 30% higher chance of being selected for masking, emulating tactics used in specialized antibody language models [45], while the framework regions remain unchanged.

In the further pre-training phase, we executed a comprehensive 128,000 training steps, with each step comprising a batch of 128 samples. These samples varied, including individual sequences from UniRef50, pairs from multimeric PDB structures, PPI interaction pairs, and paired antibody sequences. The selection probability for these sample types followed a ratio of 2:3:3:2.

The AdamW [82] optimizer was employed with hyperparameters set to *β*_1_ = 0.9, *β*_2_ = 0.999. We adopted a learning rate schedule that began with a linear warm-up from 3e*−*6 to 3e*−*5 over the initial 12,800 steps, after which the rate was held constant at 3e*−*5 for the remainder of the training period.

To efficiently manage the model across multiple GPUs, we utilized Fully Sharded Data Parallelism (FSDP) [83], which shards both model weights and optimization states. This enabled us to increase the sequence crop size to 1024. For the validation and testing stages, we applied an Exponential Moving Average (EMA) [84] to the model parameters with a decay rate of 0.995, ensuring smoother parameter updates. The model selection was based on achieving the lowest average loss on a combined validation subset.

The entire further pre-training was conducted on a cluster of 8 NVIDIA A100 GPUs, completing in around 30 hours. This setup allowed us to efficiently scale our training process while managing computational resources effectively.

**Fig. C2:**
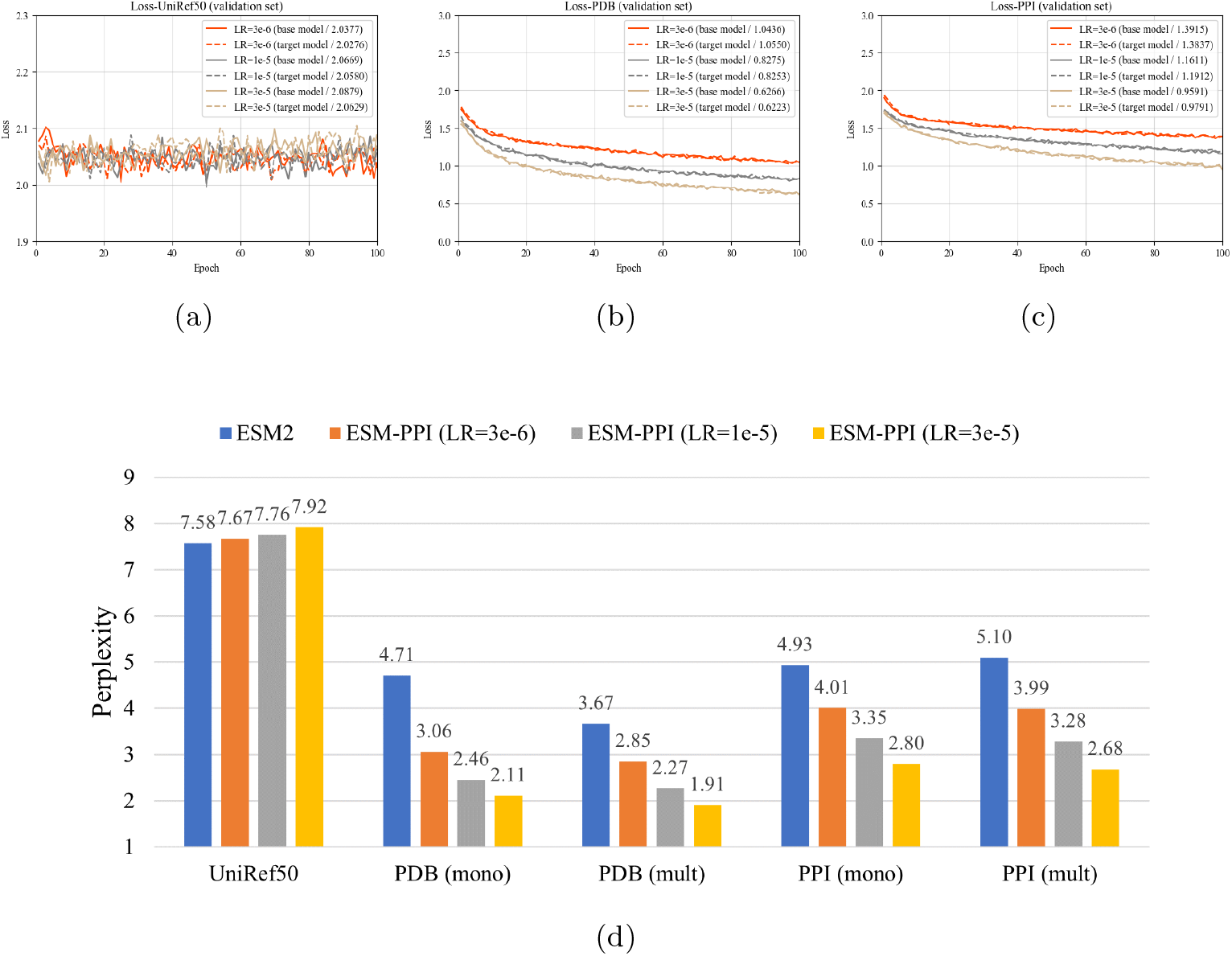
Analysis of ESM-PPI. (a-c) ESM-PPI masked language modeling loss curves on three validation sets. Models are trained to 128k updates. The curves in different colors represent the model trained using different learning rates. ‘base’ refers to the original model, while ‘target’ refers to the model after the parameters have been updated using EMA. (d) ESM-PPI perplexity in different test subsets with respect to varying learning rates. ‘mono’ and ‘mult’ refer to the evaluation of paired sequences either as monomer proteins or as a multimer during assessment.

#### C.3.3 Validation & test analysis

Fig. C2 shows the ESM-PPI validation loss curves on three validation sets. Since ESM-2 has already been well-trained on monomers using UniRef50, the further pretraining stage did not yield any performance improvement. However, for datasets with multimer, the loss gradually decreased with model updates and converged after 100,000 iterations. A learning rate of 3e*−*5 proved to be the most suitable for ESM-PPI. Furthermore, the model updated using EMA showed a slight performance improvement compared to the original model, although the difference was not significant.

We evaluate the performance of ESM-PPI on the test set using perplexity, a metric that measures the model’s ability to predict amino acids based on their context within a sequence. The perplexity score can range from 1, which would indicate a perfect model, to 20, which would suggest a model making random predictions. Essentially, perplexity provides insight into the number of amino acids the model is uncertain about during prediction. In addition, we report perplexity scores for multimer sequences in the ‘mono’ and ‘mult’ modes, respectively. The major difference between these two modes lies in that when recovering the sequence of the first masked chain, whether the second chain is taken as inputs (‘mult’ mode) or not (‘mono’ mode). The reduction in perplexity scores from the ‘mono’ to ‘mult’ mode reflects how well the ESM-PPI model captures cross-chain dependencies for sequence recovery. For the original ESM2 model, which does not accept multimer inputs, we concatenate two sequences with a short linker to evaluate its performance in the ‘mult’ mode.

Fig. C2d indeed shows that compared to ESM2, the perplexity of ESM-PPI on UniRef50 has decreased, indicating an improvement in the model’s predictive ability. When PPI and PDB data are input into the model as monomeric proteins, the performance of ESM-PPI markedly outperforms that of ESM2. This can be attributed to the fact that multimeric protein data is less abundant and diverse than monomeric protein data, making it easier for the model to learn the individual features of the protein, rather than the cross-chain features. When the entire protein complex is used as input, the model’s perplexity decreases relative to when only a monomeric chain is inputted. For example, the perplexity for the PDB validation set drops from 2.11 to 1.91. This suggests that ESM-PPI has successfully learned to interpret cross-chain information, and is not merely relying on the individual chain’s information for amino acids prediction. This showcases the model’s capability to comprehend and predict complex protein interactions.

Fig. C2d indeed shows that the perplexity of ESM-PPI on UniRef50 has slightly increased over ESM2, indicating minor performance degradation in the model’s ability of recovering monomer sequences. This is as expected since ESM2 is solely pretrained on UniRef50, while ESM-PPI is jointly optimized on UniRef50, PDB, and PPI datasets. As for multimer sequences in PDB and PPI validation sets, the performance of ESM-PPI markedly outperforms that of ESM2 in both ‘mono’ and ‘mult’ modes. For ESM-PPI models, the performance gain in the ‘mono’ mode can be partially explained by the over-sampling of PDB and PPI sequences than their original proportion in UniRef50. However, all the ESM-PPI models consistently achieve a lower perplexity in the ‘mult’ mode than that in the ‘mono’ mode, implying that these models indeed exploit cross-chain information to more accurate sequence recovery. It is worth noting that simple concatenation of multimer sequences does not always yields a lower perplexity. For the PPI validation set, ESM2 with concatenated sequences leads to an increased perplexity (from 4.93 to 5.10), in contrast with ESM-PPI models. In summary, this showcases the model’s capability to comprehend and predict complex protein interactions.

### C.4 tFold-Ab

#### C.4.1 tFold-Ab inference

Algorithm 1 shows the inference pipeline of tFold-Ab. The tFold-Ab takes the antibody’s amino-acid sequence as input and generates outputs that include sequence embedding, pairwise representations, atom coordinates and confidence scores.

The entire network is primarily composed of four components. The first part of the network initializes antibody feature using pre-trained ESM-PPI (refer to Appendix section C.3 for details.). This is followed by the Embedding Module (Appendix section C.4.2), which generates the initial sequence embedding. The sequence embedding is then updated iteratively with the second part of the network, Evoformer-Single stack (Appendix section C.4.3), creating the inputs for the structure module and recycling. The structure module uses invariant point attention (IPA) [17] for structure prediction, which is the third part of the network. Finally, the outputs from the previous execution are recycled as inputs for the next execution for additional refinement (Appendix section C.4.4). This recycling process forms the fourth and final part of the network. This iterative refinement process helps to improve the accuracy of the predicted structures.

##### Algorithm 1

Model inference with recycling iterations for tFold-Ab

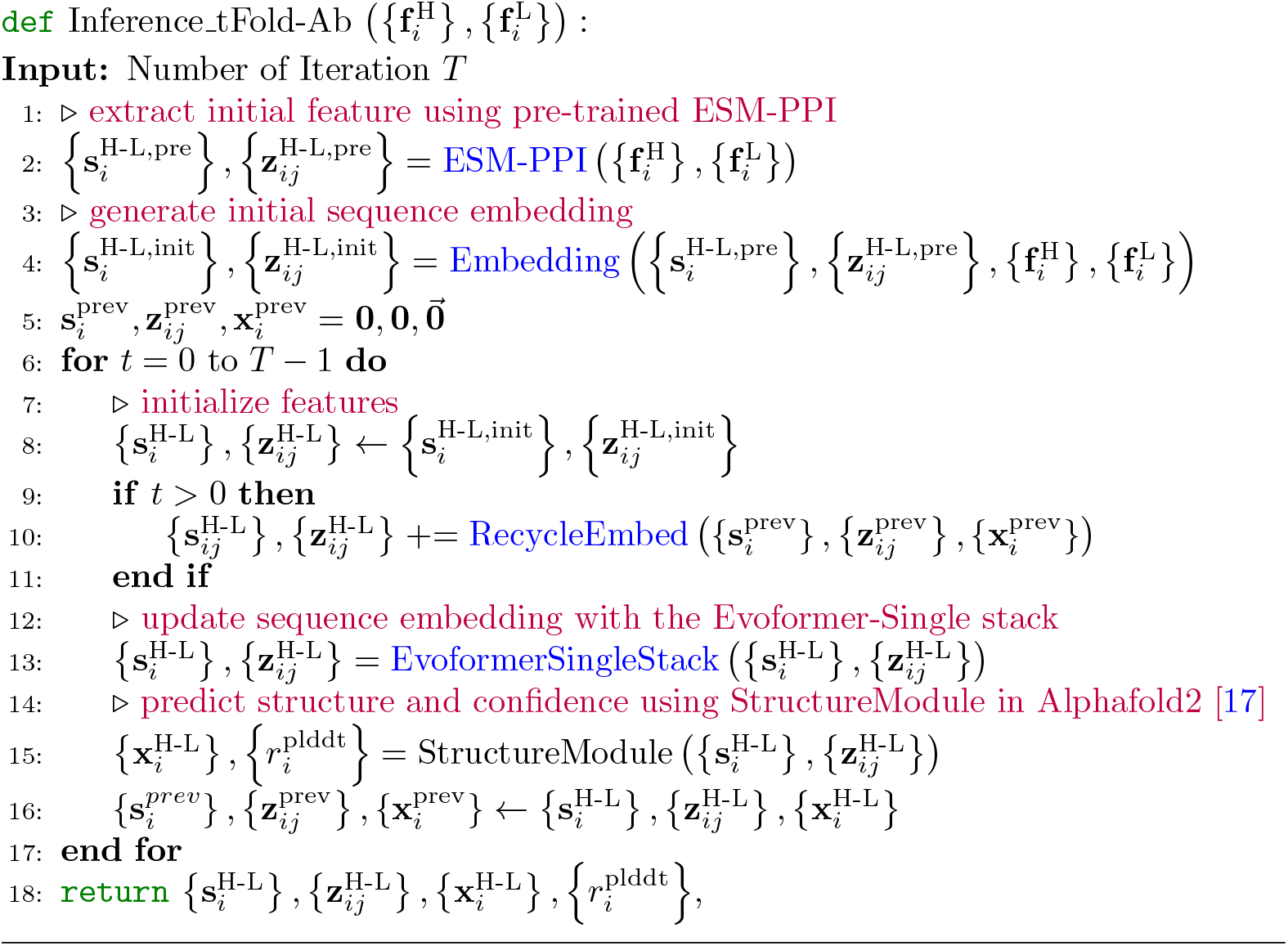

#### C.4.2 Embedding module

Algorithm 2 illustrates the use of three distinct positional encoders. These encoders not only provide the network with information about the positions of residues in the chain, but also help distinguishing between heavy and light chains in antibodies, which is crucial for accurate protein folding prediction.

The multimer positional encoder uses a shared sinusoidal positional encoder for different chains. This type of encoder helps the model understand the relative positions of residues within each chain. The relative positional encoder is a version adapted from AlphaFold2. This encoder provides information about the relative positions of residues to each other, which is important for understanding the 3D structure of the protein. The chain relative positional encoder is a simplified version of the chain relative positional encoding used by AlphaFold-Multimer. This encoder provides information about the relative positions of different chains to each other.

Given the differences between antibody heavy and light chains, we have omitted the *entity id* and *sym id*, which were previously used by AlphaFold-Multimer. These identifiers were used to distinguish between different entities and symmetries in the protein structure, but in the context of antibodies, distinguishing between heavy and light chains is more relevant.

##### Algorithm 2

Embedding Module for tFold-Ab

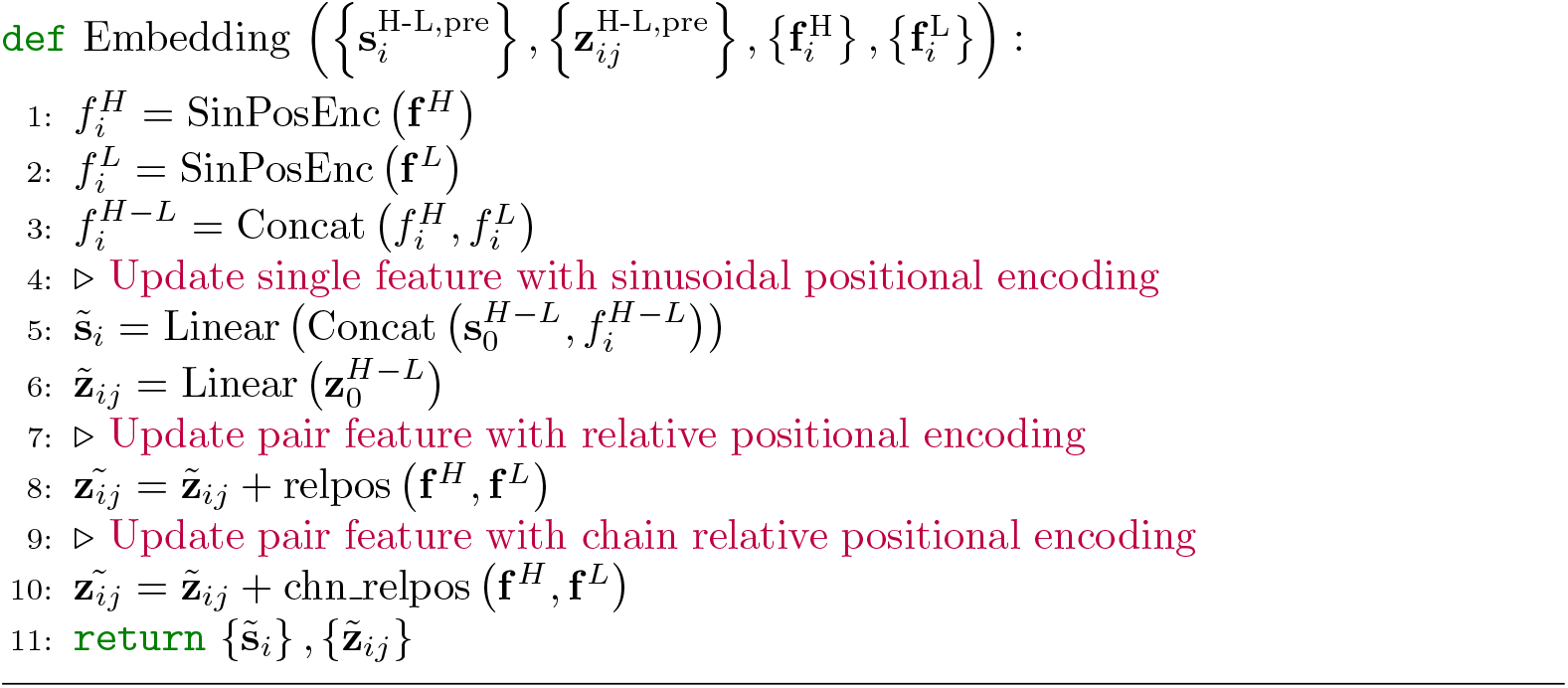

#### C.4.3 Evoformer-Single stack

As mentioned earlier, our method only takes the amino-acid sequence itself as inputs, without any sequence homologs. Therefore, we simplify AlphaFold’s Evoformer stack [17] (originally designed for updating MSA and pair features) to handle single sequence inputs for iterative updates of single and pair features, as illustrated in Algorithm 3.

Specifically, ‘MSAColumnAttention’ in the original Evoformer block is removed since calculating attention weights among sequence homologs is no longer applicable for single sequence inputs. Other MSA-related modules are correspondingly simplified, while modules involving only pair features remain unchanged. It is worth noting that we employ *N* ^block^ = 16 blocks for antibody-only input and *N* ^block^ = 32 blocks for antibodies with antigen.

##### Algorithm 3

Evoformer-Single stack

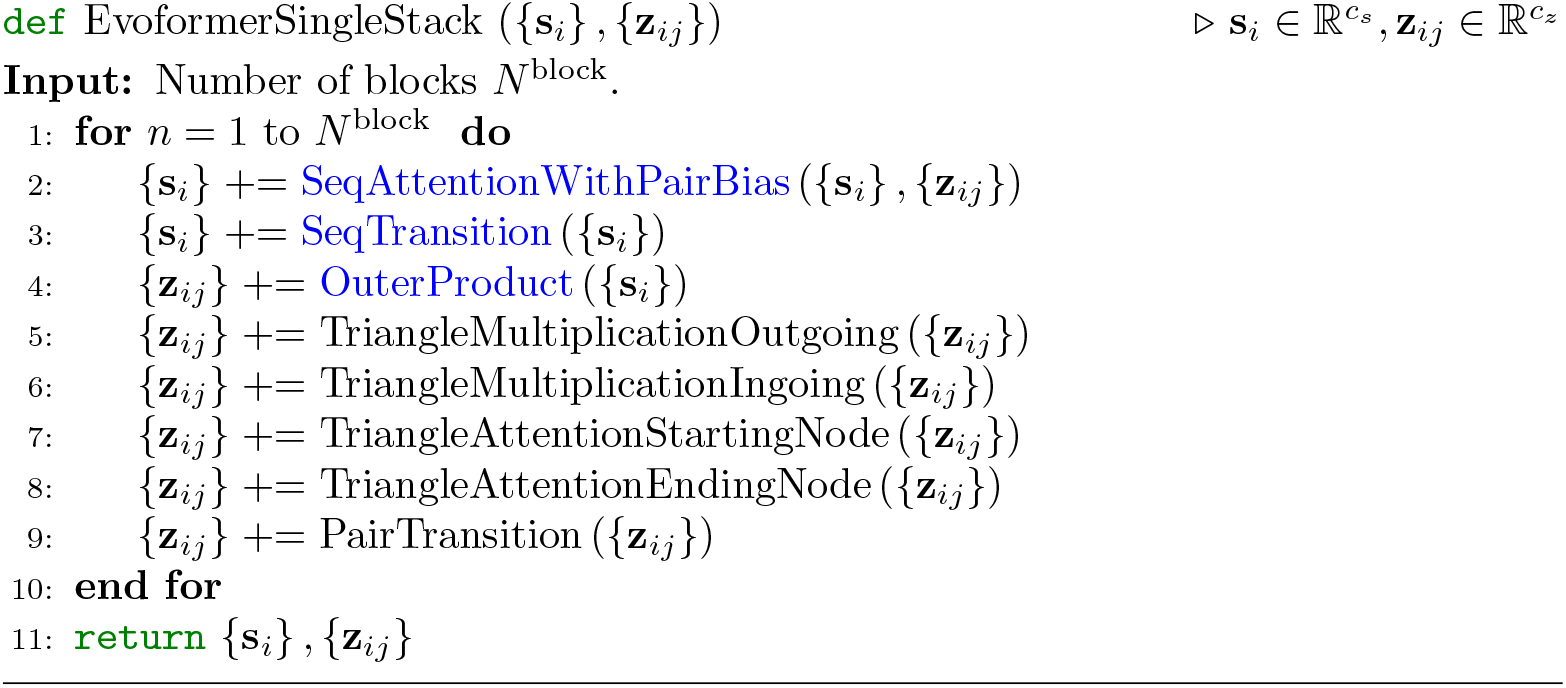

In Algorithm 4, single features are updated via the gated self-attention mechanism along the sequence dimension to formulate the inter-residue interaction. Additionally, pair features are linearly projected into bias terms, which are then imposed onto attention weights. This allows updating single features under the guidance of interresidue pair features.

##### Algorithm 4

Sequence gated self-attention with pair bias

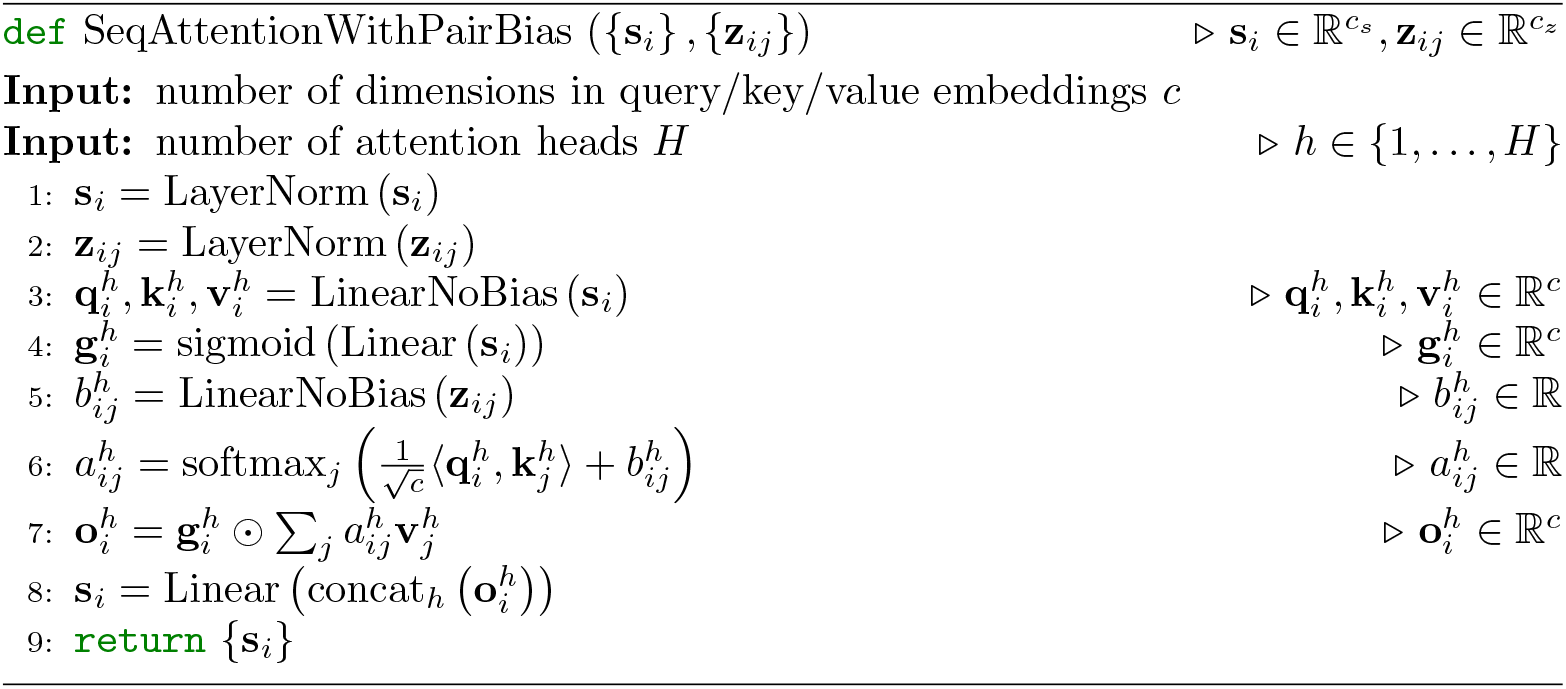

In Algorithm 5, single features are updated via a two-layer feed-forward network. We set the number of dimensions in hidden embeddings as *c* = 4*c*_*s*_, the same as AlphaFold.

Afterwards, pair features are updated based on the outer-product of single features, as described in Algorithm 6.

##### Algorithm 5

Transition layer in the Evoformer-Single stack

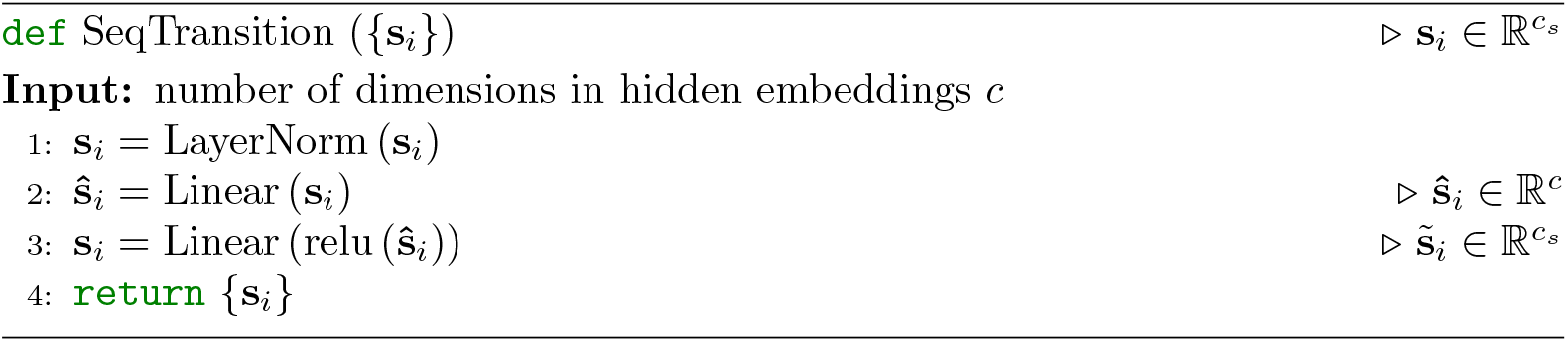

##### Algorithm 6

Outer product

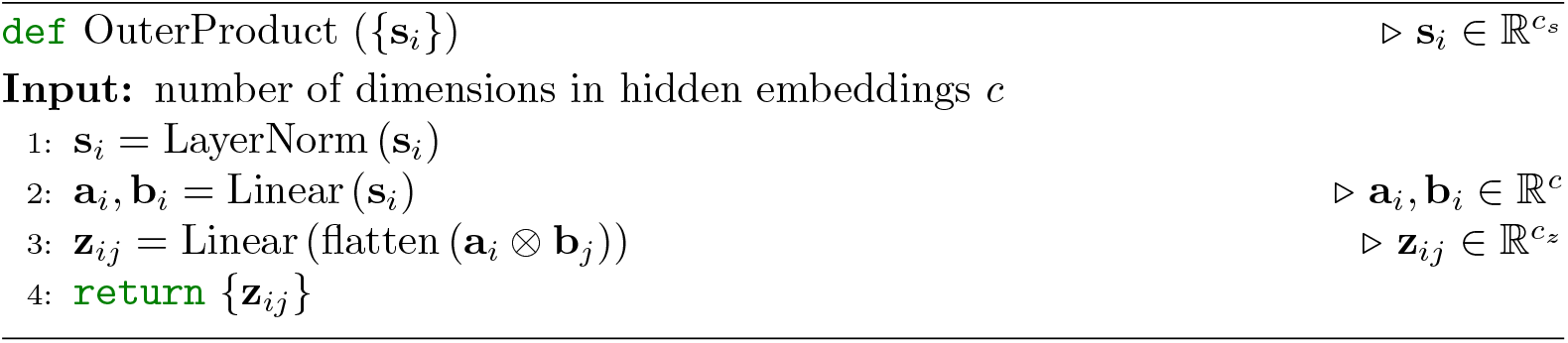

#### C.4.4 Recycling embedding

In order to utilize feature embeddings and structure predictions from the previous iteration (as in Algorithm 1), we slightly modify the original ‘RecyclingEmbedder’ module in AlphaFold2 [17] for single sequence inputs. The modified recycling embeddings module is as described in Algorithm 7.

Specifically, single and pair features from the previous iteration are normalized to produce residue update terms for the current iteration. Atomic coordinates of *C*_*β*_ (*C*_*α*_ for glycines) atoms are extracted from the predicted structure, from which pairwise Euclidean distance is computed. Such distance is then discretized into histogram bins to generate one-hot encodings for the final linear projection.

#### C.4.5 Loss function

The proposed model is designed to be trained with both single-chain and double-chain antibody structures. To balance the weights of the monomer and multimer loss functions, we use a mixture of monomer and multimer structure prediction loss functions by

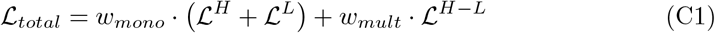

where *ℒ*^*H*^, *ℒ*^*L*^, and *ℒ*^*H−L*^ are loss functions for heavy chain, light chain, and heavy-light chain complex, respectively. We introduce *w*_*mono*_ and *w*_*mult*_ to balance these loss

##### Algorithm 7

RecycleEmbed

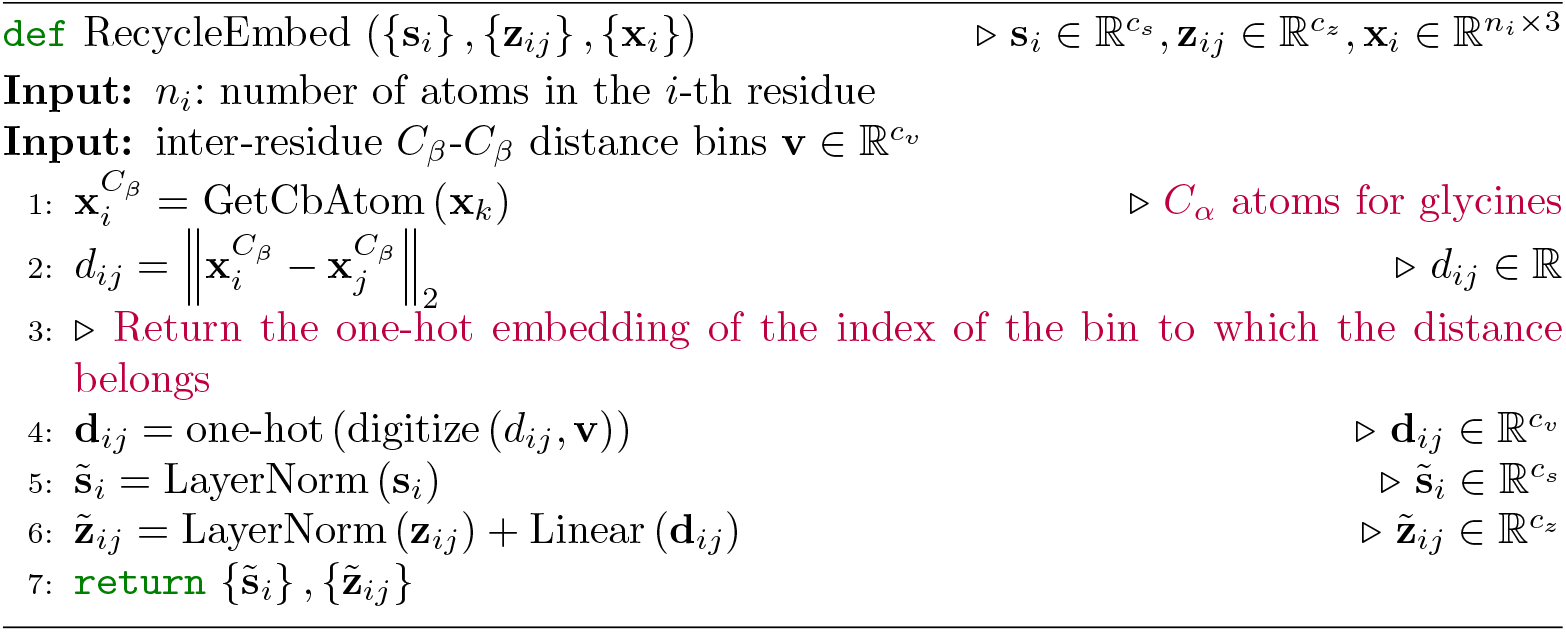

functions: if both chains exist, then *w*_*mono*_ = 0.25 and *w*_*mult*_ = 0.5; otherwise, we have *w*_*mono*_ = 1 and *w*_*mult*_ = 0. This ensures that the model doesn’t overly favor one type of structure over the other, leading to a more balanced and accurate prediction of both heavy and light chain structures.

Each loss term is defined on the predicted monomer/multimer structure and auxiliary predictions (*e*.*g*., inter-residue distance), sharing the same form:

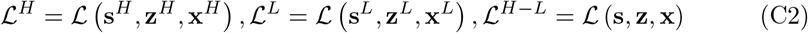

Therefore, we take the multimer loss function *L*^*H−L*^ as an example, and describe its detailed loss terms. Concretely, this loss function constitutes of the following terms:

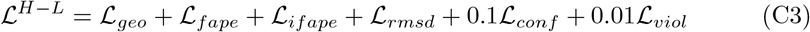

##### Inter-residue geometric loss *ℒ*_*geo*_

To provide more direct supervision in the Evoformer-Single stack, we add four auxiliary heads (implemented as feed-forward layers) on the top of final pair features for predicting inter-residue distance and angles, as defined in trRosetta [85]. This includes:

- 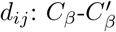distance
- 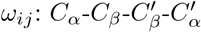 dihedral angle
- 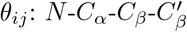 dihedral angle
- 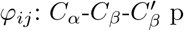 planar angle

Each prediction head outputs probabilistic estimations of the above distance and angles, denoted as 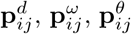, and 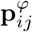. We then calculate the cross-entropy loss for each term and sum them up to as the final inter-residue geometric loss:

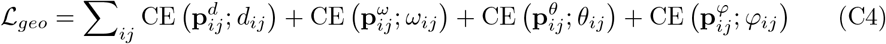

##### Frame aligned point error *ℒ*_*fape*_

This is identical to the FAPE loss used in AlphaFold [17]. After reconstructing full-atom 3D coordinates from per-residue backbone frame and torsion angle predictions, each atom is projected into all the local frames (both backbone and side-chain) in the ground-truth and predicted structures for comparison.

##### Interface frame aligned point error *ℒ*_*ifape*_

This is identical to the second part of FAPE loss used in AlphaFold-Multimer [5], which is applied to inter-chain residue pairs and clamped at 30Å. Please note that this loss is only computed over multimer structure predictions.

##### Coordinate RMSD loss *ℒ*_*rmsd*_

In order to better estimate the overall conformation, we calculate the coordinate RMSD (root-of-mean-squared-deviation) loss between ground-truth and predicted structures after alignment. The optimal alignment is determined by the Kabsch algorithm [86] for finding the optimal rotation and translation between two sets of point clouds. The *ℒ*_*rmsd*_ resulting are penalized with a clamped L1-loss with a length scale *Z* = 10Å to make the loss unitless. For the *i*-th residue’s *j*-th atom, we denote its 3D coordinate in the ground-truth structure as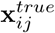, and 3D coordinate in the aligned predicted structures as **x**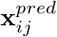. *T* represents SE(3)-transformation from the prediction frame to the ground-truth reference frame. The coordinate RMSD loss is defined as:

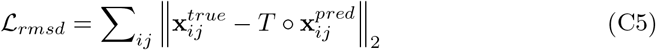

##### Confidence loss *L*_*conf*_

This includes loss functions for pLDDT and pTM predictions, same as AlphaFold [17]. We detach single and pair features before estimating pLDDT and pTM scores, similar to IgFold [18], to prevent the model from generating problematic structures whose lDDT and TM-score can be accurately predicted.

##### Structure violation loss *L*_*viol*_

Similar to AlphaFold [17], we introduce penalty terms for incorrect peptide bond length and angles, as well as steric clashes between non-bonded atoms. For multimer structure prediction, we do not penalize the bond length and angle between the last residue in the heavy chain and the first residue in the light chain, since there is no peptide bond between them. Besides, we normalize the steric clash loss by the number of non-bonded atom pairs in clash to stabilize the model optimization, as suggested in AlphaFold-Multimer [5].

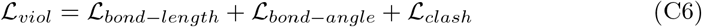

#### C.4.6 Pre-training on PDB

A recent study [87] shows that pre-training on regular protein monomers followed by fine-tuning on antibody monomers can improve the accuracy of antibody prediction. This finding is consistent with the capabilities of our language model, which is adept at extracting features from both monomeric proteins and protein complexes. Given that the antibodies our model predicts may be either single-chain monomers or double-chain complexes, it is reasonable to broaden the scope of our pre-training for the tFold-Ab model to include both regular monomeric proteins and protein complexes. We collected all structures from the PDB released before January 1, 2022. Structures with a resolution lower than 9Å, or those with a proportion of missing residues greater than 0.8 are filtered out. Initially, we separated each chain from every PDB complex and employed MMseqs2 [80] to cluster them with 40% sequence identity. We then mapped each chain back to its corresponding PDB complex based on the clustering result, ensuring that each complex was represented only once within each cluster. This means that the probability of sampling a specific PDB entry is proportional to the aggregate probabilities of the individual chain clusters within the file.

Following the clustering process, we identified approximately 48,000 clusters. From this extensive collection, we selected 1,000 representative families randomly and excluded any remaining families that contained these selected representatives. This resulted in a final training set comprising 45,000 unique clusters, each chosen to maximize the diversity and coverage of the structural space relevant to our model’s training objectives.

During each training epoch, our protocol involves the random selection of a protein complex from each cluster. From the chosen PDB complex, we then randomly select two chains that in contact, defined by having *C*_*α*_ atom pairs within a distance of less than 10Å) to form training samples for the current epoch. If the PDB complex does not contain any interacting chains, a single chain is chosen at random. Additionally, in accordance with methodologies employed in other protein structure prediction research [5, 17], we have compiled a distillation dataset comprising 408,000 structures predicted by AlphaFold2, all with a mean predicted Local Distance Difference Test (pLDDT) score exceeding 70. During the training process, there is an equal probability of selecting a training example from either the distillation set or the experimentally determined structures from the PDB. The ratio of multimeric to monomeric structures in the training set is maintained at 1:1.

Each model was trained on 64 NVIDIA A100 GPUs for approximately 210,000 steps, equivalent to 150 epochs, and was completed within roughly 15 days. The batch size for the training was consistently maintained at 64. The sequence crop size was set to 256 in the first 100 epochs and subsequently increased to 450 for the remaining 50 epochs. When cropping the complexes, we ensured that essential contacts with a distance less than 10 Å between different chains were preserved, even after cropping the protein complexes. This is crucial for accurate modeling of protein-protein interactions. We used Adam [88] as the optimizer with *β*_1_ = 0.9, *β*_2_ = 0.99. The learning rate was warmed up in the first 10 epochs from the initial value of 1e*−*4 to a peak value of 1e*−*3. After the initial 100 epochs of training, we employed two distinct fine-tuning strategies to train two different models. One model without recycling iterations while the other with the number of recycling iterations (*T* ) set to 2.

To evaluate the performance of our tFold-Ab model on regular protein multimers after the pre-training phase, we have meticulously curated a test set, which we refer to as PDB-22H2-Multimer. This dataset comprises a collection of protein structures that were publicly available in the PDB from July 1, 2022, to December 31, 2022. Our selection process was guided by several criteria: Initially, we focused exclusively on complexes that encompassed between 2 to 9 chains. From these, we selected two interacting chains for sampling. Additionally, we ensured that the combined length of these two chains did not exceed 800. We also confirmed that the length-weighted sequence similarity of this two-chain complex, when compared to the training set, was below 40%. Subsequently, we performed clustering based on a 40% sequence identity threshold and selected a representative from each cluster. By adhering to these steps, we successfully compiled 324 complex proteins, including 101 heteromers and 223 homomers, to form our test set.

**Table C14:**
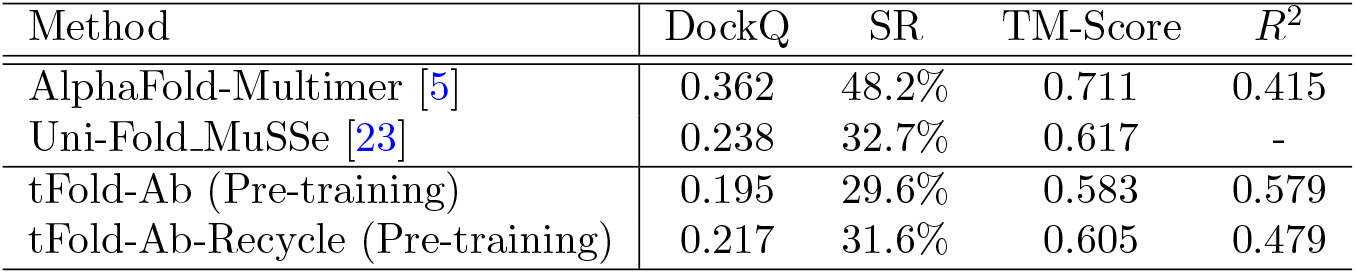
Multimer structure prediction performance on the PDB-22H2-Multimer benchmark. SR denote DockQ success rate defined by DockQ algorithm. TM-score denotes the accuracy of the prediction in comparison to the ground truth structure, with a range from 0 to 1 and a threshold of 0.5 denoting the correct prediction. *R*^2^ is correlation of determination between the confidence score and DockQ.

We compared our pre-training phase tFold-Ab with Uni-Fold MuSSe [23], a method for single-sequence complex prediction, and AlphaFold-Multimer [5], the current state-of-art complex structure prediction method based on MSA and templates, using the PDB-22H2-Multimer dataset. As illustrated in Table C14, AlphaFold-Multimer demonstrates superior performance across all evaluated metrics, attributable to its utilization of paired MSAs that provide ancillary co-evolutionary insights. Our method, tFold-Ab, exhibits a performance marginally inferior to that of Uni-Fold MuSSe. The latter employs a larger language model to extract sequence pair features (using ESM-3B for further pre-training) and utilizes more network model parameters. However, Given that tFold-Ab primarily focuses on antibody structure prediction, it does not require as many parameters. In addition, we found that the recycling process can lead to a minor improvement in the performance of tFold-Ab. However, it also results in a decrease in the correlation between the confidence scores and the prediction performance.

Pre-training on PDB allows the model to learn a broader range of protein features, which could potentially improve its ability to predict the properties and structure of antibodies, especially for CDRs. The importance of this pre-training is confirmed by the ablation studies presented in Appendix C.4.8, which highlight its relative importance in enhancing the model’s predictive capabilities.

#### C.4.7 Training detail of tFold-Ab

We employ the Adam [88] optimizer and set the batch size to 32 for our training process. To initialize our model, we use a model that has been pre-trained for 150 epochs on regular protein complexes from the PDB, as detailed in Appendix C.4.6. For the first 50 epochs, we set the learning rate to 1e*−*3. For the subsequent 150 epochs, we adjust the learning rate to 3e*−*4. Based on the results from our ablation study, which are documented in Appendix C.4.8, we decided not to incorporate additional recycling iterations into our training protocol, as they did not provide a significant improvement in performance. We also maintain the EMA of the model parameters with a decay of 0.999, and we use this model for evaluation. The model that yields the best full-atom RMSD scores on the validation subset is selected as the optimal model.

#### C.4.8 Ablation study of tFold-Ab

We estimate the relative importance of key components of the tFold-Ab architecture by training and evaluating a number of ablation models:

##### Baseline

Baseline model as described in the paper without pre-training using general proteins. All subsequent ablation studies should be interpreted in relation to this foundational model.

##### No ESM-PPI

We replace our ESM-PPI with two other models: the original ESM-2 [15], which has 650M parameters, and ProtTrans-XLNet [56], which has 409M parameters. Given that these two language models are pre-trained on protein monomers and are incapable of handling multi-chain inputs, we follow the approach described in the previous work [15]. We construct an extended antibody sequence where the heavy and light chains are connected by a linker composed of 25 glycines. The embeddings and attention weights from corresponding positions in the antibody are then used to initialize the sequence features and pair features, which serve as the inputs to the model.

##### No one-stage prediction

In our initial version [89], tFold-Ab employed a two-stage architecture. It utilized a language model (*e*.*g*.,ProtTrans-XLNet) to independently predict the structures of the heavy and light chains, followed by a heavy-light chain feature fusion model to integrate the features, which then served as inputs for the multimer structure prediction module. However, given that our language model is capable of extracting features from either single or multiple chains, we have since adopted a one-stage architecture.

##### With pre-training on general protein

Full tFold-Ab model as described in the paper with pre-training using general protein, as described in C.4.6. We also evaluated the performance of the pre-trained initial model without fine-tuning on antibody data, and compared it with the model that was fine-tuned on antibody data.

##### With recycling

employs a recycling strategy during both training and inference phases (refer to Algorithm 7 for details). Specifically, the recycling iterations are set to either 1 or 3 for further refinement. All the recycling models are fine-tuned based on the models that have undergone pre-training on general protein data.

For all ablations, we kept the hyperparameters from the main model configuration, which we have not re-tuned. For each ablation, we trained the model with the same random seeds, all proteins used the same validation set, and we selected the model using the same metrics. Although we would have preferred a larger test set for ablation analysis, the quantity and diversity of antibodies are relatively poor compared to general proteins. Therefore, we used two test sets described earlier, SAbDab-22H2-Ab and SAbDab-22H2-Nano, for ablation analysis.

Ablation results are presented in Table C15 and Table C16. The adoption of a one-stage model architecture simplifies the prediction process of the model and improves performance, reducing the RMSD of CDR-H3 from 3.27 Å to 3.11 Å. Further pre-training of the pre-trained model can improve the performance of antibody structure prediction. Thanks to the use of ESM-PPI, the model can extract features of both monomeric proteins and multimeric proteins, eliminating the need for a glycine linker. The CDR-H3 RMSD of model using ESM-PPI is reduced from 3.30 Å to 3.07 Å compared to the model using ESM2.

**Table C15:** Ablations performance for tFold-Ab on the SAbDab-22H2-Ab benchmark. OCD, backbone RMSD in different framework and CDRs are reported. The antibody numbering scheme we use is Chothia. H-Fr indicates the Fr of H chain and H1-H3 indicate the CDRs of H chain. L-Fr indicates the Fr of L chain and L1-L3 indicate the CDRs of L chain.

| Method | OCD | H-Fr | H1 | H2 | H3 | L-Fr | L1 | L2 | L3 |
| --- | --- | --- | --- | --- | --- | --- | --- | --- | --- |
| baseline | 5.36 | 0.64 | 0.93 | 0.89 | 3.07 | 0.60 | 0.94 | 0.77 | 1.17 |
| -ESM-PPI (ESM2) | 5.18 | 0.64 | 0.97 | 0.90 | 3.30 | 0.59 | 0.93 | 0.75 | 1.19 |
| -ESM-PPI (ProtTrans) | 5.54 | 0.65 | 0.96 | 0.89 | 3.11 | 0.61 | 0.93 | 0.77 | 1.23 |
| -one-stage prediction | 5.55 | 0.70 | 1.07 | 0.97 | 3.27 | 0.66 | 1.03 | 0.79 | 1.35 |
| pre-training only | 6.08 | 0.66 | 1.00 | 0.96 | 3.44 | 0.61 | 1.05 | 0.78 | 1.37 |
| +fine-tuning | 4.74 | 0.61 | 0.92 | 0.84 | 3.04 | 0.57 | 0.87 | 0.73 | 1.12 |
| with Recycling T=2 | 4.99 | 0.61 | 0.94 | 0.84 | 3.03 | 0.57 | 0.89 | 0.73 | 1.10 |
| with Recycling T=4 | 4.88 | 0.60 | 0.93 | 0.83 | 3.03 | 0.57 | 0.87 | 0.72 | 1.11 |

**Table C16:** Ablations performance for tFold-Ab on the SAbDab-22H2-Nano bench-mark. Backbone RMSD in different framework and CDR regions are reported. The antibody numbering scheme we use is Chothia.

| Method | Fr | CDR-1 | CDR-2 | CDR-3 |
| --- | --- | --- | --- | --- |
| baseline | 0.68 | 1.96 | 1.18 | 4.14 |
| -ESM-PPI (ESM2) | 0.70 | 1.98 | 1.21 | 4.55 |
| -ESM-PPI (ProtTrans) | 0.68 | 2.00 | 1.18 | 4.12 |
| -one-stage prediction | 0.77 | 2.02 | 1.31 | 4.35 |
| pre-training only | 0.70 | 1.88 | 1.24 | 4.09 |
| +fine-tuning | 0.67 | 1.95 | 1.20 | 3.57 |
| with Recycling T=1 | 0.66 | 1.90 | 1.16 | 3.64 |
| with Recycling T=3 | 0.67 | 1.86 | 1.18 | 3.61 |

Pretraining on general proteins and then fine-tuning on antibody data greatly improves antibody structure prediction. Although the model pretrained on general proteins does not have good antibody structure prediction performance, after fine-tuning with antibody structure data, the performance in any region of Antibody and Nanobody has been improved. For nanobodies, the improvement is most noticeable, with the RMSD of CDR-3 in SAbDab-22H2-Nano reduced from 4.14 Å to 3.57 Å. This is because the CDR-3 of nanobodies is longer and more flexible, and the generalization of models trained only on antibodies is poor. The use of Recycle can improve the accuracy of Antibody and Nanobody structure prediction, but the improvement is not significant.

Considering that the use of the Recycle strategy will increase additional computational overhead, in tFold-Ag, the tFold-Ab we does not use the Recycle strategy.

#### C.4.9 Inference speed

One of the major advantages of tFold-Ab is its elimination of the time-consuming MSA search procedure. This efficiency is achieved through the integration of pre-trained language models that obviate the need for traditional MSA-based approaches. Additionally, our model formulates both backbone and side-chain conformations with a unified neural network, while previous antibody structure prediction methods, *e*.*g*., DeepAb [19] and IgFold [18], rely on Rosetta-based energy minimization to predict side-chain structures. AlphaFold-Multimer [5] predicts full-atom structures in a single forward pass, its computational complexity, due to the Evoformer stack, is significantly higher than ours.

**Fig. C3:**
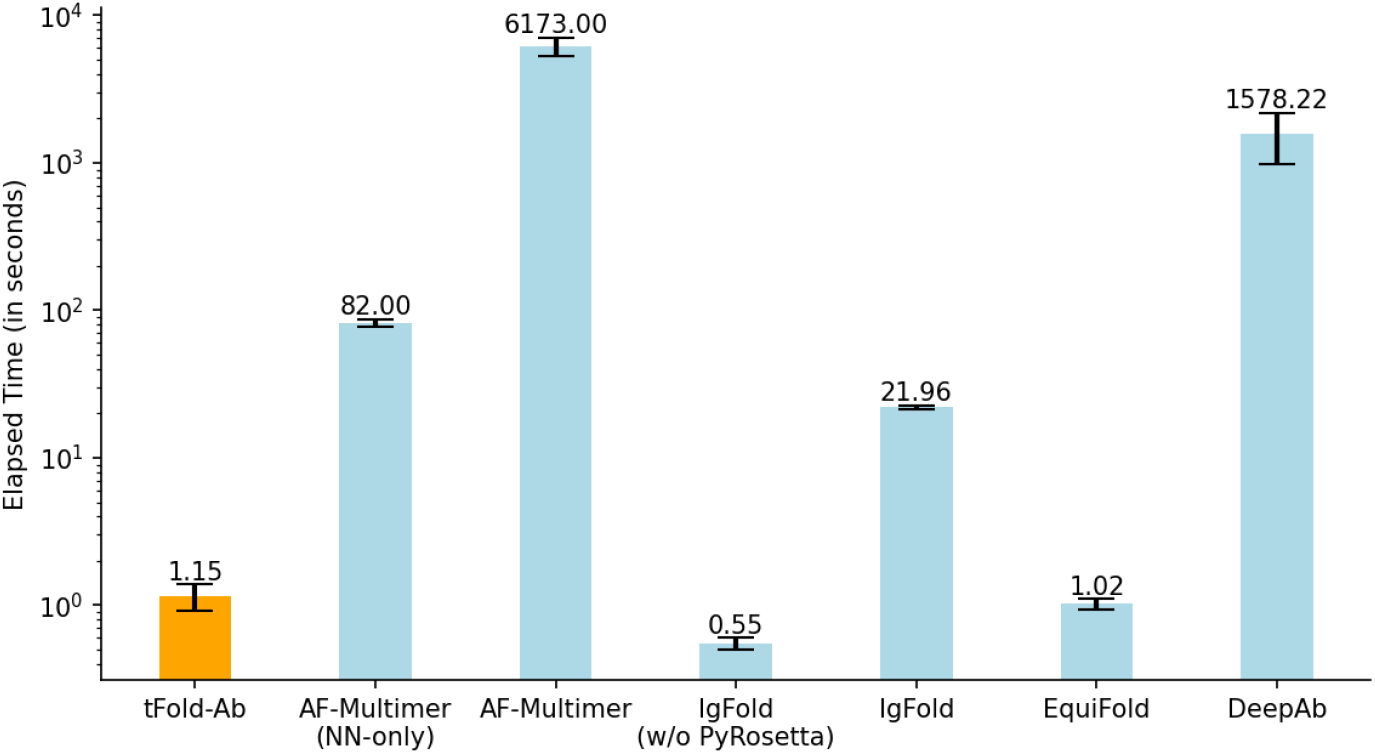
Runtime analysis for tFold-Ab on 170 antibodies in SAbDab-22H2-Ab. Comparison to DeepAb, IgFold, AlphaFold-Multimer and EquiFold. All runtime are measured on a single NVIDIA A100 GPU with 21 CPU cores.

In Fig. C3, we report the time consumption of various antibody structure prediction methods on the SAbDab-22H2-Ab benchmark. All run times are measured on a single NVIDIA A100 GPU with 21 CPU cores. For AlphaFold-Multimer, all the sequence and template databases are stored on a distributed file system (Ceph), thus the time consumption may be further reduced if local SSD disks are used instead. Consequently, we also report its execution time excluding MSA and template search procedures, denoted as ‘AF-Multimer (NN-only)’.

tFold-Ab is able to predict full-atom antibody structures in 1.15 seconds, only slower than ‘IgFold (w/o PyRosetta)’ which only produces backbone structure predictions and ‘EquiFold’ which replaces pre-trained language models with geometrical structur representation. It is noteworthy that the pre-trained language models do not constitute a significant computational bottleneck within our framework. Although marginally slower than EquiFold, tFold-Ab demonstrates better prediction accuracy.

### C.5 tFold-Ag

#### C.5.1 tFold-Ag inference

Algorithm 8 outlines the inference pipeline of tFold-Ag. The tFold-Ag receives input features derived from the antibody sequence, antigen MSA and outputs including atom coordinates and confidence scores.

The entire network is composed of three components. The first part of the network generates antibody features from the sequence using the pre-trained tFold-Ab model (see Appendix section C.4 for details). The second part generates antigen features from the MSA using the pre-trained AlphaFold2 model [17]. It’s important to note that both of these models are used with their parameters frozen. This approach allows us to leverage the knowledge these models have already learned without overfitting to the new data. Following these initial steps, the AI-driven flexible docking module comes into play as the third component. This module employs a FeatureFusion module (Appendix section C.5.2) to integrate the features of the antibody and antigen. This integration generates an embedding of the antibody-antigen complex, which is then updated iteratively through the EvoformerSingle Stack. Finally, the structure module uses this embedding to predict the final structure of the antibody-antigen complex. This comprehensive approach allows us to accurately model the complex interactions between antibodies and antigens, leading to more precise predictions.

##### Algorithm 8

Model inference for tFold-Ag

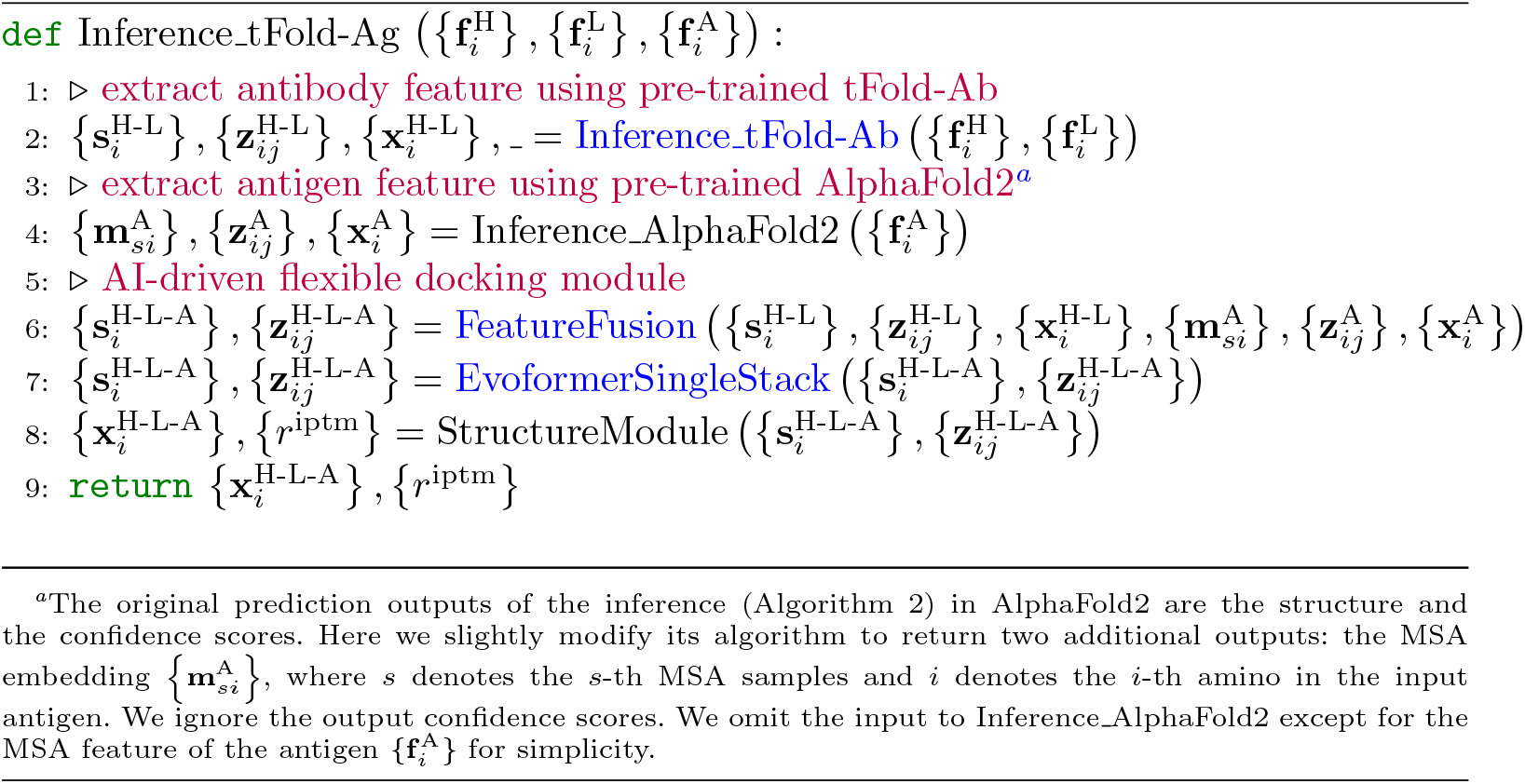

Once additional inter-chain features are available, tFold-Ag can incorporate these features to improve prediction accuracy using the Inter-chain feature embedding module (Appendix section C.5.3). This module is designed to capture the interactions between different chains in the antibody-antigen complex, providing a more accurate representation of the antibody-antigen pair.

In addition to this, if the antibody sequence contains amino acids that need to be designed, tFold-Ag employs a sequence prediction head to determine the amino acid types for the targeted design region. This means that tFold-Ag is capable of predicting the structure and simultaneously recovering the sequence. (Appendix section C.5.4). This dual functionality makes tFold-Ag a versatile tool that can be used for antibody design tasks without any fine-tuning. This is a significant advantage, as it allows researchers to not only use the model directly for their design tasks, but also to infer the function of the designed antibodies based on their predicted structures. This capability can greatly expedite the process of antibody design and validation, saving both time and computational resources.

#### C.5.2 Feature fusion module

As mentioned earlier, we transform the docking task into a complex prediction task based on the given monomer feature. In order to enable the model to predict structures using a paradigm similar to the ‘Evoformer-Single stack+structure module’ employed by tFold-Ab, we have designed a Feature Fusion Module to generate the initial features for antibody-antigen pairs.

**Fig. C4:**
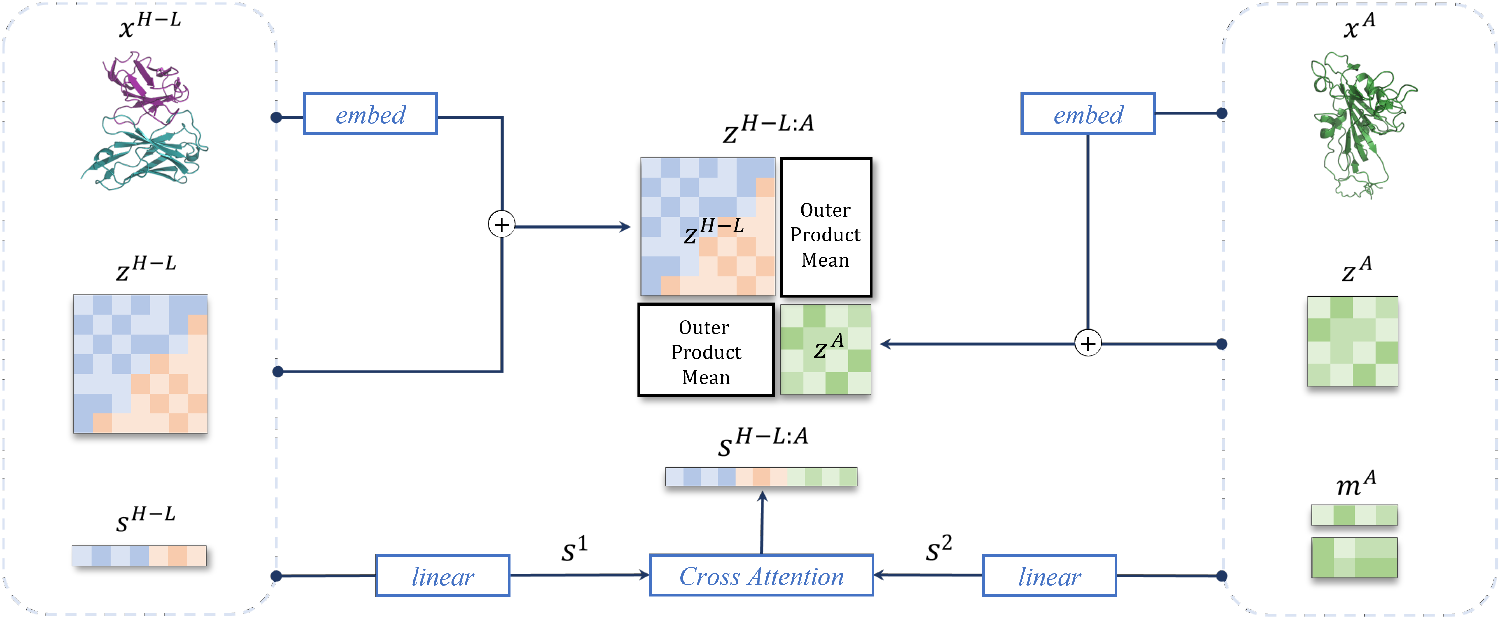
Feature fusion module for tFold-Ag.

As shown in Fig. C4 and Algorithm 9, the diagonal blocks (*z*^ab^ and *z*^ag^) of pair features are initialized by the pair features of the antibody and antigen chains and the initial coordinates. The predicted coordinates of the beta carbon atoms (or alpha carbon for glycine) are used to compute pairwise distances. These distances are then discretized into 20 bins of equal width 1.25Å, spanning a range up to approximately 20Å. The resulting one-hot distogram is linearly projected and added to the pair representation update, similar to the recycle embedding algorithm described in 7. For off-diagonal ones, we use ‘OuterProductMean’ in Algorithm 10 to transform the antigen MSA representation and antibody sequence representation into inter chain pair feature.

##### Algorithm 9

Feature Fusion Module for tFold-Ag

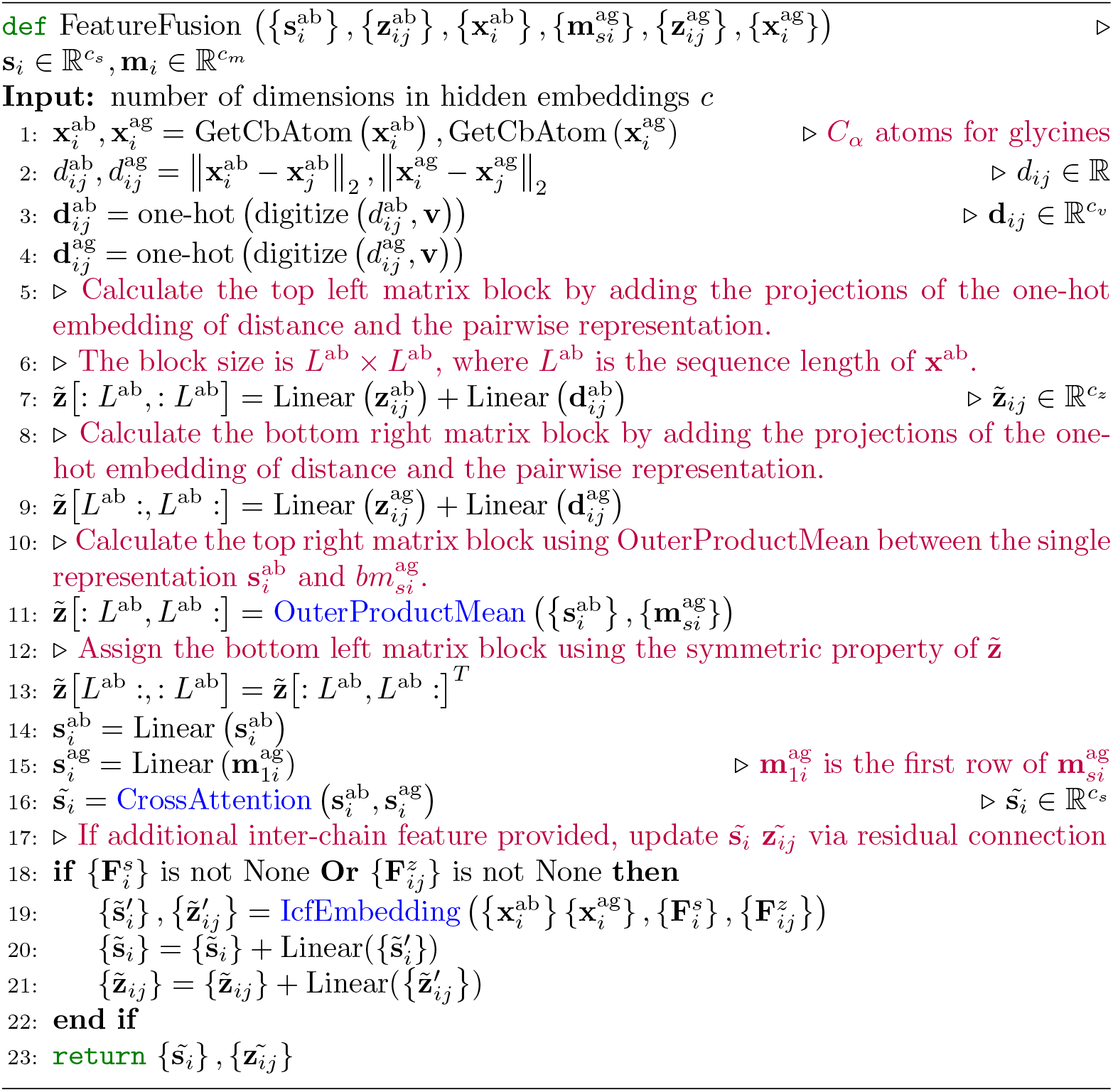

Our ‘OuterProductMean’ algorithm is adapted from the AlphaFold ‘Outer product mean’ algorithm (Algorithm 10 in [17]). While the algorithm in AlphaFold is an MSA-based prediction method with paired antibody-antigen MSA input, our algorithm is specifically designed for the antibody’s sequence feature and antigen MSA feature. For each position *i* in the antibody sequence and each column *j* in the antigen MSA, we linearly project their sequence features into a lower-dimensional space using two separate linear transformations. We then calculate the outer products of these lower-dimensional vectors from the two columns, average them across the sequences, and subsequently project them into a ℝ*c* dimensional space. This process yields an updated value for the (*i, j*) entry in the pair representation’s off-diagonal block. Given the symmetric nature of the interactions and distance map, the values in the two off-diagonal regions, (*i, j*) and (*j, i*), are equal. This algorithm simulates the ‘pairing’ between antibody and antigen sequences. By doing so, it allows us to discern the nuanced interactions between them, enhancing the accuracy of our predictions.

**Fig. C5:**
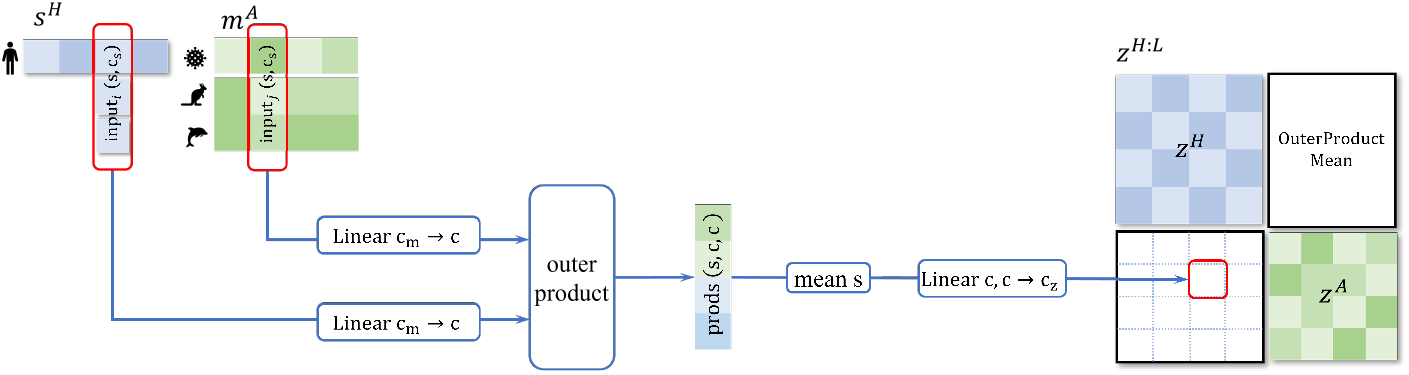
OuterProductMean Module

##### Algorithm 10

Outer product Mean for tFold-Ag

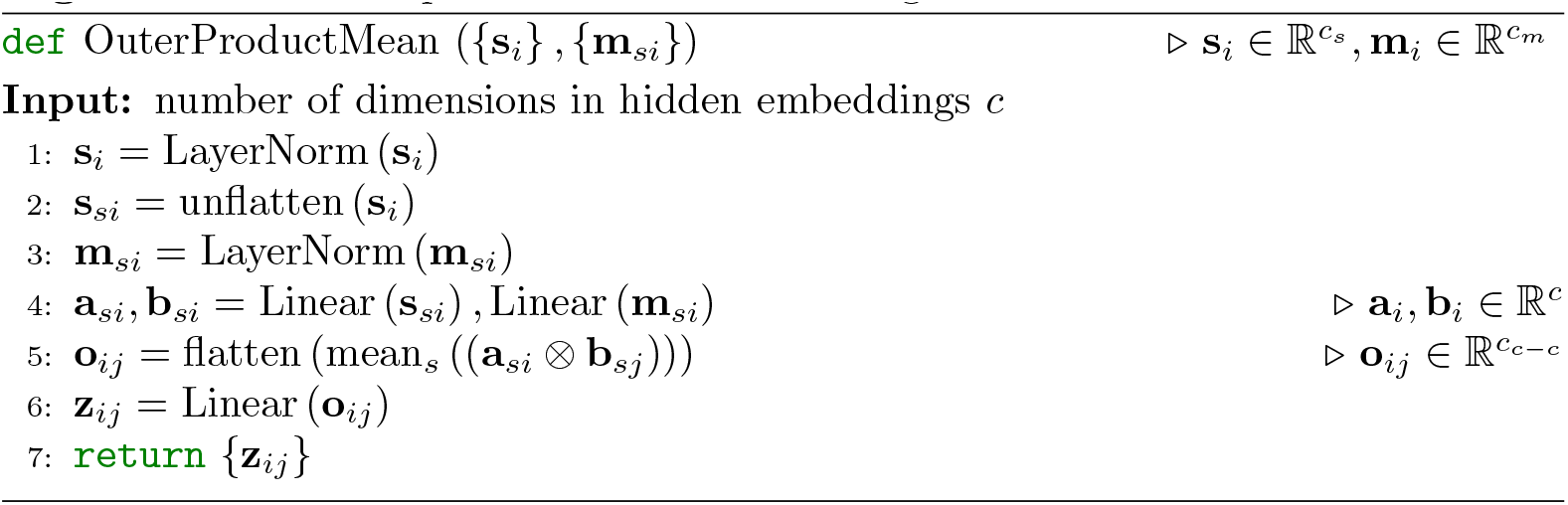

Regarding the initial sequence feature, we employ cross attention [59] to generate sequence embedding for the antibody-antigen complex, with the specifics outlined in Algorithm 11. Cross-attention permits each residue within one sequence to focus on every other residue within the counterpart sequence. By using cross-attention, the output embeds the sequence features of both the antibody and antigen into a unified distribution, facilitating faster convergence of the model.

##### Algorithm 11

Cross attention for tFold-Ag

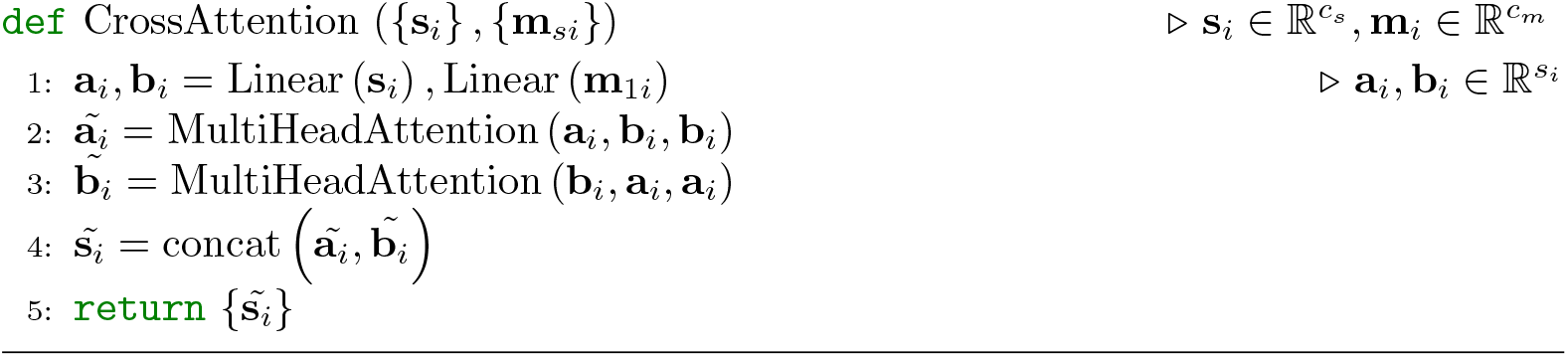

#### C.5.3. Inter-chain feature embedding module

Once additional inter-chain features are available, an Inter-chain feature embedding module embeds these features and adds them via a residual connection to the output of the feature fusion module.

PPI and inter-chain contact are used to encode the additional inter-chain feature. **PPI** feature 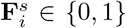 is an optional binary flag indicating whether the **C**_*α*_ atom of residue i is within 10 Å of a **C**_*α*_ atom belonging to a residue in a different chain. This flag is set to 0 if this criterion is not met. **Inter-Chain Contact** 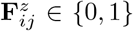 indicates whether two residues in separate chains are in contact. This is determined when the distance between 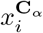 and 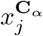 is less than 10 Å. This flag is set to 0 if this criterion is not met.

**Fig. C6:**
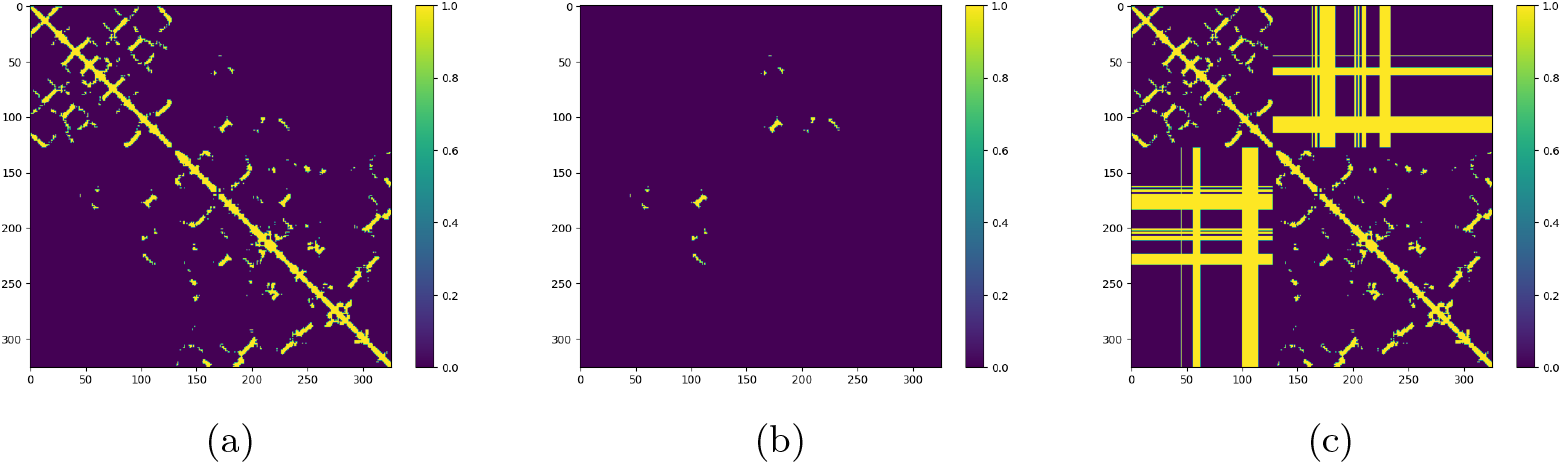
(a) Contact map of chain F and chain A in 7q3r; (b) Inter-chain contact map of chain F and chain A in 7q3r; (c) Pesudo contact map of chain F and chain A in 7q3r. All inter-chain positions are generated based on PPI while intra-chain positions are initialized by structures of monomer.

In real-world applications, PPI and Inter-Chain Contact are typically calculated through different experimental methods and cannot be obtained simultaneously. However, they can be converted into each other. Simply put, Inter-Chain Contact can be used to compute PPI, while PPI can be used to generate a ‘pseudo contact’. Fig. C6 presents the maps of three features: contact, inter-chain contact, and pseudo contact. When inter-chain contact is provided, the intra-chain contact is extracted from the antibody structure coordinates predicted by tFold-Ab and the antigen structure coordinates predicted by AlphaFold to complete the entire contact map. Given the high prediction accuracy of tFold-Ab for antibodies and AlphaFold for antigens, the generated contact map is accurate. We denote the process to generate the entire contact map as an algorithm **GenContact**, which takes the structure coordinates of antibody and antigen and the inter-chain contact as input.

If inter-chain contact is not provided, we generate the pseudo inter-chain contact using the PPI features as input by the algorithm **PPI2Contact**. Based on the PPI features, PPI2Contact returns an inter-chain contact where all rows and columns on the contact map with any positive PPI are set to 1. This, integrated with intra-chain contact, results in a complete contact map, which we refer to as pseudo-contact. The actual inter-chain contact is a subset of the inter-chain contact of the pseudo contact. This feature construction approach allows both types of features to have the same data format, enabling a single model to be trained to accommodate multiple input types. If the PPI features are not available, we generate the PPI features using inter-chain contact by the algorithm **Contact2PPI**. Concretely, Contact2PPI returns a PPI feature whose *i*-th entry is set to 1 if there exists any positive entry in the *i*-th column of the inter-chain contact.

At the beginning of training, we introduce no additional features, PPI, or contact features in a probabilistic manner for each input. Specifically, we allocate a probability of 40% for including no additional features, 30% for incorporating PPI features, and 30% for integrating contact features. This approach ensures a balance to various types of input data, enhancing the model’s ability to generalize across different scenarios. Furthermore, we fine-tune three models, each tailored to one of the three distinct input types, to address specific application contexts and optimize performance in each respective input data.

##### Algorithm 12

Inter-chain feature embedding for tFold-Ag

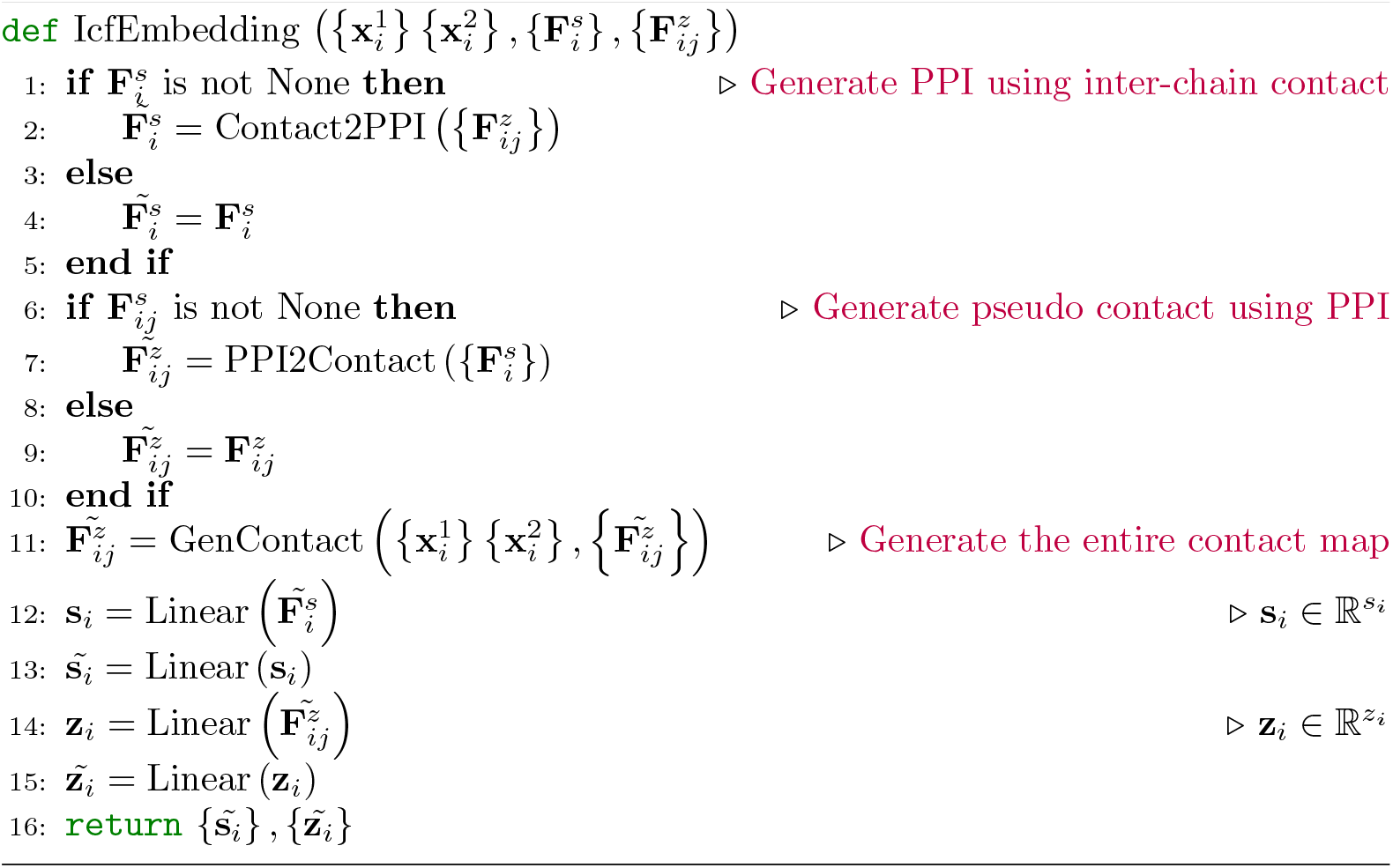

Algorithm 12 shows the inter-chain feature embedding module for tFold-Ag. Note that exactly one of the inputs from 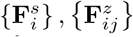 will be provided to the algorithm. To accommodate various inter-chain features, we transform the inter-chain feature into uniformly formatted sequential and pairwise features. These are subsequently processed through two distinct linear layers, generating inter-chain sequence and pair representations respectively. The outputs of the Inter-chain embedding module are integrated with the output of the Feature Fusion module via a residual connection.

This approach ensures that the model’s training process remains undisturbed, making it a robust method for handling diverse input types.

In real-world applications, information regarding the epitope, paratope, or contact map may be limited or even unknown. Acknowledging this, our approach involves providing only a select number of contacts or binding residues, sampled randomly for each training example. Specifically, when an antibody-antigen complex requires the incorporation of PPI features as input, we adopt a random subsampling strategy. A designated number of interface residues are selected with a probability of 50%. This approach ensures that only half of the interface residues are chosen, thereby providing a diverse and representative sample for model training. Similarly, only half of interchain contacts are selected when this feature is used, further bolstering the robustness of the methodology, and making it suitable for real-world application.

#### C.5.4 Sequence recovery for antibody design

Due to the complexity of antigen-antibody structure prediction, the reverse process, antibody design based on an antigen, has not been well resolved. Existing antibody design methodologies predominantly depend on the availability of known antibody-antigen complex structures, predefined epitopes, and antigen conformations, or they fail to consider the intrinsic interactions within the antigen. We have integrated the tasks of structure prediction and sequence design into a single challenge, structure and sequence co-design. For areas requiring design (*e*.*g*.,CDR-H3 loop), amino acids are substituted with a *<*Mask*>* placeholder, which is then processed by ESM-PPI to extract antibody characteristics. Given that tFold-Ab solely utilizes features generated by ESM-PPI as input, and the Feature Fusion Module only requires the sequence embedding, pairwise representation and predicted coordinates of the beta carbon atoms, there is no need for additional fine-tuning for tFold-Ab. After the Evoformer-Stack module within the Docking module, we append a sequence prediction head to determine the amino acid types for the targeted design region. These predicted amino acid types are then fed into the Structure Module, thereby facilitating the prediction of the antibody-antigen complex structure, inclusive of the side chains. In our sequence recovery model, we consider not only sequence information but also structural information. As a result, the model is capable of generating more rational sequences based on the complex structures of antigens and antibodies. Furthermore, as ESM-PPI utilizes antibody sequences for pre-training, our model has not only learned from the antibody sequences with known structures in the SAbDab database, but also to some extent from the antibody sequences without known structures in the OAS dataset. This knowledge aids in the improved design of the CDRs of antibodies.

#### C.5.5 Loss function

Given the architectural similarities between tFold-Ab and tFold-Ag, they share similar loss functions. As tFold-Ab is capable of predicting features of both antibody and nanobody, and the Feature fusion module treats the antibody as a whole component (whether paired or unpaired), the loss function is identical for both antibody-antigen complexes and nanobody-antigen complexes. The loss term for the antigen-antibody complex is similar to the antibody loss term shown in Equation C2, following the same format:

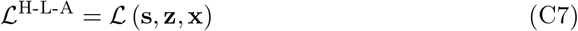

Concretely, this loss function constitutes of the following terms:

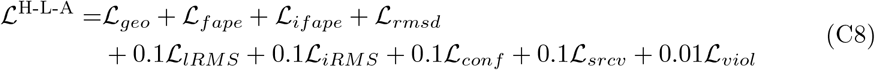

Among these, the Inter-residue geometric loss *ℒ*_*geo*_, Frame aligned point error *ℒ*_*fape*_, Interface frame aligned point error *ℒ*_*ifape*_, Coordinate RMSD loss *ℒ*_*rmsd*_, Confidence loss *ℒ*_*conf*_, and Structure violation loss *ℒ*_*viol*_ are identical as used by tFold-Ab, which are described in Section C.4.5. We will mainly introduce the remaining loss terms. **Ligand RMSD loss** *ℒ*_*LRMS*_ **and interface RMSD loss** *ℒ*_*iRMS*_: DockQ combines three different evaluation criteria: the contact surface (Fnat), the ligand RMSD (lRMSD), and the interface RMSD (iRMS). To better optimize the docking accuracy of the antibody and antigen, in addition to the coordinate RMSD loss, we calculated the lRMS Loss and iRMS Loss by aligning different regions. For lRMS, we use the Kabsch algorithm to find the optimal rotation and translation between the point clouds corresponding to the longest chain in the real and predicted structures of the antigen and antibody. We then calculate the lRMS loss in the other chain with the predicted and real structures. For iRMS, we align two sets of point clouds of the whole complex by Kabsch algorithm. We then calculate the iRMS loss at the interface of with the predicted and real structures. The interface is defined by residues with contact cut-off of 10 Å. The *ℒ*_*LRMS*_ and *ℒ*_*iRMS*_ resulting are penalized with a clamped L1-loss with a length scale *Z* = 30Å to make the loss unitless.

For the *i*-th residue’s *j*-th atom, we denote its 3D coordinate in the ground-truth structure as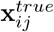, and 3D coordinate in the aligned predicted structures as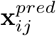 . *C*_1_ refers to the longer chain in the antibody and antigen, while *C*_2_ refers to the shorter one. *T*_*C*_ represents SE(3)-transformation from the prediction frame to the ground-truth reference frame of given chain *C*, if the chain is not specific, the whole complex will be employed. The lRMS loss is defined as:

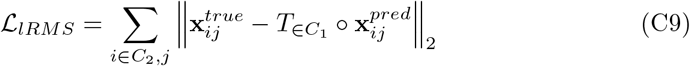

If *R*_*i*_ refers to the region of the interface, the iRMS loss is defined as:

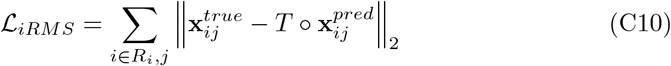

##### Sequence recovery loss *L*_*srcv*_

As illustrated in Section C.5.4, we use the sequence representation to predict amino acid types that have been previously masked out. A total 20 classes for common amino acid types are considered. sequence embedding *{***s**_*i*_*}* are linearly projected into the output classes and scored with the cross-entropy loss:

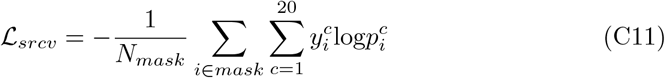

where 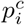 are predicted class probabilities, 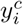 are one-hot encoded ground-truth values, and averaging across the masked positions.

#### C.5.6 Training detail of tFold-Ag

**Table C17:**
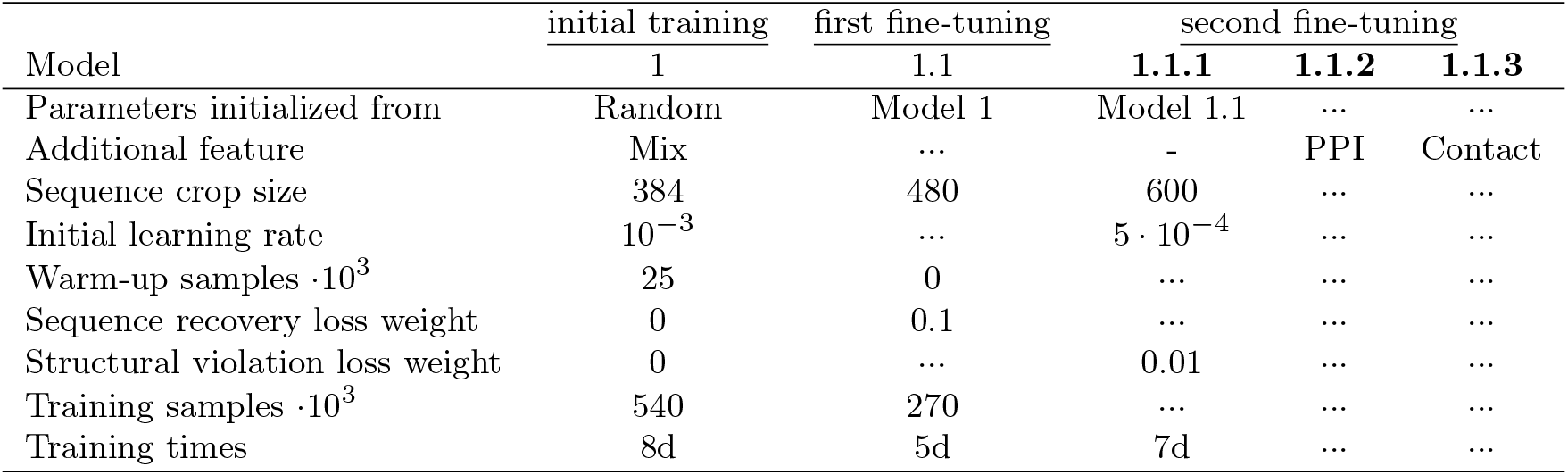
Training protocol for tFold-Ag models. The model in **bold** (i.e **1.1.1** - **1.1.3** were be released. We report the number of training samples and the training time of whole training stages. Three dots (…) indicate the same value as in the former column. Hyphen (-) means that do not input any additional feature.

In order to enable tFold-Ag for the co-design of antibody structure and sequence, we have integrated a masked language model (MLM) training strategy. During the training process, all amino acid sequences in the antibody’s CDRs loop are selected with 30% probability, aligning to the strategy used to train ESM-PPI. Unlike traditional Mask language models [15, 54, 55, 90] (mask all residues with 15% probability), our focus is primarily on the CDRs of the antibody, making this task more challenging. Beyond just sequence information, we aim to improve antibody design performance by incorporating structural information from both the antigen and the antibody.

For the training regimen, we utilize the AdamW [82] optimizer with a batch size of 32. We also maintain an EMA of the model parameters, applying a decay factor of 0.999, and employ this EMA model for evaluation purposes. The model iteration that achieves the highest DockQ scores on the validation subset is selected as the optimal model.

We adopt the training procedure of AlphaFold2, with the specific details of the three-stage training process outlined in Table. C17. Initially, Model 1 is trained with a smaller crop size and a mixture of inter-chain features for approximately one week. Subsequently, this model is then fine-tuned by enlarging the crop size and integrating MLM objectives alongside sequence recovery losses, the latter of which enhances training stability. Subsequent fine-tuning involves an expanded sequence crop size, the incorporation of violation losses, and a diminished learning rate. During this phase, we refine three distinct models: Model 1.1.1, which forgoes inter-chain features (excluding the corresponding parameters from the inter-chain feature embedding module); Model 1.1.2, which incorporates the Protein-Protein Interaction (PPI) feature; and Model 1.1.3, which utilizes the contact feature.

All tFold-Ag models employ a pre-trained tFold-Ab model for antibody feature extraction and a pre-trained AlphaFold2 model (specifically, we use params model 4 ptm as it demonstrates the best performance in antigen structure prediction when template features are not used). Attempts were made to update the tFold-Ab parameters during training and to apply a mixed loss function akin to that described in C.4.5 for tFold-Ag training. However, this method proved to be excessively time-consuming and did not significantly enhance performance.

#### C.5.7 Ablation study of tFold-Ag

We estimate the relative importance of key components of the tFold-Ag architecture by training and evaluating a number of ablation models:

##### Baseline

The baseline model, as described in the paper, does not include the Interchain feature embedding module. In other words, it corresponds to the version of the tFold-Ag model.

##### Simplified feature fusion module

We simplified the Feature fusion module in tFold-Ag in the following ways: For sequence embedding, we directly concatenated the antibody’s sequence embedding and the antigen’s sequence embedding, without using cross attention as described in Algorithm 11. For pairwise representation, instead of using OuterProductMean as described in Algorithm 10 to initialize the offdiagonal area of the antibody-antigen complex, we directly used zero initialization. This approach does not make full use of the antigen’s MSA features and relies solely on the subsequent Evoformer-Single Stack to optimize the inter-chain information of the antibody-antigen pairs.

##### No iRMS & LRMS Loss

We did not use iRMS Loss and LRMS Loss to guide the training of tFold-Ag. During the training phase, the weights of these two losses were set to zero.

##### With recycling for the pre-training structure prediction model

We set the recycling iteration of AlphaFold to 3, which matches the design of AlphaFold. As shown in Table B10, this can make the initial structure of the antigen more accurate and the features better. Consistent with training strategy for AlphaFold, we uniformly sample the number of recycling iterations between 1 and 3 during training. Every batch element trains the same iteration on each step. We take the feature input from the last recycling iteration as the input for the Feature Fusion module. During the evaluation, we set the recycling iteration of AlphaFold to 3.

##### With recycling in AI-driven flexible docking module

We employ a recycling strategy similar to tFold-Ab during both the training and inference phases (refer to Algorithm 7 for details). The difference is that the recycling strategy of tFold-Ag integrates the single and pair features from the previous iteration, as well as the embedding of coordinates of Cb, into the initial antibody-antigen complex features produced by the Feature Fusion. Specifically, the recycling iterations are set to 3 for further refinement. The recycling models are fine-tuned based on model 1 described in Table C17.

For all ablations, we kept the hyperparameters from the main model configuration, which we have not re-tuned. All models did not use any extra structure restraints, in other words, none of the models had an Inter-chain feature embedding module. In addition, except for the baseline model, no other models used the Mask language model for sequence recovery. For each ablation, we trained the model with the same random seeds, all proteins used the same validation set, and we selected the model using the same metrics. Except for the models specifically mentioned as fine-tuned, all other models were trained from scratch. Due to the high cost of training, we did not conduct many ablation experiments. Because the number of Nanobody-Antigen complexes is too small, we only used the test sets described in Section 4.1.2, SAbDab-22H2-AbAg, for ablation analysis.

**Table C18:**
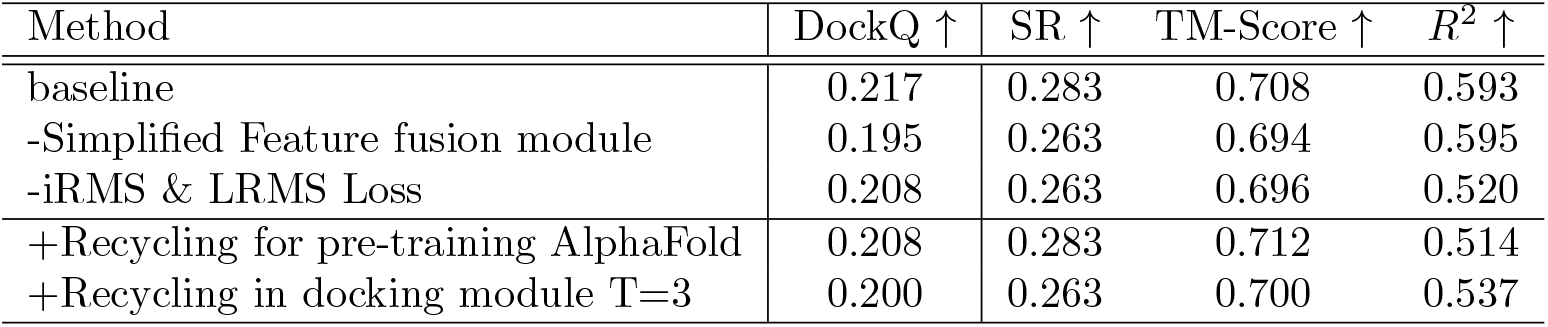
Ablations performance for tFold-Ag on the SAbDab-22H2-AbAg benchmark.

Ablation results are presented in Table C18. The Feature Fusion module we designed has improved the prediction accuracy of the antibody-antigen complex, with the average DockQ on the SAbDab-22H2-AbAg test set increasing from 0.195 to 0.217. This module simulates the pairing between antibody and antigen sequences and allows the model to find the rare co-evolution information from the antigen MSA and antibody sequence features, enhancing the accuracy of our predictions. The use of iRMS and LRMS Loss slightly improved the DockQ by 0.01, which makes the model pay more attention to the binding area of the antigen and antibody.

Using the AlphaFold Recycling strategy to generate superior antigen feature inputs for the docking module did not enhance the performance of the antibody-antigen complex prediction. Despite the fact that the features input to the Docking module would be improved when there are already sufficiently high-quality monomer antigen features, an increase in the accuracy of monomer antigen structure prediction does not impact the performance of antibody-antigen docking. Therefore, in this scenario, we disabled AlphaFold Recycling to boost computational efficiency.

Using Recycling in the Docking module also did not enhance the model’s performance. This could be because the initial sequence embedding and pairwise representation obtained from the Feature Fusion module are already good enough. Alternatively, it could be because the volume of antibody structure data is too limited, meaning that increasing the network depth does not yield any additional information.

#### C.5.8 Confidence score for complex prediction

AlphaFold-Multimer [5] has designed ipTM (Interface pTM) and employs a weighted combination of pTM and ipTM as model confidence metric. For tFold-Ag, we utilize the antibody features extracted by tFold-Ab along with the antigen features extracted by AlphaFold. In this scenario, both the initial structure of the antibody and the antigen are relatively accurate. Since TM-Score always aligns the longer chain in the antibody and antigen, the TM-Score will be greater than 0.5 even if the predicted antigen and antibody are arbitrarily arranged. Therefore, pTM is not a suitable metric for tFold-Ag, and we solely use ipTM as the confidence score. We treat the two chains of the antibody as a single entity, thereby enabling a unified approach for calculating the ipTM of both antibodies and nanobodies.

Fig. C7 illustrates the correlation between ipTM and DockQ for tFold-Ag. We evaluated three types of inputs: without using any Inter-Chain feature, antigen epitope, and PPI features. Regardless of the scenario, the confidence score of tFold-Ag has a strong correlation with DockQ, especially for the Antibody-Antigen data. The confidence scores for the predictions of the three models, which include those without any Inter-Chain feature, with antigen epitope, and with antigen epitope & antibody paratope features, have Pearson correlations of 0.77, 0.59, and 0.64, respectively. However, for the Nanobody-Antigen data, the correlation of the confidence score is less robust than that of the Antibody-Antigen data, with values of 0.49, 0.43, and 0.48 respectively. The primary reason for this is that in SAbDab, there is less data available for nanobodies compared to antibodies.

**Fig. C7:**
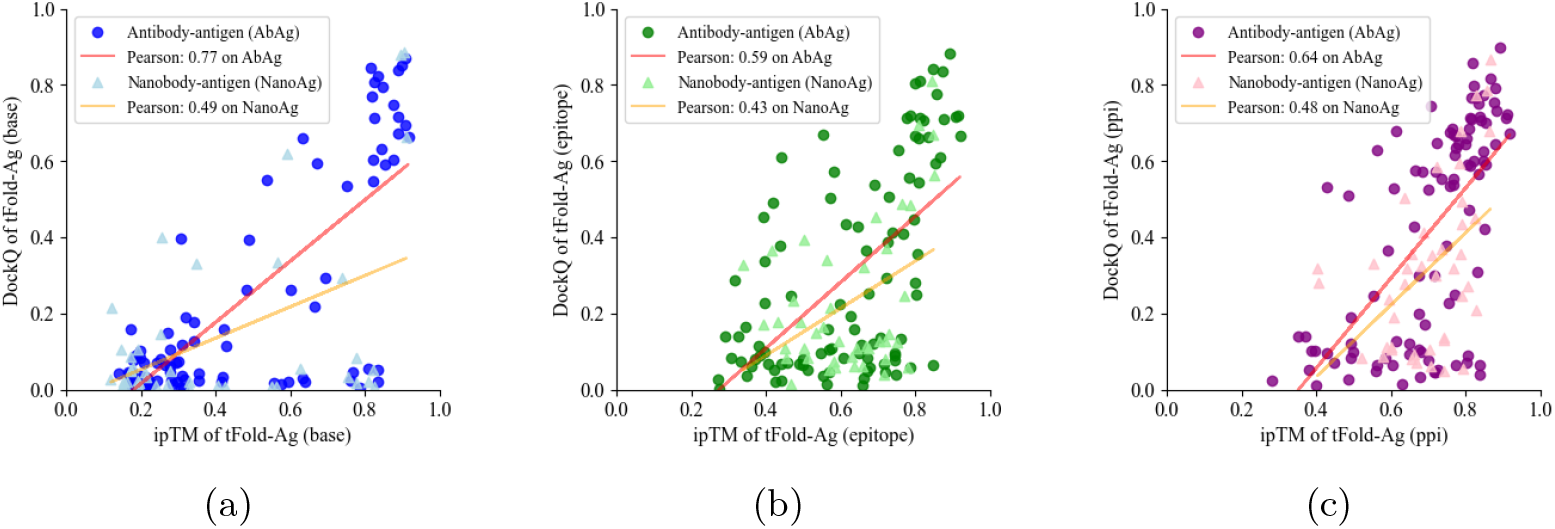
Correlation between ipTM and DockQ scores for various tFold-Ag predictions (base, epitope and ppi) on the SAbDab-22H2-AbAg and SAbDab-22H2-NanoAg datasets, with a least-squares linear fit applied to the data. (a) tFold-Ag. (b) tFold-Ag-ppi (epitope). (c) tFold-Ag-ppi (epitope and paratope).

Given the strong correlation between ipTM scores and prediction accuracy, we consider using ipTM for model ensembling, which will be discussed in section C.5.9.

#### C.5.9 Ensemble analysis

Fig. C8 presents a point-to-point comparison between tFold-Ag and AlphaFold-Multimer, The boundary for determining successful predictions (DockQ *≥* 0.23) is indicated in the figure. Although tFold-Ag exhibits a higher success rate, its advantage in structural prediction performance is relatively minor when compared to AlphaFold-Multimer on average. In a test set comprising 99 Antibody-Antigen complexes, tFold-Ag demonstrated superior performance in 49 cases. Similarly, within a test set of 43 Nanobody-Antigen complexes, tFold-Ag outperformed in 21 cases. tFold-Ag does not use MSAs for antibodies, whereas AlphaFold-Multimer employs paired MSAs to extract co-evolutionary information between antigens and antibodies. Even for antigen-antibody pairs, this co-evolutionary information provides limited assistance. The distinct inputs used by tFold-Ag and AlphaFold-Multimer result in highly complementary outcomes, which are shown in Fig. C8a and Fig. C8b.

**Table C19:** Ensemble analysis on the SAbDab-22H2-AbAg (99 complexs) and SAbDab-22H2-NanoAg benchmark (43 complexs). tF refers to tFold-Ag, and AF refers to AlphaFold-Multimer.

| Method | Antibody-Antigen |  |  | Nanobody-Antigen |  |  |
| --- | --- | --- | --- | --- | --- | --- |
| | DockQ | SR | $R^2$ | DockQ | SR | $R^2$ |
| AF | 0.158 | 0.182 | 0.600 | 0.149 | 0.140 | 0.545 |
| tF | 0.217 | 0.283 | 0.593 | 0.136 | 0.186 | 0.237 |
| tF & AF (ensemble) | 0.288 | 0.404 | 0.548 | 0.218 | 0.279 | 0.407 |
| tF & AF (best) | 0.307 | 0.414 | - | 0.246 | 0.279 | - |

Both tFold-Ag and AlphaFold-Multimer can output a confidence score while predicting structures, and this score is highly correlated with prediction accuracy. Therefore, we applied a common ensemble strategy in machine learning, using the confidence score to select the higher-scoring prediction from AlphaFold-Multimer and tFold-Ag as the final result. Table C19 demonstrates the performance of this ensemble approach. Although AlphaFold-Multimer and tFold-Ag have different definitions for confidence scores, using the higher-scoring model from the two methods as the prediction result significantly improves the overall prediction accuracy.

Fig. C8c shows that for most of the test proteins, the better model can always be selected by comparing confidence scores. Therefore, AlphaFold-Multimer and tFold-Ag can be used together to further improve the prediction accuracy of antigen-antibody complexes.

#### C.5.10 Inference speed

We conducted a comparative analysis of the runtime between tFold-Ag and AlphaFold-Multimer in two distinct scenarios. Firstly, we evaluated the time efficiency of both methods in predicting the complete structure of antigen-antibody pairs of varying lengths. This scenario corresponds to the conventional task of antigen-antibody complex structure prediction. Secondly, we assessed the time required by both tFold-Ag and AlphaFold-Multimer to predict a varying number of antibodies for a given antigen. This scenario is representative of the structure-based virtual screening of antibodies.

**Fig. C8:**
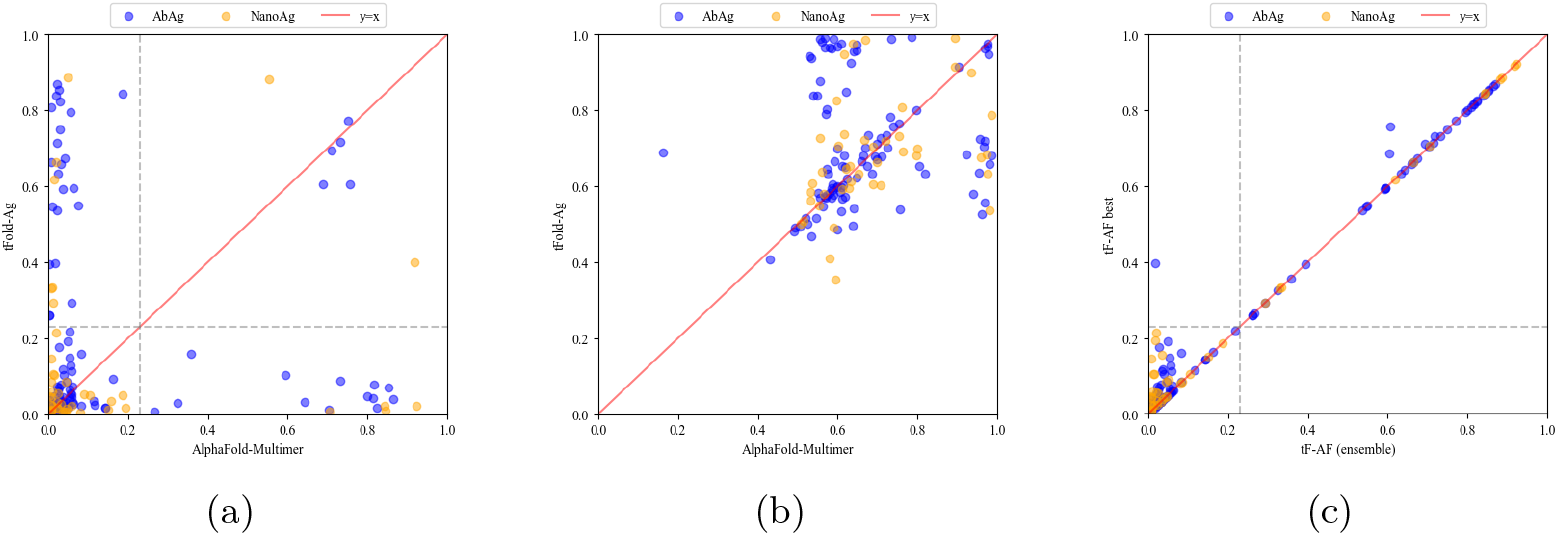
Ensemble analysis of tFold-Ag and AlphaFold-Multimer. (a) The head-to-head comparison between tFold-Ag and AlphaFold-Multimer on the SAbDab-22H2-AbAg and SAbDab-22H2-NanoAg. The model is evaluated by DockQ. (b) The head-to-head comparison between tFold-Ag and AlphaFold-Multimer on the SAbDab-22H2-AbAg and SAbDab-22H2-NanoAg. The model is evaluated by TM-Score. (c) The head-to-head comparison between the best model generated by tFold-Ag and AlphaFold-Multimer and the ensemble model selected by confidence score.

**Fig. C9:**
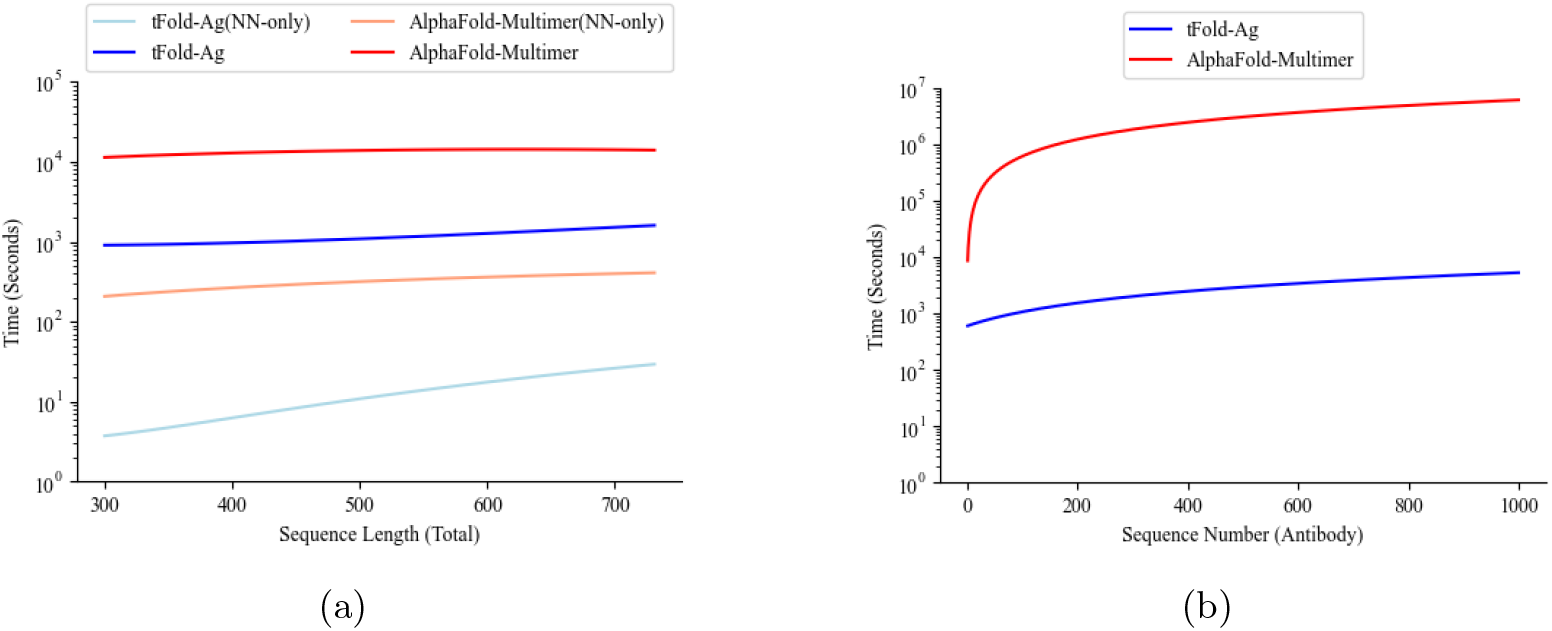
Runtime for tFold-Ag. (a) The runtime of tFold-Ag compared to AlphaFold-Multimer with different sequence length. (b) The runtime of tFold-Ag compared to AlphaFold-Multimer with different sequence number.

In Fig. C9, we report the time consumption of tFold-Ag and AlphaFold-Multimer on two scenarios. All run times are measured on a single NVIDIA A100 GPU with 21 CPU cores. For both AlphaFold-Multimer and tFold-Ag, sequence database searches are required to construct the MSA. Therefore, in addition to the total runtime, we also report their execution times excluding the MSA and template search processes. These are denoted as AlphaFold-Multimer(NN-only) and tFold-Ag(NN-only).

tFold-Ag employs a pre-trained protein language model to extract antibody features. For the antibody component, it eliminates the time-consuming process of MSA (Multiple Sequence Alignment) search.

In the task of predicting antigen-antibody complex structures shown in Fig. C9a, tFold-Ag, despite requiring approximately 1000 seconds per antigen to search for homologous sequences and construct an MSA, still achieves a time-saving of nearly two-thirds compared to AlphaFold-Multimer. The latter necessitates the search and construction of MSAs for three distinct sequences: the heavy chain, light chain, and antigen. This time discrepancy is further amplified considering that AlphaFold-Multimer conducts searches across multiple sequence databases, including UniRef90 [16], Uniprot [91], MGnify [92], and BFD [93], and requires the extraction of template features. Conversely, tFold-Ag’s MSA construction process follows the procedure utilized by ColabFold [43], which is more time-efficient. On average, tFold-Ag is 10 times faster than AlphaFold-Multimer in predicting antigen-antibody complex structures.

For structure-based antibody virtual screening tasks, the antigen sequence is fixed, so the sequence database only needs to be searched once for the antigen. In this case, the advantage of tFold-Ag becomes even more pronounced. Taking the receptor binding domain (RBD) of the spike protein as an example (with a length of 223), we evaluated the time required to predict different numbers of antigen-antibody complex structures, as shown in Fig. C9b. Given the comparable lengths of antibody sequences, the time required for tFold-Ag to predict the structures of antigen-antibody complexes exhibits a linear increase when the antigen is held constant, averaging approximately 4.7 seconds per antibody prediction. Conversely, AlphaFold-Multimer necessitates additional time for sequence database searches and constructing MSAs for antibody sequences, which can be considerably time-consuming. The disparity in time requirements between tFold-Ag and AlphaFold-Multimer becomes more pronounced with an increasing number of antibodies to be screened. For instance, when predicting 1000 antibodies, tFold-Ag outperforms AlphaFold-Multimer by a factor of 1000. This substantial time efficiency of tFold-Ag over AlphaFold-Multimer offers significant time savings and computational cost reductions for virtual screening tasks

### C.6 Structure-based virtual screening

#### C.6.1 Confidence analysis of binding antibodies

Fig. C10 illustrates the distribution of confidence scores for both binding and non-binding antibodies across different test sets as predicted by tFold-Ag. For PD-1 shown in Fig. C10a, the overall confidence scores for the predicted structures are high. Specifically, the median confidence score for the structures of binding antibodies is 0.78, indicating that tFold-Ag is capable of assigning high confidence ratings to the majority of binding structures. This is particularly noteworthy given that none of the antibodies in the test set were included in the training set, yet tFold-Ag demonstrates high prediction performance.

Non-binding antibodies have a median confidence score of 0.48, as predicted by tFold-Ag, suggesting that the confidence scores can serve as a useful metric for distinguishing antibodies that can bind to PD-1 from those that cannot. This distinction in confidence scores is a strong indicator of tFold-Ag’s utility in screening a wide array of candidate antibodies to identify those capable of binding to PD-1.

**Fig. C10:**
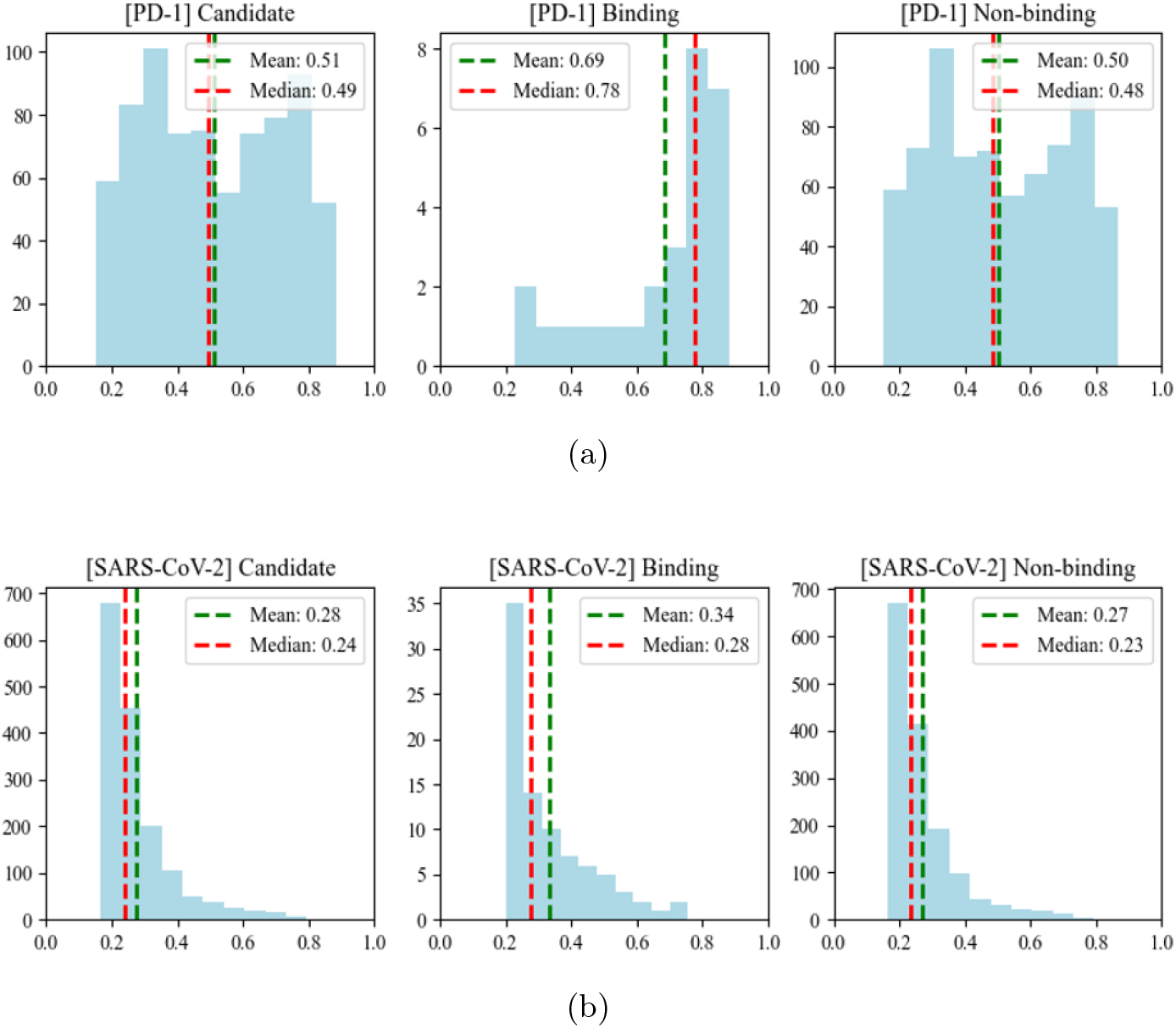
Distribution of confidence scores for binding and non-binding antibodies in different test sets as predicted by tFold-Ag. The red dashed line indicates the mean, while the green dashed line represents the median. (a) PD-1 set. (b) SARS-CoV-2 set.

Regarding the SARS-CoV-2 dataset, Fig. C10b shows that the overall structural prediction confidence score of tFold-Ag is relatively low, which could be due to the following two reasons. Firstly, The RBD (Receptor Binding Domain) is a relatively conserved part of the S protein in the newly discovered antigen SARS-CoV-2 virus, and there is a scarcity of homologous sequence data available in colabfold envdb [43]. This is reflected in the Neff of the RBD’s Multiple Sequence Alignment (MSA), which is 4.9. In contrast, the Neff of the MSA for PD-1, a well-studied protein, is 8.3. This limitation affects the ability of tFold-Ag to extract antigenic features effectively, resulting in the docking module’s inability to fully utilize the antigen’s characteristics for accurate prediction. The diversity and quantity of homologous sequences are crucial for deep learning-based structural prediction models, as these models rely on extensive data to learn how to predict protein structures accurately. Secondly, The RBD is a part of the spike protein, which typically exists as a trimer in biological systems. Modeling the RBD domain in complex with an antibody in isolation may not adequately capture the structural dynamics and epitope integrity when the entire spike protein interacts with an antibody. This means that the model might not accurately predict the binding between an antibody and the full spike protein because it does not consider the spatial constraints and interactions of the trimeric structure.

Despite the lower prediction accuracy of tFold-Ag for the SARS-CoV-2 set, its confidence scores can still somewhat differentiate between binding and non-binding antibodies. For binding antibodies, the average confidence score is 0.34, while for non-binding antibodies, the average confidence score is 0.27.

#### C.6.2 Epitope analysis of binding antibodies

As shown in Fig. C11, for the experimental structures (PDB ID: 4zqk [31]), the binding epitopes for the interaction between PD-1 and PD-L1 are located at the 30th amino acid, between the 40th and 60th amino acids, and between the 90th and 100th amino acids. For the known PD-1 antibodies in the PDB, their antigenic epitopes overlap significantly with PD-L1, indicating that they have the pharmacological function of inhibiting the PD-1/PD-L1 interaction. For the antibodies in our virtual screening task’s PD-1 set, although these antibodies lack corresponding structures, since they are either in clinical stages or approved, we assume they all have the pharmacological function of blocking the interaction between the antigen and its ligand. The structures predicted by tFold-Ag also have similar antigenic epitopes, suggesting that the predicted structures can correctly reflect the function of these PD-1 antibodies.

Figure C12 shows noticeable differences between antibodies with known competitive-or-not labels in the antigen-receptor binding process for known experimental structures in PDB. For the competitive antibody 2G1 (PDB ID: 7X08) [94], the overlap between the antibody-antigen epitope and the antigen-receptor epitope indicates that 2G1 is indeed capable of blocking the antigen-receptor binding process. In contrast, for the non-competitive antibody S309 (PDB ID: 6WPS) [95], we do not observe any epitope overlap between S309 and the ACE2 receptor.

Although the structure prediction confidence of tFold-Ag in the SARS-CoV-2 set is relatively low (with mean 0.34, median 0.28), we hope that the predicted epitopes may still help to determine whether the antibody competes with ACE2. Fig.C12a shows that for competitive antibodies, the antibody epitopes predicted by tFold-Ag have two patterns, one located between residues 125th to 130th and 175th to 185th, and the other located between residues 150th to 185th, both of which overlap with the RBD and ACE2 epitopes. However, we observed similar patterns in non-competitive antibodies as shown in Fig.C12b. This suggests that the low confidence and accuracy of the predicted structures may offer little assistance in determining competitive antibodies. Besides that, another explanation for the above phenomenon is that the non-competitive antibodies may have low RBD binding affinity that fails to compete with ACE2. This is evidenced by non-competitive antibodies, such as 6zdh and 7JW0, which share similar experimental validated binding epitope patterns with competitive antibody 7e5r [96].

**Fig. C11:**
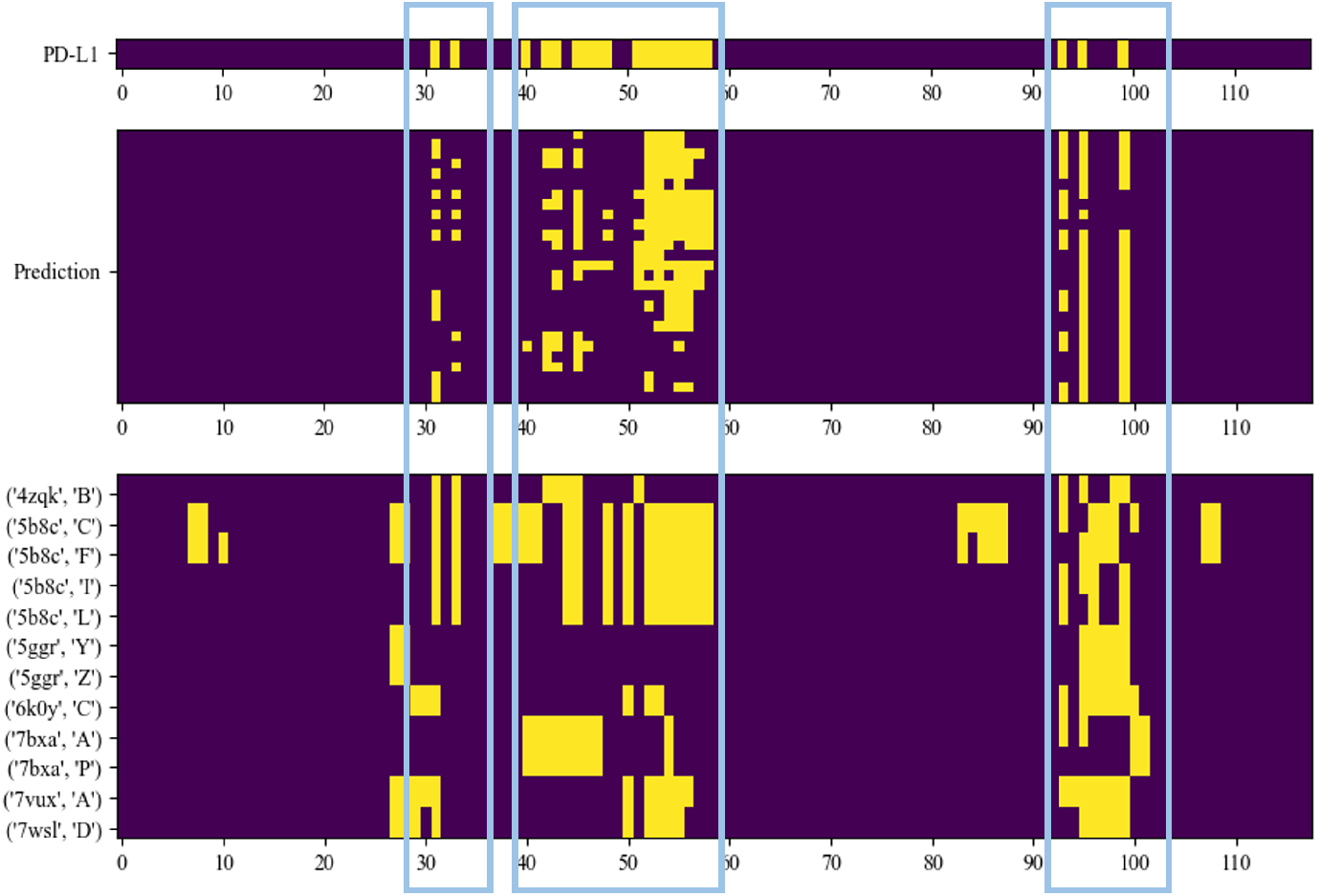
Spatial analysis of binding antibodies for the PD-1 set. A residue is considered part of the binding site if there are *C*_*α*_ atoms in another chain within a distance of less than 10 Å from the residue on the antigen. Top: Binding epitope of PD-1 in complex with PD-L1. Middle: Epitope of PD-1 in the antibody-antigen complexes capable of binding as predicted by tFold-Ag within the PD-1 set. Bottom: Epitope of PD-1 in the experimental structure of the antibody-antigen complex with PD-1 from the PDB. Areas of overlap are highlighted with a blue box.

### C.7 Antibody Design

#### C.7.1 Apply our method to nanobody design

tFold-Ag is capable of predicting both antibody-antigen and nanobody-antigen complex structures. However, due to the limited presence of nanobody-antigen complexes in the training data, the prediction accuracy for these complexes is somewhat lower compared to that for antibody-antigen complexes. The design of nanobodies has received less attention in the field due to this data scarcity. Nevertheless, tFold-Ag can be effectively utilized for nanobody design. To demonstrate this, we have constructed a test set and employed metrics such as the Amino Acid Recovery (AAR) and Contact Amino Acid Recovery (CAAR) to evaluate the nanobody design performance of tFold-Ag.

##### SAbDab-22-DesignNano

Following a similar process to the construction of SAbDab-22-DesignAb, SAbDab-22-DesignNano contains nanobody-antigen complex structures in SAbDab that were released from January 1, 2022, through December 31, 2022. We excluded any nanobody-antigen pairs in SAbDab-22-DesignNano where the antigen sequences shared more than 70% identity with sequences in our training set to avoid redundancy. Only those CDR-H3 regions that were in contact with the antigen were selected to ensure the design task was relevant. Due to the significantly fewer nanobody data in SAbDab compared to antibody data, this test set only contains 26 nanobody-antigen complexes.

**Fig. C12:**
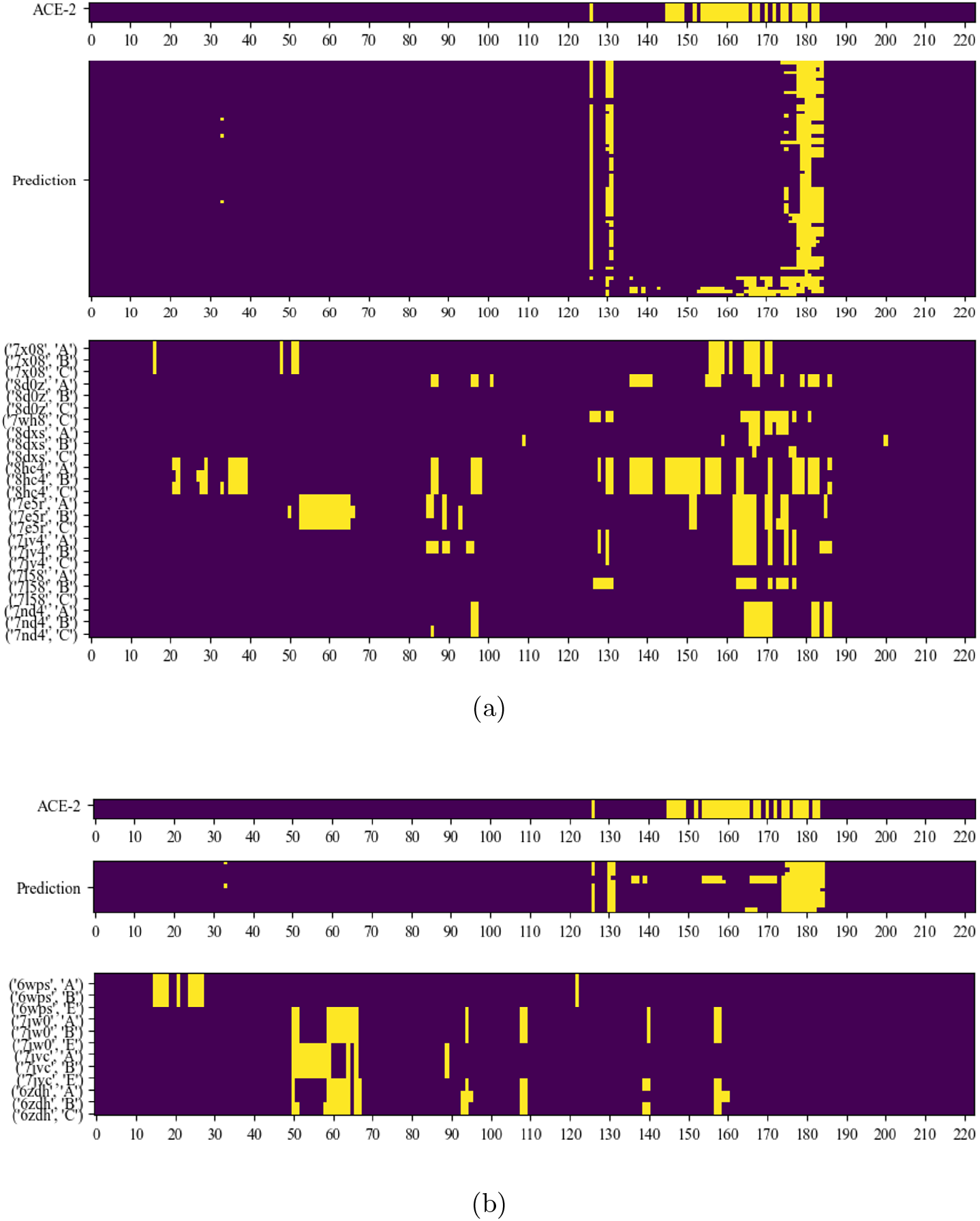
Spatial analysis of binding antibodies for the SARS-CoV-2 set. (a) Antibodies capable of competing with ACE2 for SARS-CoV-2 binding. (b) Antibodies that bind to the RBD domain of SARS-CoV-2 but do not compete with ACE2. Top: Binding epitope of the RBD in complex with ACE2. Middle: Predicted binding epitope of the RBD in antibody-antigen complexes as determined by tFold-Ag within the SARS-CoV-2 set. Bottom: Experimental structure of the RBD epitope in the antibody-antigen complex with RBD from the PDB.

**Table C20:** CDR loop deisgn accuracy on the SAbDab-22-DesingNano benchmark.

| Method | Extra Features | H1 |  | H2 |  | H3 |  |
| --- | --- | --- | --- | --- | --- | --- | --- |
|  |  | AAR | CAAR | AAR | CAAR | AAR | CAAR |
| tFold-Ag | - | 0.506 | 0.135 | 0.503 | 0.181 | 0.221 | 0.195 |
| tFold-Ag-ppi | epitope | 0.509 | 0.173 | 0.548 | 0.131 | 0.181 | 0.113 |
| tFold-Ag-ppi | epitope and paratope | 0.509 | 0.147 | 0.568 | 0.183 | 0.204 | 0.08 |
| tFold-Ag-contact | contact | 0.462 | 0.173 | 0.490 | 0.117 | 0.188 | 0.132 |

As shown in Table C20, tFold-Ag achieves higher AARs in the CDR-H1 and CDR-H2 regions compared to the CDR-H3 region, which can be attributed to the higher variability of this particular region. In terms of the contact interface, the performance, as indicated by the CAAR scores, remains relatively consistent. However, when we utilized additional prior information (epitope, paratope, contact) as input, although the prediction accuracy of the Nanobody antigen complex improved, we did not observe further performance enhancement in the AAR for Nanobody Design, except on CDR-H2. Upon comparing to antigen-binding antibody design, it becomes evident that the design of antigen-binding nanobodies poses greater challenges (refer to Table B8 and Table C20 for comparison). This could potentially be attributed to the limited availablity of training samples and the extended sequence length of the CDR-H3 region of nanobodies. Nevertheless, current proof-of-concept results suggest that our developed tFold-Ag, the first AI-driven model as far as we know, is appropriate for the design of antigen-binding nanobodies.

#### C.7.2 CDRs loop design using pre-trained ESM-PPI

In our antibody CDRs design experiments, in addition to structure-based methods, we also evaluated the ESM-PPI, the sequence language model component in tFold-Ag, to recover the amino acid of the CDRs. This method does not take antigen information into account, which means that the ESM-PPI method focuses on predicting the CDRs by analyzing patterns and regularities in the antibody sequence itself, rather than relying on interaction information between the antibody and the antigen. Table. B8 and Table. B9 illustrate that, in comparison to the structure-based method, tFold-Ag, the ESM-PPI method, which relies on antibody sequences, presents distinct patterns in the AAR of the CDRs: there is an improvement in CDR-H1, slight alterations in CDR-H2, and a significant reduction in the recovery rate of CDR-H3.

The primary reasons for these observations can be summarized are as follows: (1) V Gene Encoding and Correspondence: Both the CDR1 and CDR2 regions, along with the FR1, FR2, and FR3 regions, are encoded by the V gene, creating a strong correlation between the FR and CDRs [97]. This inherent linkage enables the language model to more effectively capture and utilize such dependency information. (2) Antigen Binding and CDR Diversity: CDR1 and CDR2 are less involved in antigen binding compared to CDR3 [98] [99]. Therefore, incorporating structural or antigen-binding information might not substantially benefit the recovery rates of CDR1 and CDR2. However, the scenario with CDR3 is markedly different. CDR3 is the most diverse region, with its diversity arising from random nucleotide insertions and the introduction of D genes. Apart from a few amino acids at its beginning and end, CDR3 is not encoded by VJ genes. Consequently, its correlation with the framework regions is minimal. Furthermore, CDR3 is the primary region for antigen contact, suggesting that information derived from antigens could be more advantageous for the recovery and design of this region. Our findings highlight the nuanced role of sequence-based models in antibody design. While they are effective in capturing dependencies in certain regions, their limitations become apparent in highly variable and antigen-interactive segments like CDR3. This underscores the need for integrative approaches that combine sequence information with structural and antigenic data to optimize antibody design.

#### C.7.3 Confidence score for antibody design

Evaluating the efficacy of antibody design presents a significant challenge. Ideally, a well-designed antibody would exhibit strong affinity; however, due to the lack of sufficient affinity data, predicting antibody affinity is extremely difficult. It is hard to assess the quality of antibody design without conducting wet lab experiments. This lack of a rapid evaluation method further complicates the task of designing confidence scores. Recent advancements [17, 28] in structure prediction methodologies have introduced confidence measures to assess the quality of predictions. In contrast, most antibody design method [36, 37] employ sequence perplexity (PPL) to establish a correlation between confidence and antibody design.

Given that tFold-Ag is an end-to-end antibody sequence-structure co-design method, we attempt to establish a relationship between the quality of antibody design and the confidence scores of structure prediction. We use the amino acid recovery (AAR) as a metric to evaluate the quality of antibody design. Theoretically, the closer the design is to a natural antibody, the superior the quality of the designed antibody.

In our model, the ipTM score is used to measure the confidence in predicted antibody-antigen interface. Considering the interaction between the antibody’s CDR3 region and the antigen, we evaluate the correlation between ipTM and the AAR of CDR-H3 under three different scenarios. As illustrated in Fig. C13a to Fig. C13c, when not using any Inter-Chain feature, antigen epitope, or PPI features, the Pearson scores for ipTM and AAR are -0.23, -0.07, and 0.10, respectively. This indicates that the confidence score ipTM, used to measure docking accuracy, has no correlation with sequence recovery. That is because AAR is a residue-level metric, while ipTM is a protein-level metric, making it difficult to establish a mapping relationship between the two.

**Fig. C13:**
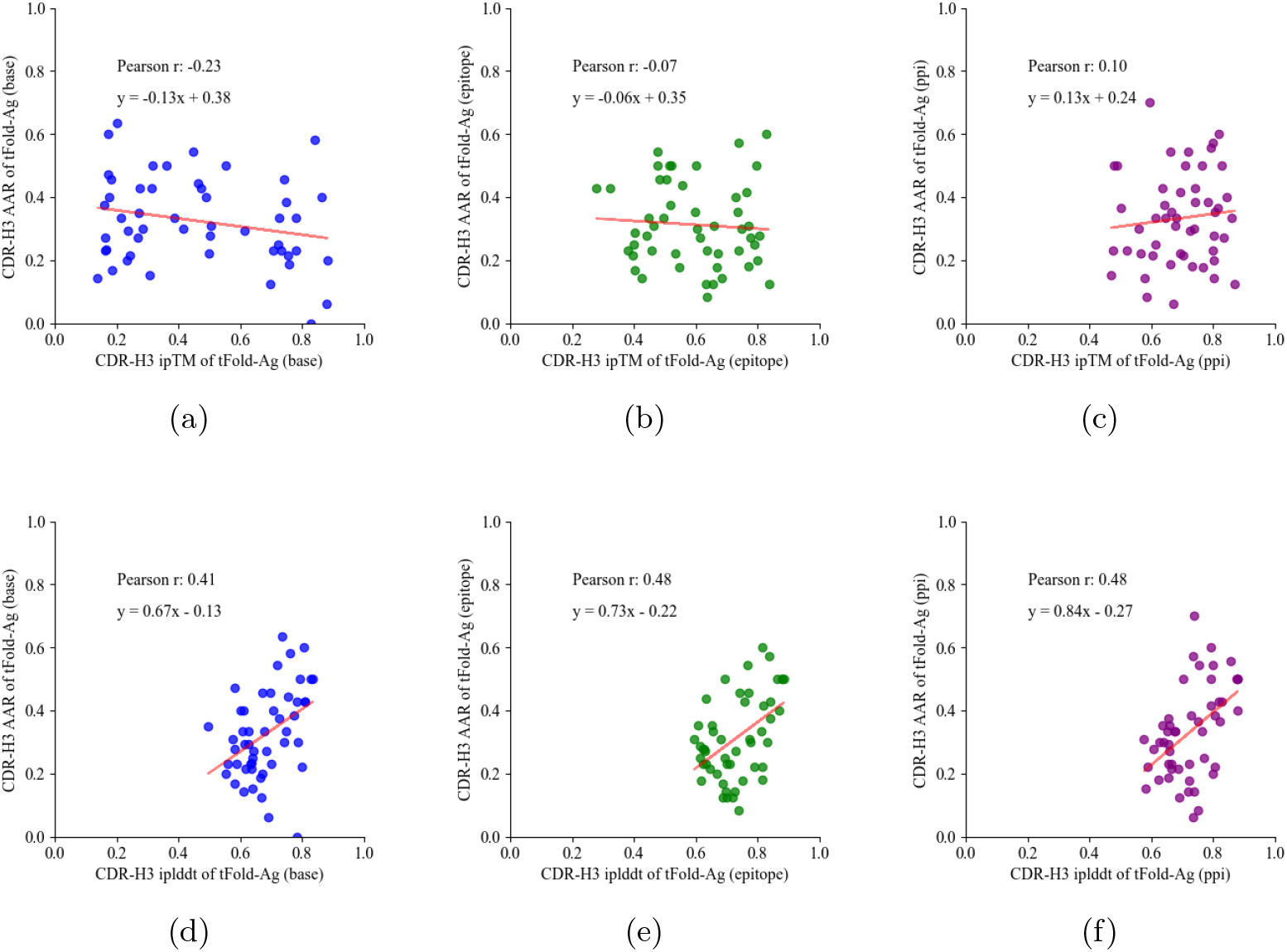
Confidence score analysis for antibody design. (a-c) Correlation between ipTM and AAR of CDR-H3 for various tFold-Ag predictions (base, epitope and ppi) on the SAbDab-22-Design-Ab, with a least-squares linear fit applied to the data. (d-f) Correlation between iplddt and AAR of CDR-H3 for various tFold-Ag predictions (base, epitope and ppi) on the SAbDab-22-Design-Ab, with a least-squares linear fit applied to the data.

Given that the CDR-H3 region are predominantly situated at the interface between the antibody and antigen, we denote the pLDDT of the designed CDR-H3 region as iplddt (interface pLDDT). We explored the correlation between the iplddt of CDR-H3 as predicted by tFold-Ag and the AAR of CDR-H3 designed by tFold-Ag. As depicted in Fig. C13d to Fig. C13f, the Pearson correlation between iplddt and the AAR of different feature is 0.41, 0.48, 0.48, respectively. iplddt reflects the confidence score of the structure of the designed CDR-H3 region as predicted by tFold-Ag. This score has a certain correlation with AAR, especially when the antigen epitope is determined, this correlation is stronger.

This allows us to use the iplddt score as a metric to assess the performance of the antibody CDR-H3 sequences designed by tFold-Ag. We also evaluated the correlation between the iplddt scores for CDR-H1 and CDR-H2 and the AAR of their respective regions. Since CDR-H1 and CDR-H2 are not always part of the antibody’s paratope, this correlation is generally weaker.

## Notes

### Competing Interest Statement

The authors have declared no competing interest.

